# Spatiotemporal Heterogeneity of HPV Integration Architectures Drive Oncogenic Features of Oropharyngeal Cancer

**DOI:** 10.64898/2026.10.03.756294

**Authors:** Wenjin Gu, Apurva Dhanesh Bhangale, Xiaomeng Du, Xinyi Deng, Elizabeth U. Gensterblum-Miller, Jordan Currie, Collin V. Brummel, Vital Nyabashi, Marisa Buchakjian, Keith A Casper, Steven E. Chinn, David Forner, Kelly M Malloy, Michelle L Mierzwa, Mark E Prince, Jennifer L Shah, Andrew G Shuman, Chaz L Stucken, Paul L Swiecicki, Francis P Worden, Pratyusha Yalamanchi, Jonathan B. McHugh, Krystle A. Lang Kuhs, Alan P. Boyle, Hui Jiang, Molly E. Heft Neal, Matthew E Spector, Ryan E. Mills, J. Chad Brenner

## Abstract

Viral integration is common in human papillomavirus-positive oropharyngeal squamous cell carcinoma (HPV+ OPSCC), but its structural and clinical significance remains unclear. We integrated targeted capture sequencing of 197 HPV+ OPSCCs with long-read nanopore sequencing of 17 tumors and models to characterize HPV–host rearrangements. We developed a novel quantitative informatics framework that classifies integration structures as Type1 (simple, single/low-copy) or Type2 (clustered, high-copy, rearranged). HPV integration occurred in 83% of tumors, with recurrent hotspots including *TP63, MYC, CD274, TRAF2,* and *KMT2C*. Type2 events were enriched for HPV *E2* disruption, focal human amplifications, preferential transcription, and noncanonical HPV–host RNA fusion isoforms. Type2 tumors showed increased HPV oncogene expression and activation of MYC, NF-κB/TNFα, PI3K–AKT–mTOR, Hippo/YAP–TAZ, and epithelial stemness programs. Multi-region and longitudinal analyses revealed intratumoral heterogeneity and clonal selection of Type2 architectures during progression, supporting integration structure as a potential marker of aggressive behavior and risk stratification.

## Introduction

Persistent infections with Human Papillomavirus (HPV) contribute to nearly 50,000 new cancers in the United States each year.^1^ HPV is responsible for nearly all cases of uterine cervical cancers and a substantial number of anogenital and head and neck cancers. Notably, the occurrence of HPV+ head and neck cancers, especially Oropharyngeal Squamous Cell Carcinoma (OPSCC), has dramatically risen in the last decade.^2^ Currently, nearly 90% of OPSCCs are associated with HPV, exceeding the incidence of HPV+ cervical cancers in the US.^3,4^

HPV encodes multiple genes, including the well-established oncogenes, *E6* and *E7*, which drive pathogenesis by interfering with cell cycle regulation by disrupting regulators including *TP53* and *RB1*, respectively.^5^ More recently, *E7* has also been shown to degrade *PTPN14*, leading to Hippo pathway activation,^6^ which is also critical for pathogenesis. In contrast, HPV also encodes *E2* which regulates viral transcription, initiates DNA replication, and partitions viral genomes during cell division, but also represses *E6/E7* transcription and is therefore frequently lost during disease pathogenesis.^7^

In addition to introducing oncogenes to the host cell, HPV is also thought to induce additional pathogenetic mechanisms through the process of viral integration.^8–11^ While integration of viral DNA into the host human genome is common in HPV-associated cancers, the mechanistic basis for this process remains poorly understood due to the structural diversity and complexity of integration events. HPV integration events are varied in complexity, with instances of single-copy segment integration, focal amplification in clustered regions with multiple fusions, and diverse patterns described in different studies.^12–15^ HPV integration can be generalized into two types: Type1, involving the integration of a single genome into cellular DNA, and Type2, featuring multiple tandem head-to-tail repeats of the HPV genome at a specific genomic locus.^16^ Subsequent studies, such as Akagi et al. in 2014, introduced a "looping model" to describe focal duplication, with both HPV and human segments forming DNA concatemers around integration events.^17^ A 2022 study focused on cervical cancer revealed additional complexities, including inter-chromosomal translocations and extrachromosomal circular DNA structures.^12^ Further long-read sequencing efforts by Akagi et al. in 2023 subsequently identified similar features in OPSCC.^18^ The 2022 study classified four types of HPV integration (Type A-D) based on different structural variations and their association with *E6* and *E7* genes.^12^ These investigations significantly enhanced our comprehension of HPV integration structures. Nevertheless, these classifications of HPV integration types predominantly relied on descriptive approaches rather than establishing quantitative thresholds that could be used to systematically evaluate integration events. This limitation arises from constraints related to sample size and the technology of resolving complex structural analyses.

In this study, we hypothesized that a comprehensive assessment and rigorous categorization of HPV integration events would provide further insight into tumor progression. We integrated multi-omics datasets encompassing 197 FFPE HPV+ OPSCC patients with targeted capture sequencing (TCS) data, 102 OPSCC samples with RNA-seq transcriptomic data (76 of which were paired with TCS), and an additional 17 samples that also have matched Oxford Nanopore long-read (LR) whole genome sequencing. We introduced a quantitative method for defining two distinct types of HPV integration structures (Type1 and Type2) based on their biological and statistical characteristics at the DNA level, as well as Type0 to indicate a probable episomal state. To accurately portray the complexity of HPV integration, we developed a novel approach to comprehensively resolve local large HPV integration events associated with complex rearrangements at high resolution for LR sequencing data, surpassing the performance of existing LR whole-genome assemblers.

Type2 HPV integrations, characterized by high copy number and clustered integration events, exhibited more aggressive profiles at the transcriptomic level. We then conducted investigations on a subcohort of 54 HNSCC patients with local recurrences, nodal recurrences, or distant metastases, along with two paired cell line models (UPCI:SCC90, derived from a primary base-of-tongue tumor, and UPCI:SCC152, recurrent model at the hypopharynx). Given that HPV integration profiles in large recurrent cohorts were not previously reported, our study has unveiled a series of pioneering findings, elucidating the heterogeneity and clonal selection of HPV integrations during tumor progression. It further addresses a technological gap in resolving local complex HPV integration events from LR sequencing and establishes a standard for further analyses of different types of HPV integrations.

## Results

### Development of an approach to classify the structural complexity of HPV integration events

The structural complexity of HPV integration has complicated efforts to define its biological effects on the host genome. Previous sequencing studies frequently identified multiple HPV host breakpoints within 500-kb genomic regions and classified these events as integration clusters,^12,15,16^ in part because breakpoint proximity was thought to reflect mapping uncertainty or sequencing artifacts. However, advances in long-read sequencing now demonstrate that HPV integration structures can contain genuine complex rearrangements containing multiple distinct HPV-host junctions. Therefore, we hypothesized that breakpoint architecture and local copy number features could be used to infer the complexity of each integration structure.

To begin evaluating this hypothesis, we developed a new inference framework incorporating these two parameters to predict structural complexity (**Figure 1A**) and then explored the relationship among these parameters. We observed that a higher number of integration breakpoints within a cluster correlated with an increased number of duplications within that cluster. Notably, this association did not strictly follow linearity (R^2 = 0.39, p_val = 3.41 × 10⁻¹⁵³) due to some HPV integration events exhibiting exceptionally high copy number (**Figure 1B**). By comparing the distribution of the expected overall copy number at the integrations (green in **Figure 1B**) to the observed overall copy number (blue in **Figure 1B**), we noticed that the observed copy number displayed a steeper slope (***β*** = 9.046) when associated with the number of clustered HPV integrations than expected. This finding suggested that multiple HPV integrations at a clustered site might induce additional host genomic amplifications in the surrounding regions.

**Figure 1.**
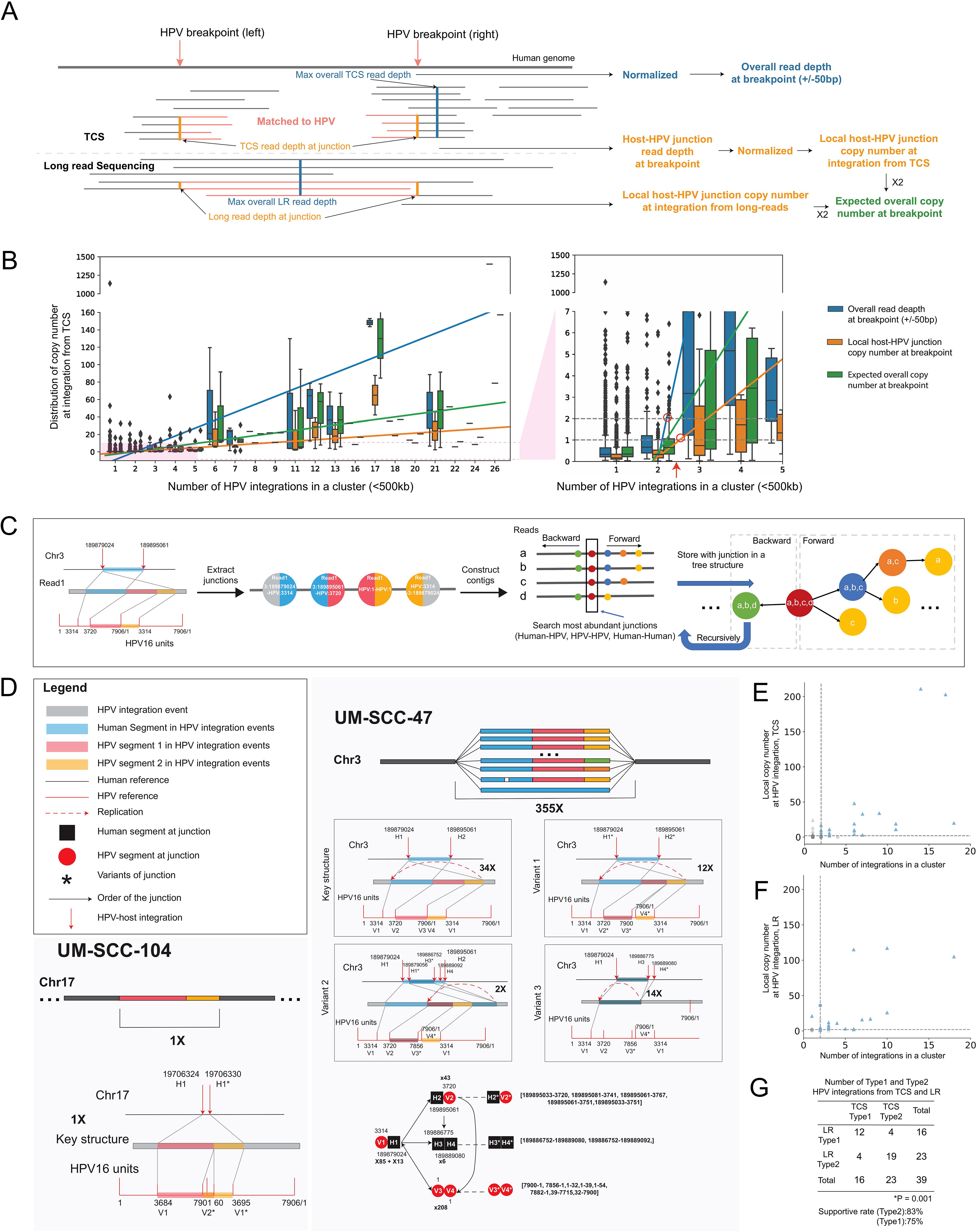
Development and validation of an HPV integration structure prediction algorithm. **A.** Approach for calculating local and overall copy number at HPV integration sites. Split reads at human-HPV breakpoints were counted to calculate the local copy number at the HPV integrations. The maximum read depth within 50-bp of each breakpoint was used to estimate overall copy number. For TCS data, read depths were normalized based on background genes (See method section for details). As a single molecular technology, for nanopore data, the normalization step was not performed. **B.** Statistical summary for three parameters of HPV integration events. X-axis: number of HPV integrations clustered within 500-kb. Y-axis: distribution of copy number at integrations from TCS. The overall copy number and local copy number are in blue and orange, respectively. The expected overall copy number is in green and calculated as twice the local copy number based on the hypothesis that HPV only integrated into one chromatid as a heterozygous event. Linear regression models were fitted as references after removing one outlier suggested by leverage. The pink area in the left panel was zoomed in as the right panel. The two dashed horizontal lines indicate the normal copy number if only one copy of HPV is integrated into the host genome without duplications. The red arrow denotes the intersection of observed copy number and the normal copy number as the number of HPV integrations within a cluster increased. **C.** Schematic plot of novel approach to construct contigs at integration sites. A read from UM-SCC-47 was shown as an example for read storage. The four junctions were called for this read and stored as four nodes. Next, all the informative reads were searched for the most abundant junctions and then stored into a two-direction tree structure. This process was repeated until no reads remained. **D.** Resolved long read structures including UM-SCC-104 as an example of Type1 and UM-SCC-47 as an example of Type2. These HPV integration events were resolved from nanopore sequencing data using the novel approach. Only structures supported by more than three reads are shown. The number of reads supporting the most abundant junction was used as the copy number to summarize the whole structure. For each junction, H denotes human sequence, and V denotes viral sequence and numbers indicate different junctions. If two junctions were within 10-bp, the same number was assigned, however, “*” was added to indicate a variant form. Such cases of variant form junctions were displayed in square brackets. Detailed legends are shown in the legend panel. **E and F.** Type2 HPV integration events identified using TCS and LR sequencing data. Type2 events (shown in blue) were defined as clusters containing more than two integration sites, with a local copy number >1 and a maximum overall copy number >2. Triangles indicate HPV integration events with a maximum overall copy number >2. The plot also shows the concordance of HPV integration-type classifications across different sequencing technologies. **G.** Chi-square test was performed to assess the association between HPV integration types identified by the two sequencing technologies, revealing a significant concordance between the LR and TCS results.

Expanding on this concept, we subsequently sought to determine if a threshold for the number of HPV integrations within a cluster could be used to predict the structural complexity of an HPV-host integration. To test this, we first phased the long reads for a patient-derived xenograft (PDX) model, PDX-294R, using 10X linked-read sequencing data. The result indicated that HPV integration occurred in only one chromatid as a heterozygous event (**Supplemental Figure 4**). Integrations clustered within 500-kb were considered as one HPV integration cluster following previous conventions, and samples without a detected integration were designated Type0 (e.g. episomal, non-ecDNA). If only one copy of HPV is integrated into the host human genome without duplications, which we term a Type1 integration, then the local copy number would be one (**Figure 1B**). If the local copy number at the host-HPV breakpoint was greater than 1, the maximum overall copy number was greater than 2, and the number of HPV integrations in a cluster was greater than 2, we classified them as Type2 HPV integration events.

To validate our approach, we performed matched TCS and LR sequencing on a cohort of 17 samples including 8 HPV+ HNSCC cell lines, 1 HPV+ PDX model and 8 HPV+ HNSCC frozen tissue samples (**Table S1, Supplemental Figure 3**). To our surprise, we discovered that existing long-read genome assemblers, even Telomere-to-Telomere assemblers, failed to accurately assemble HPV integration structures.^19^

This was further borne out by a recent analysis of the HPV16+ UM-SCC-47 cell line, where prior studies have identified clustered HPV-host integration sites on Chromosome 3, near and within *TP63*, accompanied by extensive duplications, pointing towards complex fusion events.^17,20–24^ Using integrated TCS and LR whole genome sequencing in a previous investigation,^24^ we identified six high-confidence HPV-host integration sites by SearcHPV (see **Methods**) and then conducted local assembly to define the HPV16 integration structure in UM-SCC-47.^25^ The resulting 60-kb scaffold comprised a 15-kb, twice-amplified human segment and a 7.5-kb, twice-amplified HPV segment. This was inconsistent with previous 10x linked-read sequencing data, where the integration locus was estimated to span approximately 130-kb in total,^24^ and suggests that current assembly-based approaches may not adequately handle more complex integration structures as seen with HPV. The copy number of the individual HPV integration breakpoints in UM-SCC-47, as defined from our TCS data and WGS reports of others,^17^ varies from 10-50 copies per breakpoint, while the overall copy number at the HPV integration region was uniformly presented as ∼50 copies (**Supplemental Figure 1A**). The variations at each HPV-host integration breakpoint implied that some clustered HPV integration structures have multiple independent sub-structures containing regions of the HPV and human genome that are amplified to varying degrees and adjacent to other HPV and human genome sub-structures to create a larger and more complex superstructure which is difficult for genome assemblers to resolve.

Therefore, to comprehensively resolve complex HPV integration superstructures containing focal genomic sub-structural amplifications, we developed a novel framework based on a ‘junction as nodes’ tree structure called **HPVInsResolver** (see **Methods**). Critically, in our new approach, we preserve the non-canonical junction information from each read and then consider the structure in terms of edges between junctions rather than collapsing them as can occur with de Bruijn graph-based methods with duplicated sequences (**Figure 1C**). As expected, this new method demonstrated superior performance compared to de Bruijn graph-based long-read assemblers as it was the only approach capable of elucidating the local sequence structure within the large duplication regions of the UM-SCC-47 integration superstructure. We recently published the performance of existing assemblers on these genomic data.^19^ For integration structures with relatively low copy number, traditional methods produced concordant structural reconstructions. However, our method was able to capture additional complex configurations, including repetitive HPV–HPV junctions, tandem viral duplications, and looped structures that were incompletely represented or collapsed in the traditional assemblies.

To more rigorously assess the accuracy of our approach, we first examined two distinct classes of HPV integration structures: UM-SCC-47 (Type2) and UM-SCC-104 (Type1). In UM-SCC-47 (**Figure 1D**), the predominant repetitive structure was amplified 34-fold and consisted of a single human genomic segment flanked by two HPV segments joined across the viral reference genome origin. The human-HPV junctions flanking this structure were “*E2* to *TP63* intron10” and “*TP63* exon14 to *E2*”. The two HPV segments spanned nearly the entire HPV genome, with a 406-bp gap at the *E2* gene. Variant 1 of the key structure, with 12 copies, exhibited slight offsets at the human-HPV and HPV-HPV junctions at the HPV origin. Another structure, variant 2, encompassed variant 1 followed by an additional human segment from *TP63* intron10 to intron11, partially amplified twice from the two HPV segments to the second human segment. Variant 3 initiated from a human-HPV junction, “*E2* to *TP63* intron10,” connecting to a human segment, followed by a human-human junction indicating a deletion. The human segment extended from *TP63* intron10 to intron11 and was present in 14 copies. The mapping plot delineates the connections among these junctions within this intricate duplication of HPV integration events, highlighting intratumoral heterogeneity within this event. A portion of this structure was previously captured in an assembly generated by wtdbg2, indicating amplification twice.^26^ Of note, a previous study attempted to elucidate the same structure through whole-genome sequencing (WGS) and employed targeted PCR, sanger sequencing, and chromosome walking to establish connectivities.^15^ In their map, they identified the key structure amplified 20 times and variant 3 amplified twice.

Accordingly, in the UM-SCC-104 cell line model, an entirely distinct structure revealed only one copy of the integrated HPV segments within the host gene *SLC47A2* intron 9. The integrated HPV broke at the *E2* region, covering nearly the entire HPV genome with an 11-bp gap and a 65-bp overlap at the origin of the HPV reference (**Figure 1D**). This structure, which did not have the long repetitive sub-structures that impaired the accuracy of existing assemblers, was corroborated by TCS data and nanopore assembly from wtdbg2 (**Supplemental Figure 2**). By resolving these two representative models, our novel approach demonstrated a higher resolution and more comprehensive performance compared to existing methods, and the establishment of this method facilitated further confirmation of HPV integration events with complex structural variations. As an additional orthogonal validation of complex structure resolution, we used the approach to solve the HPV integration structure in UD-SCC-2 and then performed PCR with primers on either side of multiple HPV-human breakpoints, a representative example of a complex structure with multiple breakpoints clustered together on the X chromosome. The resulting PCR product lengths support the predicted breakpoints and fragment orientations (**Supplemental Figure 1B**).

After validating our framework for solving HPV integration structures from long-read data, we were then able to test our original hypothesis and the accuracy of the threshold for the short-read based HPV integration structural complexity prediction. Due to distinct coverage between these two sequencing technologies, TCS exhibited higher sensitivity than LR, as previously validated by PCR sequencing in UM-SCC-47 and PDX-294R.^24^ Supportive rates were calculated based on common HPV integration events (see the method section) shared by these two sequencing technologies (**Figures 1E and 1F**). The supportive rate from LR to TCS for Type2 events was 83%, while for Type1 events, it was 75%. The definitions of Type1 and Type2 events from LR and TCS demonstrated significant associations (p_val < 0.01; **Figure 1G**), showing that this threshold could be used to accurately infer the structure of an HPV integration cluster using TCS data alone.

### Defining relationships of integration status and structure with outcome

Having established an approach to use short-read TCS data to infer integration structure complexity, we then sought to study the relationship of HPV integration structures with clinical outcome in a larger HPV+ OPSCC cohort. We performed TCS on a cohort of 197 HPV+ OPSCCs. The majority were men (81%) and white (89%) with an average age of 58 years (39-90). Forty-four percent had never smoked, 13% had ≤10 pack-year smoking history, and 39% with >10 pack-years history. Early T stage (T0-T2) made up 66% (130/197) of the cohort and 34% (67/197) with advanced stage (T3-4). The majority were node-positive (87% - 172/197) with most of the node-positive patients having N1 disease (74.6%) (**Table S2**).

HPV genotyping using HPViewer^27^ revealed that all tumor samples analyzed were HPV-positive. Among the 197 tumors assessed, 183 (92.8%) harbored HPV16 as the sole detectable viral type. A subset of eleven tumors (5.6%) exhibited co-infection with HPV33, while two tumors (1.02%) showed co-infection with HPV18, and one tumor (0.5%) contained both HPV16 and HPV35. No other HPV genotypes were detected. There was no significant difference in median five-year disease-free survival between patients with HPV16 mono-infection and those with infections with HPV18 or HPV33 (74.9% vs. 55.6%, p_val > 0.5) (**Supplemental Figure 5**).

Among the 183 HPV16+ OPSCC tumors analyzed (excluding tumors with co-infection), viral integration was detected in 155 cases (85%). In total, 3,824 HPV integration events were identified. Integration events into host genes were identified in 149 tumors (81.4%), with affected tumors harboring a median of 75 (range: 0–275) host genes containing integration sites. The number of host genes with integration, analyzed both as a continuous variable and dichotomized based on the median, was not significantly associated with disease-free survival in Type1 and Type2 tumors, although a non-significant trend toward improved outcomes was observed in tumors with fewer integration events (**Supplemental Figure 6A**). Of the 149 integrated tumors, 55 (35.5%) showed integration within established cancer genes included on COSMIC Cancer Gene Census Tier 1 or 2 lists (**Supplemental Figure 6B, Table S2**). The most frequently targeted loci were *CD274* (PD-L1; n=9), *TP63* (n=8), and *KMT2C* (n=6). Additional recurrently affected genes included *CSMD3* (n=4), as well as *PTPRD*, *ROBO2*, *LPP*, *MSI2*, *ESR1*, *CTNNA2*, and *LRP1B* (n=3 each). Less frequent, yet notable, recurrent integration events were also identified in *ARID1B*, *RUNX1*, *NFIB*, and *RAD51B, FAS, CDK6, ERBB4,* among others (n=2 each).

Pathway enrichment analysis of the host genes harboring viral integrations revealed a strong convergence on oncogenic networks central to HPV+ OPSCC biology, supporting a clear pathogenic role of direct disruption of host genes by integration. The most significantly overrepresented pathways (**Supplemental Figure 7**) included MYC signaling (FDR < 0.005), NF-κB and TNFα inflammatory signaling (FDR < 0.005), and PI3K-AKT-mTOR activation (FDR < 0.05), all canonical drivers of proliferation, metabolic adaptation, and survival in HPV+ OPSCC. Integrations also mapped near regulators of epithelial stemness and differentiation, including the TP63 (FDR<0.005), Hippo/YAP-TAZ (FDR < 0.005), Hallmark UV Response Down (FDR < 0.0002) and WNT/β-catenin pathways (FDR < 0.005), supporting a role in sustaining a basal, progenitor-like state characteristic of aggressive squamous tumors. Additional integrations into chromatin remodeling (*KMT2C*, *NSD3*, *ARID1B*) and DNA damage repair (*RAD51B*, *CHEK1*, *POLQ*) genes suggest that viral integration may further destabilize the genome and reinforce transcriptional reprogramming. Collectively, these findings indicate that HPV integration preferentially targets genomic regions controlling cell-cycle regulation, inflammatory signaling, and epithelial lineage plasticity, thereby regulating pathways that are key drivers of HNSCC initiation, progression, and therapeutic resistance. Critically, this supports that molecular analysis of HPV+ tumors must include HPV integration analysis as it is a core mechanism disrupting the host genome.

### Characteristics of Type1 and Type2 HPV integrations

We then applied our structure prediction algorithm to the tumor cohort. In total, 4,287 HPV integration events were identified across 214 samples, including 3,975 integrations from 197 HPV+ samples in the core cohort (of which 3,824 were from 183 HPV16+ samples) and 312 integrations from 17 cell line and tissue samples. Of all identified integration events, 3,601 (84%) were classified as Type 1 and 686 (16%) as Type 2. Among the Type 2 events, 552 were identified in the core cohort and 134 in the cell line and tissue cohort. Approximately 39% (83/214) of samples contained at least one Type2 HPV integration and were defined as Type2 samples; the rest of the samples with HPV integration detected were defined as Type1 (**Figures 2A & 2B**). Type2 integrations were identified across the entire HPV genome and from all autosomes and chromosome X. Several hotspots were observed for Type2 integrations, including *TP63*, *MYC*, *LINC00484*, *TRAF2*, *DTX4*, and *PTCSC2*. Of note, several of these genes are highly expressed in keratinocytes or deregulated in HNSCC including *TP63* and *MYC.*^28–32^ Notably, multiple Type2 HPV integrations with high copy number were located on chromosomes 8 and 9 (**Figure 2B**). The distribution of breakpoints across the HPV genome revealed distinct patterns between the two integration types. Type2 breakpoints were significantly enriched within the *E1* and *E2* regions relative to a uniform distribution across the HPV genome, while significantly fewer breakpoints occurred within *E6*, the *URR*, *L1*, and *L2* regions. In contrast, Type1 integration events showed significant enrichment in the *E1* and *E5* regions and depletion in the *E2* and *E7* regions, in addition to the breakpoint patterns observed for Type2 events (**Figures 2C & 2D**).

**Figure 2.**
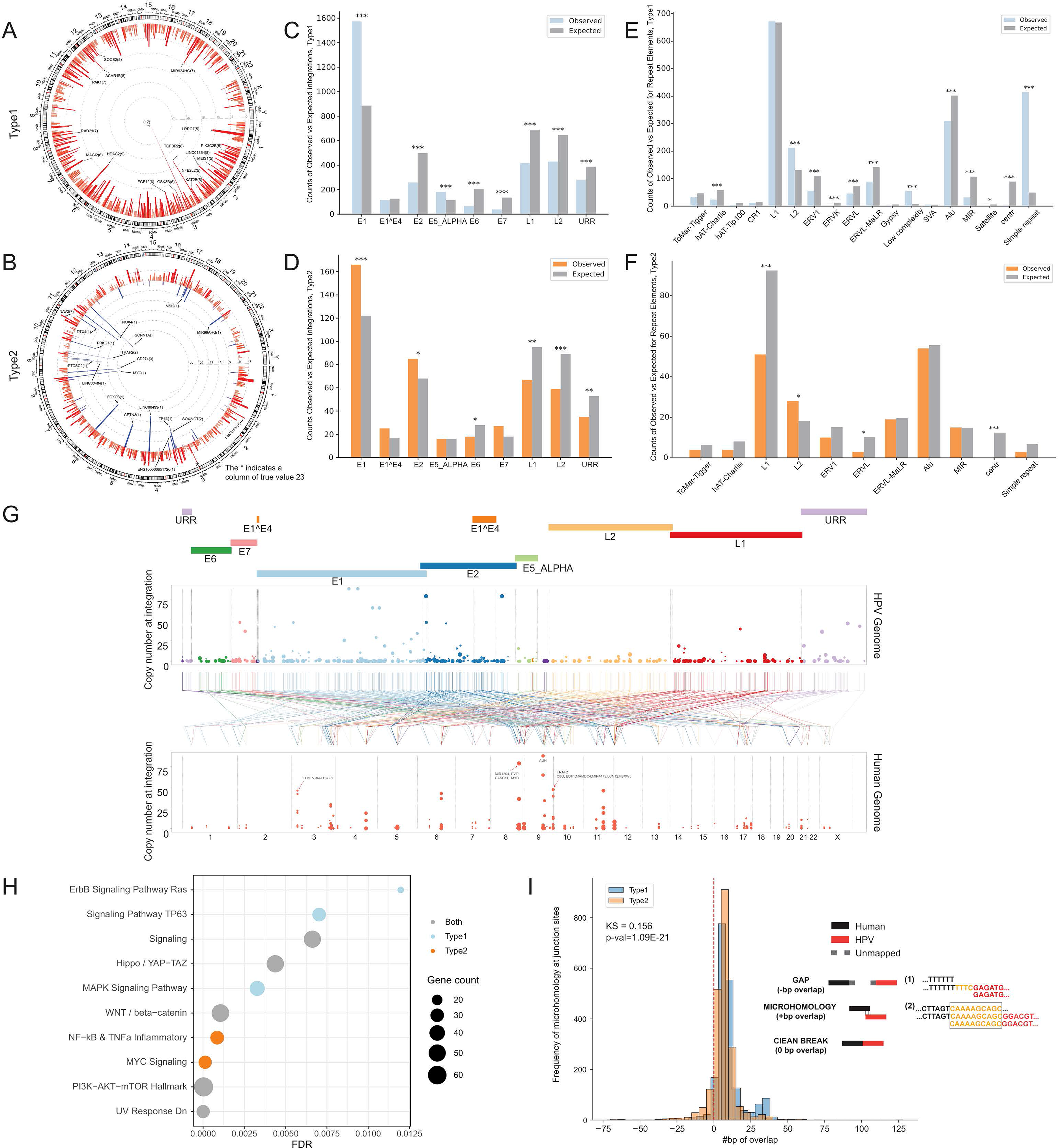
HPV16 integration events converge to disrupt key functional pathways in HPV+ OPSCC. **A-B.** Distribution of Type1 and Type2 samples’ HPV integration on the human genome. Red bars indicate the number of integrations marked by a dashed line within a genome window. Hotspot genes close to the main breakpoints were annotated. Numbers in the brackets indicate the corresponding sample counts within each window. **C-D.** Distribution of Type1 and Type2 HPV integrations on HPV genome. Expected values were based on the assumption that HPV integrations were randomly distributed on HPV genome. Binomial tests were performed. 1, 2, and 3 “*” indicated the p_val thresholds of <0.05, <0.01 and <0.001 correspondingly. **E-F.** Distribution of repeat elements enriched in Type1 and Type2 HPV integration. Expected values were based on the assumption that HPV integrations were randomly distributed on the human genome with the blackout regions excluded. Definition of repeated elements is based on UCSC repeat mask dataset. Binomial tests were performed. 1, 2, and 3 “*” indicated the p_val thresholds of <0.05, <0.01 and <0.001 correspondingly. **G.** HPV breakpoints for Type2 events on the human genome and HPV genome. Bottom panel: Type2 HPV integration events in human genome; Top panel: Type2 integration events in HPV genome. Breakpoints on the human and HPV genome were linked and colored by the HPV gene the breakpoints fell into. The size of the dot denoted the number of integrations in this event. Note that since HPV integration events were defined based on location of breakpoints on the human genome, one HPV integration event might have 1-2 breakpoints captured on the HPV genome. Hotspot genes were marked. **H.** Frequency of size of microhomology at Type1 and Type2 junctions. Microhomology was measured using the number of base pairs overlapping at the host-HPV junctions. Positive values indicated microhomology, negative values indicated gaps and zero indicated a clean break. Kolmogorov–Smirnov test was performed to compare Type1 and Type2 junctions. **I.** Gene set enrichment analysis of host gene pathways disrupted by HPV integration. Results are shown as disrupted by Type1 only, Type2 only, or both integration structure types.

We further investigated the enrichment of HPV integration sites within repetitive elements of the human genome across Type1 and Type2 integration events. Compared to an expected model of uniform distribution, we see less than expected integration events on several LTR retrotransposon elements and Alu elements in both Type1 and Type2 samples, as well as a much higher number of events in L2-LINE elements and simple repeats (**Figures 2E & 2F**). We focused on the junction area to capture potential microhomology by calculating the size of read overlaps that mapped to both human genome and HPV genome. We observed overall positive overlaps in both Type1 and Type2 events with a significant difference in the distribution of overlap sizes between the two integration types (p ≤ 1.09 × 10⁻²¹; **Figure 2H**). Furthermore, the pathway enrichment of integration genes between Type1 and Type2 have both differences and similarities. In Type1, the integration genes were enriched for ErbB signaling pathway, Ras signaling pathway and MAPK signaling pathway; Type2 tumors showed enrichment for NF−kB & TNFa inflammatory and MYC signaling. Several pathways were commonly enriched in both subtypes, including PI3K–AKT–mTOR, Hallmark UV response down, WNT/β-catenin, Hippo/YAP–TAZ, and TP63 signaling (**Figure 2I**).

Importantly, our standard TCS panel also contained capture probes not only for HPV, but also 225 host genes commonly altered in HPV+ OPSCC tumors providing the opportunity to test for relationships between HPV integration events and structures with common host genome alterations found in this disease.^33,34^ Focusing on all HPV16+ tumors, the average number of mutations identified per tumor was 2.67 (range 0-50) across the 225 genes tested, and gene-specific alteration frequencies were consistent with previous reports^35,36^ (**Supplemental Figure 8, 9**). The total number of alterations detected in a tumor was not significantly correlated with the number of HPV16 integration sites, the number of host genes disrupted by integration, or the type of integration structure. We therefore grouped the tumor alterations by molecular pathway and, for each pathway, stratified tumors according to the presence or absence of mutations in pathway-associated genes. The pathways analyzed were RTK/PI3K/RAS, NOTCH, DNA damage, epithelial differentiation, immune signaling, cell cycle, cell death, TP53/MDM2, chromatin modification, and oxidation. We then tested for associations between pathway-level mutation status and HPV integration status. After false discovery rate (FDR) correction, only the TP53/MDM2 pathway showed significant enrichment (p_val = 0.0086), with pathway mutations occurring more frequently in Type0 tumors (25%) than in tumors with integrated HPV (3.9%). No other pathways differed significantly by integration status (**Supplemental Figure 10**).

We then performed Kaplan-Meier survival analysis for the cohort of 183 HPV16+ OPSCC patients to determine the effect of clinicopathologic variables, integration status, and predetermined genomic events on survival outcomes. Overall, there were 27 recurrent or metastatic tumors and 156 primary tumors. We first looked at survival outcomes by primary versus recurrent disease and as anticipated, the recurrent/metastatic cohort did significantly worse (five-year disease-free survival (DFS), defined as death from any cause or disease recurrence: 74.4% (95% CI: 66.3-80.8%) versus 18.5% (95% CI: 6.75-34.8%), p_val <0.0001, (**Supplemental Figure 11A**). We therefore performed the remaining survival analysis on the subgroup of primary tumors. On univariate analysis, tumor stage, HPV16 integration type were significantly associated with disease-free survival. For T stage, early stage (T0-2) had improved survival compared to advanced stage disease (T3-4), 80.5% (95% CI: 71.0-87.2%) versus 60.9% (95% CI: 45.2-73.3%), p_val < 0.005. For integration status, Type1 and Type2 integration had similar five-year DFS 76.1% (95% CI: 66.8-81.2) versus 82.5% (95% CI: 65.2-84.1), p_val > 0.5. Type0 (episomal) status had the worst five-year DFS 51.9% (95% CI: 29.1-70.6%) and this was significantly worse than tumors with either Type1 or Type2 integrations (p_val < 0.02). Interestingly, patients with integration affecting more than five host genes had better disease-free survival than patients with five or fewer integrated genes (p = 0.05) (**Supplemental Figure 11B-E**).

Next, we performed additional univariate survival analysis for tumors segmented as having a host gene mutation in a pathway (e.g. oxidation pathway) or not. This analysis showed that tumors with host mutations in the epithelial differentiation pathway had significantly worse five-year DFS compared to patients without an alteration 52.7% (95% CI: 23.4-75.5%) versus 76.3% (95% CI: 67.9-82.8%), p_val < 0.01 (**Supplemental Figure 11F**). As previously noted, HPV integrates into key genes from several of these pathways. Therefore, we tested for the impact of convergent host pathway disruption by either host gene mutation or HPV16 integration. We evaluated whether HPV16 fusion-mediated disruption of key pathways and gene sets, including MAPK, RAS, and ERBB signaling; genes downregulated in response to ultraviolet radiation; PI3K–AKT–mTOR, WNT/β-catenin, TNF-α/NF-κB, Hippo, and noncanonical NF-κB signaling; and MYC and TP63γ target genes, was associated with survival outcomes. No significant association was observed between pathway disruption and survival (**Table S3)**.

On multivariate analysis using cox regression, tumor stage, epithelial differentiation pathway alterations, and integration status (episomal/Type0 vs. integrated/Type1 or Type2) were independent predictors of disease-free survival. **Supplemental Figure 11G** demonstrates hazard ratios and 95% confidence intervals for each of these variables.

We were further interested in looking at clinicopathologic and genomic variables associated with integration status. Chi-square analysis was performed to evaluate differences between Type0 and Type1 or Type2 integration cohorts (**Table S4**). We also evaluated the total number of HPV16 integration events as an independent variable. The total number of integrations was not significantly associated with disease-free survival (p_val > 0.5, **Supplemental Figure 12).**

### The impact of HPV integration structure on the tumor transcriptome

To then understand the impact of HPV integration structures on the transcriptome, we performed bulk RNA sequencing on 102 HPV+ OPSCC samples (96/102 (94%) were HPV16+). With this cohort, we first annotated HPV-host gene fusions using SurVirus and identified 162 fusions in 35 samples (**Table S5**). In previously published work, the presence of an HPV-host fusion gene has been associated with poor outcome in HPV16+ OPSCC.^37^ Importantly, here we were able to independently validate that patients with at least one HPV-host fusion had a significantly worse 5-year disease-free survival (DFS) of only 60% (95% CI 35.7-77.6%), while patients without an HPV-host fusion detected had a 5-year DFS of 82.3% (95% CI 66.4-91.1%), p_val < 0.05 (**Figure 3A**).

**Figure 3.**
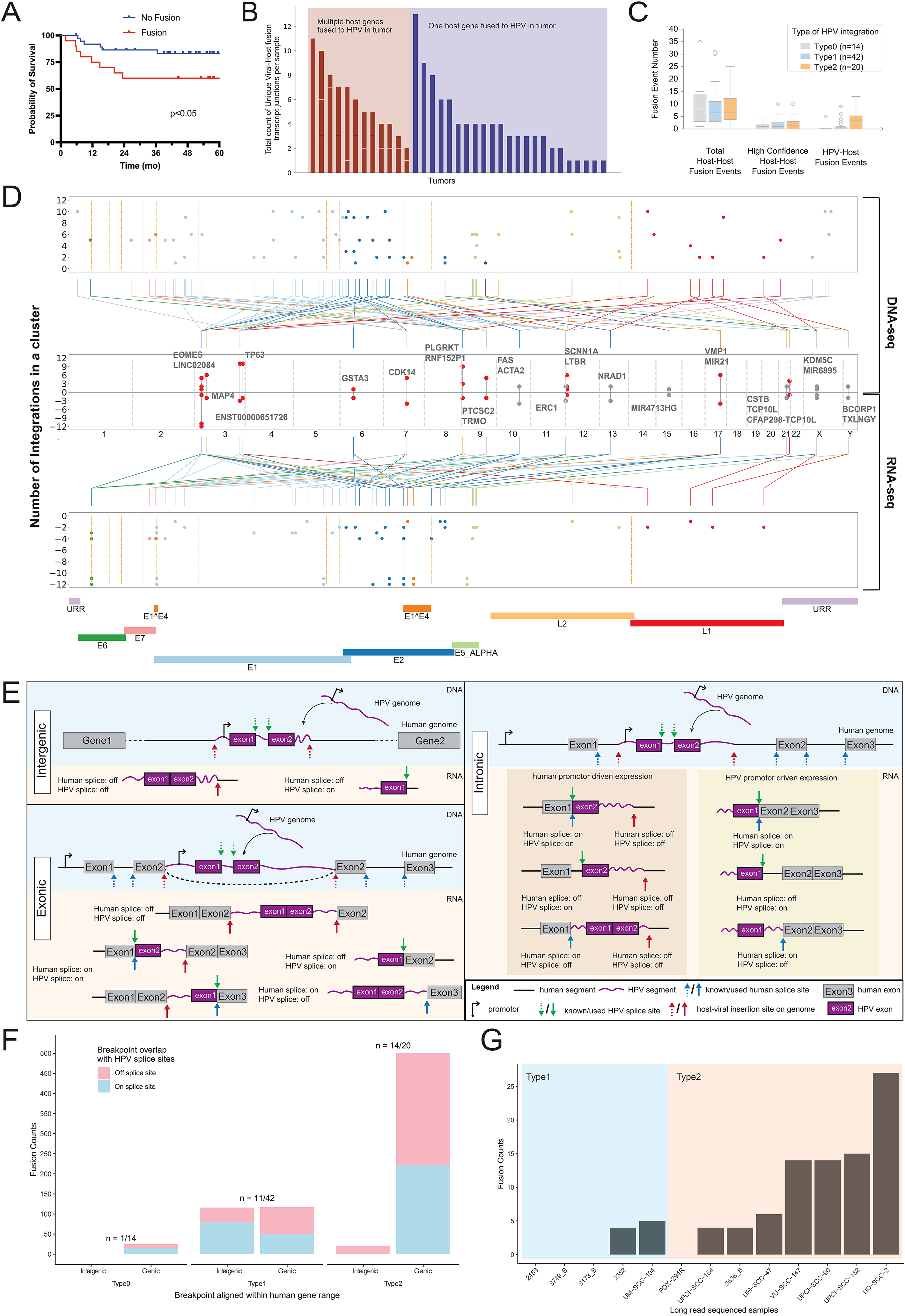
HPV integration structural complexity drives increased structural complexity of HPV-host fusion transcripts in HPV+ OPSCC. **A.** Five-year disease-free survival (DFS) by HPV-host fusion status. **B.** HPV-host fusion site counted per sample. The red section on the left are samples with multiple genes fused with HPV in the transcript; each bar is stacked by multiple gene groups of that sample. Fusion site locations are grouped into one gene range if they are within 50-kb. The blue section on the right are samples with only one fused gene. **C.** Comparison of HPV fusion and host fusion events across HPV integration types. The numbers of host–host and HPV–host fusion events were compared among the three HPV integration types. **D.** Common HPV-host fusions between TCS and RNA-seq. Top panel: HPV breakpoints on HPV genome of TCS; Bottom panel: HPV breakpoints on HPV genome of RNA-seq. Each breakpoint was presented as a dot colored by HPV genes. Middle panel: common HPV fusions on human genome from TCS and RNA-seq. Red: Type2; Grey: Type1. Each paired HPV fusion event was connected. Y-axis indicated the number of HPV integrations in the event (clustered within 500-kb). Breakpoints on the human and HPV genome were linked and colored by HPV genes. Orange dashed lines on top and bottom panel were canonical splicing sites of HPV transcripts (https://pave.niaid.nih.gov/ (VanDoorslaer et al. 2017)). **E.** Scheme of how splicing events may affect fusion transcript formation. HPV genome may integrate into the human genome in the intergenic region or intronic/exonic regions. HPV and human known canonical splice sites (green and blue arrows, respectively) may be leveraged to form shifted breakpoint location in the RNA compared to original integration site in the DNA (red arrows), and noncanonical splice sites may be used for splicing paired with canonical splice sites. With intergenic integrations, HPV promoters drive the transcription, and intergenic DNA segments may be expressed. With intronic/exonic integrations, transcription may start with HPV or human promoters, and human gene exons, full or truncated, may be expressed in the fusion transcripts. Not all possible scenarios are denoted here. Models of chimeric transcripts. Chimeric transcripts were generated by HPV integration events that were transcribed. If HPV integrated into the intron of the gene, after splicing, the original breakpoints at the DNA level were lost. We observed HPV fusions the same as the splicing sites. If HPV integrated into exon regions, chimeric transcripts might preserve the fusions at DNA level. The same logic also applied to HPV integrated parts and resulted in three different patterns of human-HPV fusions in transcriptomic level: human gene splicing site-HPV gene splicing site; human DNA breakpoint-HPV gene splicing site; human DNA breakpoint-HPV DNA breakpoint. **F.** RNA fusion site counts breakdown by sample integration subtypes, overlapping with human gene range and known HPV splicing sites. “n=Samples with RNA fusion detected/total count of subtype samples” is annotated above each integration type. The y-axis is the count of RNA fusions detected. The x-axis is grouped first by sample integration type and then by whether the RNA fusion site overlaps a human gene region (defined as ±50 kb). Overlap with known HPV splice sites was defined using a 2-bp window. To assess whether RNA fusion sites were significantly associated with (a) human gene regions or (b) known HPV splice sites, we performed chi-square tests. The association was significant for both Type 1 and Type 2 samples (p < 0.0002) and remained significant when considering all samples (p < 0.01). **G.** RNA fusion counts for 13 cell line samples with integration types.

After confirming the association between HPV–host fusions and clinical outcome, we then sought to understand the potential mechanistic role of these molecular events. Within each sample, we grouped the detected fusions by their annotated location on the human genome with a 50-kb distance range, then merged the clusters within 50-kb to form each final gene cluster. Notably, while 23 tumors harbored only a single HPV-host fusion gene cluster, we observed that 12 tumors harbored more than one fusion gene cluster (with a median of 2, mean of 2.1 and ranging from 2 to 3) (**Figure 3B**). Interestingly, ssGSEA analysis of the host fusion partners involved in HPV-host fusions demonstrated an enrichment of several key squamous-epithelial oncogenes and tumor suppressors (*TP63*, *TRAF3*, *RELB*, *WWOX*, *SOCS1*) that converged on potentially critical pathways including non-canonical NF-κB signaling and epithelial stemness pathways (**Supplemental Figure 15**). Some of the remaining coding gene fusion partners including *CDK14*, *MAP4*, *PRCC*, *NRG2* are also known to contribute oncogenic phenotypes including cell-cycle, cytoskeletal, and signaling dysregulation in various contexts, but did not converge on common pathways. Likewise, a few non-coding entries (e.g., *MIR21*, *STXBP5-AS1*, *LINC02084*) were also identified as the host fusion partner and represent potential cis-regulatory partners for the associated portion of the HPV transcript contained in the fusion.

Given the complexity of HPV-host fusion transcripts and association with DFS, we then sought to test whether host-host fusion transcripts also had a significant role in HPV16+ OPSCC survival and identified 124 high confidence host-host fusions in 76/102 tumors (Average = 1.6 total fusion calls per tumor, STDEV=2.2, **Figure 3C**). Tumors with HPV– host fusion had approximately 2.5-fold higher odds of harboring high-confidence host–host fusion than tumors without HPV–host fusion (73.1% versus 52.0%; odds ratio = 2.51). However, this association did not reach statistical significance (two-sided Fisher’s exact test, p_val > 0.05). (**Supplemental Figure 13**). Likewise, no significant difference in survival by host-host fusion status was observed 72.4% (95% CI: 50.5-85.8%) versus 77.2% (95% CI: 58.1-88.5%), p_val > 0.5) (**Supplemental Figure 13A**).

From this analysis, we noted the presence of several interesting host-host fusion transcripts, including one case (5343-CB-5) with the *FGFR3*-*TACC3* fusion recently found to drive oncogenic phenotypes in a small subset of HPV+ OPSCCs.^38,39^ Other oncogenic fusions found in the cohort, include *MAPK9*-*DNMT1*, *MAPK9*-*RASGEF1C*, *NBEAL1*-*ERBB4*, *ALDH1L1*-*ERBB4*, *MET*-*KAT2B*, and *C3orf52*-*TIGIT* and the respective fusion structures are shown in (**Supplemental Figure 14**). Given that we did not identify highly recurrent host-host fusion transcripts, we then performed ssGSEA pathway enrichment analysis to test enrichment of pathways from genes involved in the high confidence host-host fusions. In contrast to the HPV-host fusion analysis, we did not observe any significant pathway enrichment. This demonstrates that host-host fusions are non-convergent in HPV+ OPSCC and further supports the importance of HPV-host fusion formation for disease pathogenesis.

Of the 96 HPV16+ OPSCCs with transcriptome data, 76 had paired TCS DNA data enabling an integrated analysis on these tumors (**Table S6)**. 14 samples harbored exclusively episomal DNA (Type0), 42 samples harbored only Type1 HPV structures and 20 harbored at least one Type2 HPV integration enabling integrative analysis. Importantly, HPV–host fusion transcripts were detected in 1 of 14 (7%) Type 0 tumors, 11 of 42 (26%) Type 1 tumors, and 14 of 20 (70%) Type 2 tumors, indicating significant enrichment of tumors harboring at least one fusion transcript among those with Type 2 structures (chi-square test, p_val < 0.01). This represented 122 total HPV-host fusion genes in the 76 tumors (5 fusions in Type0, 45 in Type1, and 72 in Type2).

We then used the dataset to gain deeper insights into the molecular mechanism of HPV-host fusion formation given the discovery that many of the HPV-host RNA structures that we identified contained more than one HPV-host junction (**Figure 3B**). We observed significantly more RNA fusion events in Type2 samples than Type1 or Type0 (**Figure 3C**). Among this DNA-RNA paired data, 40 pairs of HPV breakpoints were found to be identical across DNA and RNA levels (**Figure 3D**). Most integration breakpoints occurred within the HPV genes, *E1* and *E2*, consistent with known HPV integration hotspots that disrupt the viral replication genes, leading to loss of episomal maintenance and constitutive *E6*/*E7* oncogene expression. Secondary sites (*L2, E1^E4,* and *E5α*) occurred less frequently, and rare breakpoints in *URR* and *E7* suggest occasional regulatory or promoter disruptions in the host (**Figure 3D**). Another pattern emerged when comparing the breakpoints on the HPV genome between the DNA and RNA levels. We noted that multiple HPV fusion transcript breakpoints (bottom panel, **Figure 3D**) aligned with canonical HPV splicing sites (orange lines), a phenomenon not observed in DNA integration breakpoints, suggesting that normal splicing machinery is commonly used in HPV-host fusion formation. This is observed in both Type1 and Type2 samples and quantified in **Figure 3F**. We also noticed that some DNA breakpoints are preserved in RNA fusion on the HPV genome and human genome, though shifting of the breakpoint is commonly observed. Based on these findings, we propose a chimeric transcript model (**Figure 3E**) to illustrate how different RNA fusion mechanisms correspond to underlying HPV integration events. When HPV integrates into the intron of a human gene, the viral–host breakpoints may be removed during RNA splicing. In contrast, if integration occurs within an exon and provided that transcription is not disrupted, the breakpoint can be retained in the mature fusion transcript. When the HPV integrates into an intergenic region of the human genome, transcription is most likely driven by HPV promoters, and the HPV splicing sites are probably utilized. For the HPV genome, if the entire genome is inserted, HPV might be transcribed as canonical isoforms, generating additional fusion genes only present at RNA level. However, when only part of the HPV genome integrates, or when the integration occurs at non-canonical sites, breakpoints may also be present and consistent at both the DNA and RNA levels.

We then examined the number of integrations of Type1 and Type2 events that aligned with canonical splicing sites of HPV genes or non-canonical breakpoints on the HPV genome, grouped by genic and intergenic alignment in the human genome. Surprisingly, we observed that the majority of RNA fusion breakpoints are genic for Type2 samples, whereas the intergenic and genic breakpoints are quite equal in Type1 samples (**Figure 3F**). We also observed that intergenic RNA breakpoints are significantly more likely to be on canonical HPV splice sites than off compared to genic, within both Type1 samples (Chi-square p_val < 0.05, **Figure 3F**). Interestingly, Type2 events presented significantly more non-canonical breakpoints in the HPV-host fusion transcripts, while Type1 aligned more with canonical splicing sites (p_val <= 0.01). As a control for this test, we then compared the number of host-to-host fusion events by integration type, which demonstrated no significant differences (**Figure 3C**). This supported that the complexity of HPV-host fusions is related to the complexity of HPV-host DNA integrations, and was not due to disruption of splicing machinery that would more broadly impact host-host fusion formation. Overall, Type2 tumors in the 76-tumor cohort were more likely to express HPV–host fusion transcripts than Type1 or Type0 tumors (**Figure 3F**). This pattern was also observed in the cohort of 13 cell-line and tissue samples with available RNA-sequencing data, in which Type2 samples exhibited substantially more HPV–host fusion transcripts than Type1 samples (**Figure 3G**).

Expanding on the impact of integration structure on HPV-host fusion gene complexity, we then sought to test if the inferred structure of an integration also related to the expression of HPV genes in the tumor. Thus, we calculated the expression of viral transcripts for *E6, E7, E1, E1^E4, E2, E5, L2* and *L1* and compared the distribution between samples with different types of HPV integration **(Figure 4B)**, which showed a significantly higher expression of HPV oncogenes *E6* and *E7* transcripts in Type2 tumors (Kruskal-Wallis Test: p_val < 0.05). This demonstrated that the structure of HPV integration not only influences HPV-host fusion gene formation, but also associates with high expression of HPV genes.

**Figure 4.**
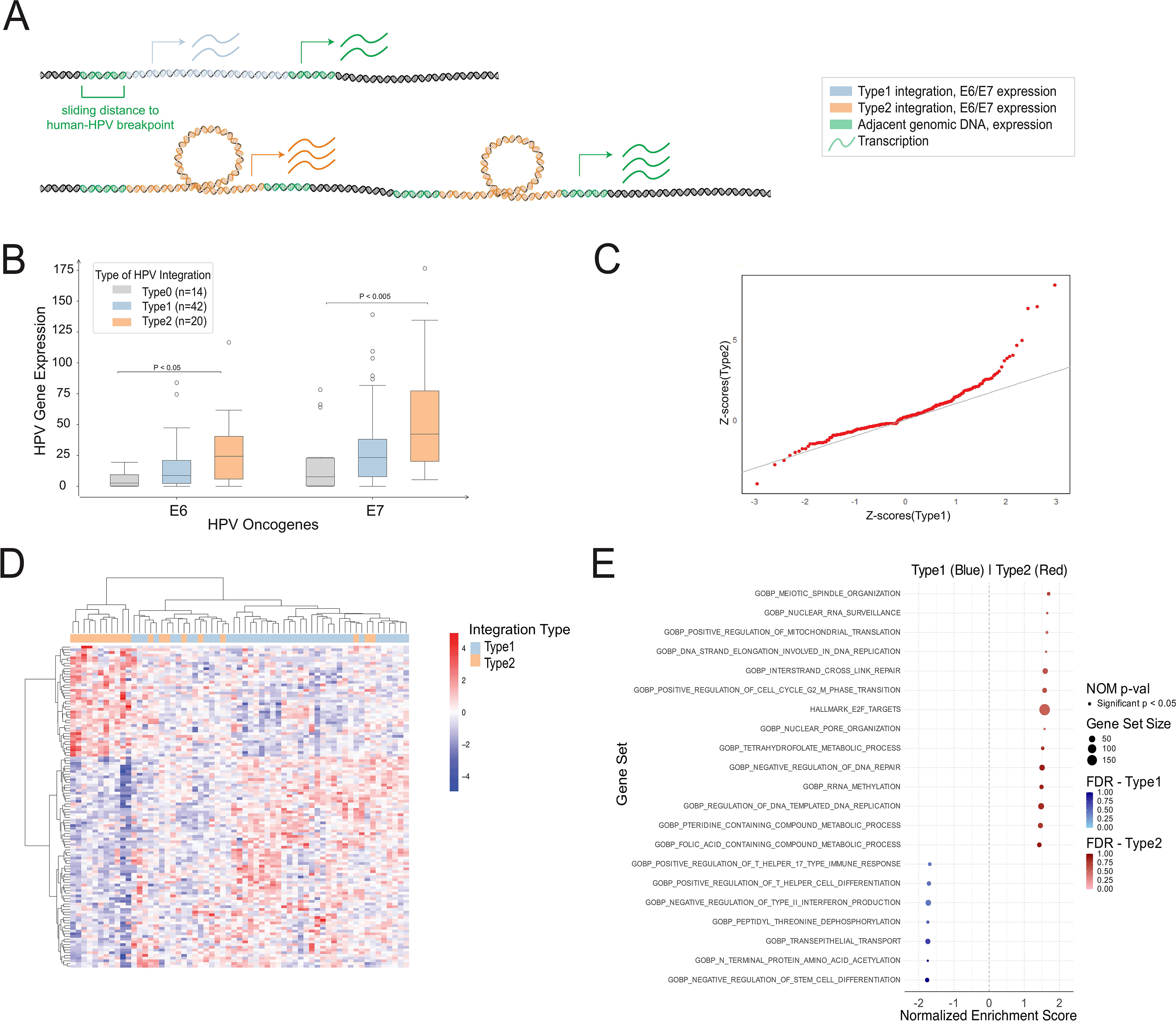
HPV integration type is associated with distinct gene-expression programs and immune landscapes in HPV+ OPSCC. **A.** Schematic representation of relationship of HPV integration with adjacent gene expression. **B.** Comparison of HPV oncogene expression between Type0, Type1 and Type2 integration status using the Wilcoxon rank-sum test to assess statistical significance. **C.** Q–Q plots comparing the distributions of gene expression levels for genes associated with Type2 events versus those associated with Type1 events. Deviations from the diagonal indicate differences in the underlying expression distributions between the two gene sets. **D.** Heatmap of normalized gene expression matrix clustering between Type1 and Type2 samples after batch correction. Genes with |log_2_ fold change| > 1 are shown in the clustered heatmap, highlighting distinct expression profiles. **E.** Gene Set Enrichment Analysis (GSEA) of Type1 vs. Type2 samples, showing enriched biological pathways associated with differential gene expression between Type1 and Type2.

To assess the effect of HPV integration on the expression of nearby host genes, we calculated gene-expression Z-scores across the 76 OPSCC tumors. We then compared the Z-scores of genes located within 500-kb of each integration event in a given tumor with those of the same genes in all other tumors in the cohort, enabling comparison of the local transcriptional effects of Type1 and Type2 integrations. Q-Q plots were used to visualize the expression level distributions of genes associated with Type2 events versus those associated with Type1 events (**Figure 4C**). The analysis revealed that, in tumors with Type2 breakpoints, adjacent genes were frequently overexpressed and occasionally under expressed compared to Type1 integrations, as evidenced by the significant diversion of many data points from the line of identity (KS test; p_val < 0.005). Distance of the host gene from the integration breakpoint also impacted expression, with Type2 integrations driving outlier expression of genes at a greater distance from the integration site than Type1 integrations (**Table S7**).

Given the impact of integration structure on the expression of HPV-host fusions, HPV genes, and integration-adjacent transcripts, we then sought to derive *de novo* signatures that differentiated the transcriptional profiles of tumors with Type1 and Type2 integration structures. Through differential gene expression analysis, we identified 94 significantly differential transcripts (p_val < 0.005) (**Figure 4D, Table S8**). Gene Set Enrichment Analysis (GSEA)^40,41^ further revealed distinct pathway alterations between Type1 and Type2 integration. Type1 tumors were enriched for seven pathways (p_val < 0.05), whereas Type2 tumors were enriched for fourteen pathways (p_val < 0.05; **Figure 4E**). These findings show that different integration types are associated with different gene expression programs and downstream biological pathways. Further, we evaluated whether Type2 tumors were enriched for gene signatures recently found to be active in HPV+ single cell states. We therefore compared signature scores across Type0, Type1, and Type2 tumors using the recently defined HPVon signature.^42^ This analysis showed that Type2 tumors trended towards having higher HPVon signature scores (**Supplemental Figure 16A**).

Using the overall cohort, and knowing that fusion status was significantly associated with survival **(Figure 4A)**, we then tested for relationships between HPV fusion status and transcriptional signatures. This demonstrated that fusion-positive tumors had significantly higher *E6* and *E7* expression than fusion-negative tumors (**Supplemental Figure 17A**). Through subsequent differential gene expression analysis, we identified 100 significant differentially expressed transcripts between samples with and without a HPV-host fusion transcript (**Supplemental Figure 17C)**. GSEA of this fusion signature identified 4 significant pathways involved in adhesion, migration, and apoptosis, enriched in fusion-negative tumors and 7 pathways, including those regulating CDK activity, centrosome separation, protein turnover and DNA cross link repair that were enriched in fusion-positive samples (**Supplemental Figure 17D**).

Given previously reported associations between T-cell infiltration and survival by our group and others in HPV+ OPSCC,^43–45^ we then tested for relationships of HPV integration structures and HPV fusions with inferred features of the surrounding tumor immune microenvironment. Thus, we tested for relationships with CytoSig inferred cytokine content and Cibersort inferred immune content (**Supplemental Figure 18**). This analysis revealed that Type2 tumors showed a significant depletion of CD4 memory T cells (Kruskal-Wallis Test: p_val < 0.05) (**Supplemental Figure 16B**). Likewise, both Ly endothelial cell abundance and BMP4 cytokine activity were also significantly reduced in Type2 tumors (Kruskal-Wallis Test: p_val < 0.05) (**Supplemental Figure 16C-D)**. Testing relationships with fusion status revealed that fusion-positive tumors also had a trend of lower CD4 abundance content (CD4⁺ T-cells, CD4⁺ Tcm, CD4⁺ Tem, CD4⁺ memory T-cells, CD4⁺ naïve T-cells) **(Supplemental Figure 18-C)**. Moreover, cytokine scores for BMP2, BMP4, FGF2 exhibited a trend toward depletion in the fusion-negative group **(Supplemental Figure 18-D)**. Interestingly, this is consistent with data showing that HPV+ OPSCC tumors enriched with CD4 memory cells, including CD4 cells with HPV memory, are associated with good survival outcomes,^44^ and suggests that fusion-positive tumors, which tend to have Type2 integrations, may have poor outcome due to low CD4 memory content.

Next, we analyzed the correlation between inferred cytokine or immune cell content with HPV gene expression and the HPVon score. Notably, several cytokines (IL12, IL1A, ECF, GM-CSF, and VEGFA) and immune cell populations (TGD, CLP, and CD8⁺ naïve T cells) showed positive correlations with *E6* and *E7* expression as well as the HPVon score (Pearson *r* > 0.4) (**Supplemental Figure 17B**), while MCSF and FGF2 were negatively correlated with *E1, E2, E6, E7*, and the HPVon score (Pearson *r* < - 0.4) and several stromal and immune populations, including endothelial cells, CD4⁺ Tcm, and CD4⁺ Tem cells were negatively correlated with the HPVon score (Pearson *r* < - 0.4), showing that HPV activity was strongly immunosuppressive in our cohort.

Finally, given that we observed that HPV integration events and HPV-host fusions converged to disrupt common pathways including HIPPO/YAP, non-canonical NFKB, MYC, and epithelial stemness, we then scored the activity of each of these pathways in the HPV16+ OPSCC RNAseq cohort using ssGSEA and tested for correlations of pathway activity and either inferred cytokine signaling or inferred immune content in the TME. This showed that activity of non-canonical NFKB strongly correlated with inferred memory CD8+ (Pearson correlation test p_val < 0.05) and CD4 (Pearson correlation test p_val < 0.001) cell score, while HIPPO/YAP, epithelial stemness or MYC activity scores were inversely correlated with CD4, CD8 and CD20 cell content (Pearson correlation test p_val < 0.005) (**Supplemental Figure 19A**). These data were consistent with the cytokine profiles, non-canonical NFKB activity strongly correlated with interferon gamma and TNF alpha activity (Pearson correlation test p<0.005). In contrast, HIPPO/YAP and epithelial stemness activity score were strongly inversely correlated with interferon gamma (Pearson correlation test p_val < 0.005), but highly correlated with TGF Beta (Pearson correlation test p_val < 0.0001) (**Supplemental Figure 19B**). Together, these data suggest that convergent disruption of pathways through the action of both the viral oncogenes as well as through integration and viral-host fusion gene mechanisms lead to convergent pathway disruption that is associated with robust cytokine and immune infiltration phenotypes in many cases.

### HPV integration architectures exhibit heterogeneity and clonal evolution during recurrence

Our initial cell line panel included UPCI:SCC90, derived from a primary tumor at the base of the tongue, and UPCI:SCC152, derived approximately one year later from a recurrent lymph node at the hypopharynx. This pair of cell lines provided an opportunity to examine changes in HPV integration structures over time. As expected for a clonal HPV integration event, both cell lines harbored two predominant HPV–HPV junctions with high copy number.

Interestingly, however, UPCI:SCC90 displayed higher local copy number for both junctions. In addition, we unexpectedly identified five unique HPV-HPV junctions exclusively in UPCI:SCC90, albeit with low copy number (**Figure 5A**).

**Figure 5.**
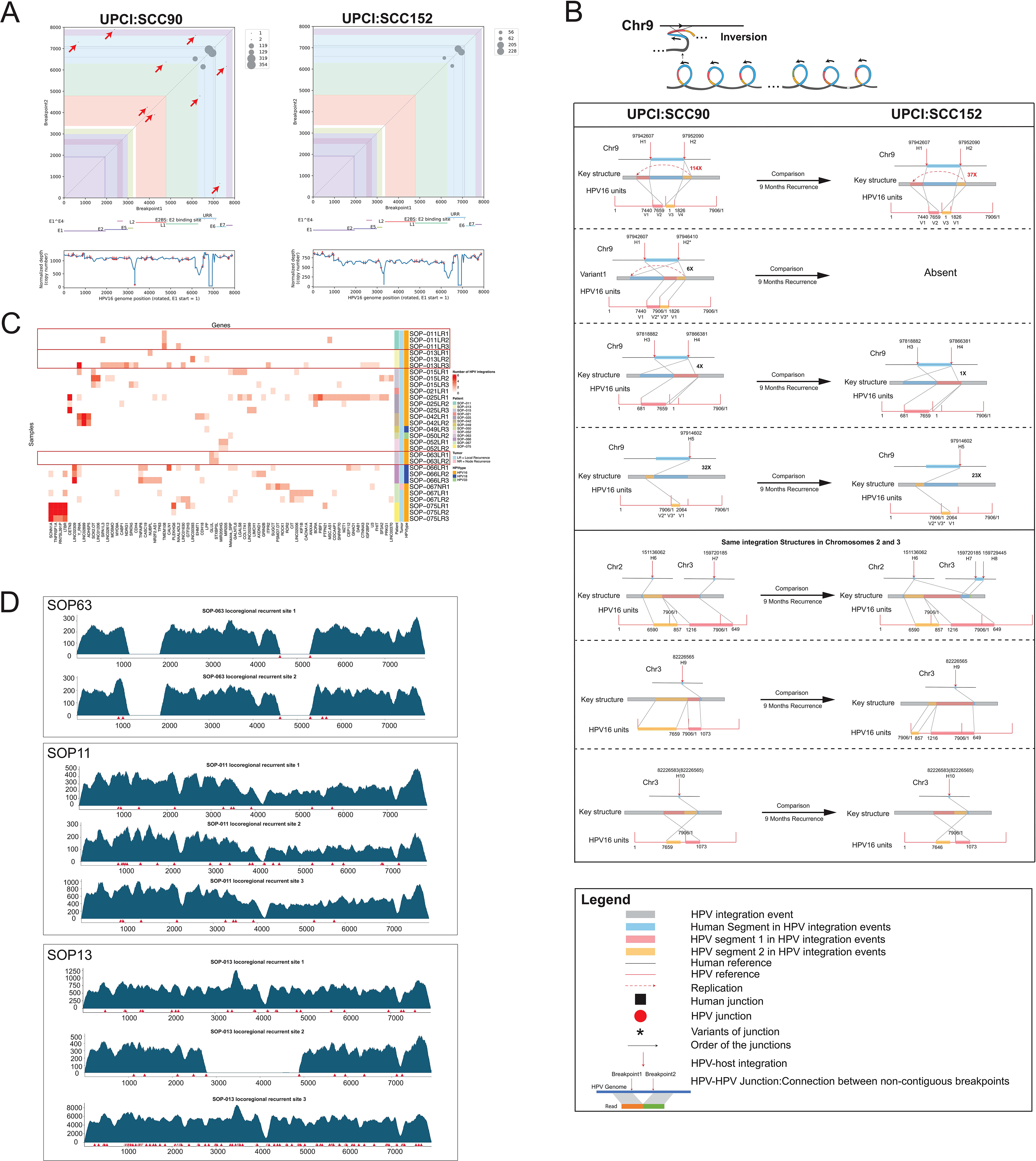
HPV integration events and structures demonstrate spatiotemporal heterogeneity and evolution over the course of HPV+ OPSCC progression. **A.** HPV-HPV junctions in UPCI:SCC90 and UPCI:SCC152 called from TCS. HPV-HPV junctions were plotted according to the coordinates of their two breakpoints as depicted in the schematic plot. The size of dots indicates the normalized copy number at HPV-HPV junctions. Red arrow indicates small unique HPV-HPV junctions that were detected only in UPCI:SCC90. **B.** Structure of one Type2 HPV event on chromosome 9 in UPCI:SCC90 and UPCI:SCC152 resolved from LR data. Only structures covered by more than 3 reads are displayed. **C.** Spatially distinct tumors from patients with HPV+ OPSCC were analyzed by TCS to characterize the distribution of HPV integration sites. Heatmap shows the number of HPV integration events associated with each cellular gene across samples. Columns represent cellular genes, and rows represent recurrent tumor samples. LR, local recurrence; NR, nodal recurrence. **D.** Representative HPV read-density plots from spatially matched locoregional recurrence sites in HPV+ OPSCC tumors: sites 1 and 2 for SOP63, and sites 1–3 for SOP11 and SOP13.

To elucidate the structural features of HPV integration events in these models, we applied our novel approach, as outlined in **Figure 1**, to resolve the underlying integration structures. A particularly complex integration event on chromosome 9 is shown in **Figure 5B**. This structure involved amplification of both human and HPV segments and was characterized by the presence of numerous human structural-variant inversions flanking the integration locus.

Although the core integration structure was conserved between the two models, the extent of amplification differed substantially, with copy numbers of 114X in UPCI:SCC90 and 37X in UPCI:SCC152. The host-HPV junctions were located at "*HEMGN* intron1 to *URR*" and "*E1* to intergenic upstream of *HEMGN*," while the HPV-HPV junctions fell within the *URR* regions. Two variant structures were shared by both models, with larger duplications observed in UPCI:SCC90. Host-HPV junctions for variant 2 were "*E6* to *PTCSC2* intron1" and "intergenic upstream of *PTCSC2* intron1 to *URR*"; for variant 3, they were "*TRMO* intron2 and *E1*." A unique structure, variant 1, was exclusively presented in UPCI:SCC90 as a variant of the key structure. Intriguingly, all three genes involved in this event were situated on the negative strand of the human genome. Considering that HPV genes are on the positive strand, the inversion structures suggested the potential for this complex rearrangement to be transcribed. Our results indicate that the heterogeneity of integration structures can be summarized by variations in (1) copy number at HPV integrations; (2) independent HPV-HPV, HPV-host, and host-host junctions; and (3) distinct characteristics of rearrangement breakpoints. Furthermore, by combining findings from TCS (**Figure 5A**) and LR (**Figure 5B**), we demonstrated that UPCI:SCC90 displayed an overall more complex rearrangement with various unique junctions and much higher local copy number, suggesting a potential clonal selection process from UPCI:SCC90 to UPCI:SCC152.

Building on this observation, we next sought to determine whether HPV integration heterogeneity was also present within our tissue cohort. From our cohort of 54 patients with recurrent HPV+ OPSCC, we selected 13 patients in whom HPV had integrated into cellular genes and performed HPV TCS across multiple independent spatial regions of each tumor (**Figure 5C**). Consistent with our cell line findings, this analysis revealed substantial heterogeneity in HPV integration profiles across spatially distinct tumor regions. We observed two key features of HPV integration heterogeneity: (1) substantial variation in the copy number of HPV integrations occurring within the same gene across independent recurrent sites, as exemplified by *SCNN1A, TNFRSF1A, LTBR,* and *PLEKHG6* across three independent lymph nodes; and (2) multiple unique HPV integration events across different samples from the same patients, further demonstrating the heterogeneity of HPV integration patterns within tumors. Consistent with this observation, we noticed pronounced differences in HPV integration structures and copy number among tumors sampled from distinct anatomical sites within the same patient, including recurrent tumors arising in independent lymph nodes (**Figure 5D**). For example, SOP63 exhibited distinct integration sites across tumors (highlighted in red), while SOP11 showed differences in both integration-site location and copy number. In SOP13, tumors differed not only in integration-site location and copy number, but also in the composition of the integrated HPV genome. Collectively, these multiregional TCS analyses revealed substantial heterogeneity in HPV integration sites, structural configurations, and copy number across spatially distinct tumor regions. The presence of both shared clonal integrations and private integration events, together with structural alterations of clonal integrations across tumor sites, supports a model of ongoing clonal evolution during tumor progression.

## Discussion

We introduced an innovative approach for resolving the structure of complex HPV integrations from long-read sequencing data, and developed a method to infer integration structures from short-read data, unlocking the ability to define HPV integration structures from FFPE samples. We established criteria for Type1 and Type2 HPV integration structures based on statistical and biological features, providing a potential standard cutoff for future studies. Notably, we identified several integration hotspots, including *CD274* (*PDL1*), *TP63* and *MYC*, which have been reported in previous research.^15,24,46,47^ Integration sites such as *HEMGN*, *TRMO*, *TRAF2*, *CASC11*, and *KLF4* were also detected in various studies.^15,24,48–51^ Specifically, *CASC11* and *PVT1* were previously identified as potential biomarkers in cervical cancer,^52,53^ and *KLF4* shown to play a role in the life cycle of HPV31 late viral gene expression.^54^ More importantly, by sequencing the largest cohort to date, we were able to demonstrate for the first time that HPV integrations converge on genes involved in key common pathways.

In our investigation, we unveiled distinctive transcriptomic profiles and gene sets associated with Type1 and Type2 HPV16 integration events. Type2 events were more likely to be transcribed, potentially resulting in the generation of complex chimeric transcripts featuring non-canonical fusion sites. The occurrence of such non-canonical fusions aligns with observations from prior research^15^ and we extend previous data showing that HPV-host fusion status is likely a poor prognostic feature of integrated tumors. Interestingly, oncogenes in proximity to integrations demonstrated elevated expression levels in Type2 events compared to Type1 events. This impact of HPV16 integrations on nearby genes diminished as the distance increased, notably becoming less pronounced beyond 200-kb, consistent with a previous analysis^15^ showing a gradual decline in the impact of HPV16 integration on expression when the distance reached 150-kb from the breakpoint. This validates a key quantitative metric for how integration events impact host transcriptional activity.

Our data show that diverse molecular mechanisms lead to convergent disruption of key complementary pathways including Hippo/Yap, Epithelial Stemness, Non-canonical NFKB and UV DNA damage repair. The activity of these pathways is strongly correlated with cytokine signaling and immune content in tumors, suggesting that disruption of these pathways may inform future therapeutic strategies. Interestingly, *E6* and *E7* have been directly linked to disruption of some of these pathways.^55,56^ These findings raise the possibility that the alternative molecular mechanisms of disruption identified here, such as integration or host gene fusion, serve to stabilize, or “lock in,” key signaling pathway alterations. Emerging evidence suggests that in certain cellular contexts, including specific squamous differentiation states^57^ or cells at the invasive leading edge of tumors,^42^ HPV oncogene expression and associated signaling programs can be transiently attenuated, potentially providing a context-dependent selective advantage. In such settings, stable genomic alterations that drive complementary pathway activity may help sustain oncogenic signaling even when viral gene expression is temporarily reduced.

Indeed, single cell studies have identified intratumoral heterogeneity of HPV expression and, more recently, of HPV integration events,^12,18^ suggesting that clonal evolution selecting for specific HPV-host events could be a key feature of recurrent tumors. Our approach is the first to reveal heterogeneity in integration structures, encompassing variations in local HPV copy number, distribution of specific HPV integration sites within structures, and distinctive characteristics of rearrangement breakpoints (human-HPV, HPV-HPV, human-human).

Building on these observations, we hypothesized that HPV integration events undergo clonal selection. To test this hypothesis, we compared the integration profiles of matched primary and recurrent tumors, focusing on one recurrent HPV+ model pair, UPCI:SCC90 and UPCI:SCC152, using long-read whole-genome sequencing. Our findings provide compelling evidence indicating the heterogeneity and clonal selection of HPV integration events during pathogenesis. Based on these data, we propose a new working model for disease pathogenesis, wherein the heterogeneity and clonal evolution of HPV integration events and structures play a pivotal role in driving oncogenesis and cancer progression.

In summary, we developed a framework to resolve complex local HPV integration events from long-read sequencing, and optimized a prediction algorithm to identify HPV integration structure from short-read data, thereby establishing a new standard for analyzing HPV integrations. Our findings demonstrate that Type2 HPV integration events are associated with transcriptional programs characteristic of aggressive tumors and suggest that heterogeneity in HPV integration may contribute to tumor progression. Together, these findings identify previously unrecognized candidate molecular biomarkers of HPV integration that warrant further investigation in translational studies.

## Resource availability

### Lead contact

Further information and requests for resources should be directed to, and will be fulfilled by, the lead contact, Dr. Chad Brenner.

### Data and code availability

Sequencing data have been deposited in the NCBI Sequence Read Archive (SRA) database under the accession numbers SRA: PRJNA1515176, PRJNA1515142, PRJNA1518217. Original code is available for download at our repository: https://github.com/mills-lab/HPV_heterogeneity

## Acknowledgments

This work was supported by National Institutes of Health (NIH, Grant number R01DE032699) awarded to A.P.B, J.C.B, and R.E.M and (NIH, Grant number R01DE034740). J.C. and E.U.G-M were supported by NIH T32 GM145470. X.Du was supported by NIH T32 HG000040. We also thank generous donation from Johnna & Justin Lesch in support of this research.

## Author contributions

Conceptualization, W.G., A.D.B., X.Du, X.Deng, R.E.M., and J.C.B.; Methodology, W.G., A.D.B., X.Du, X.Deng, A.P.B., H.J., R.E.M., and J.C.B.; Software, W.G.; Validation, W.G., A.D.B., X.Du, X.Deng, E.U.G.-M., J.C., C.V.B., and V.N.; Formal Analysis, W.G., X.Du, X.Deng, H.J., and R.E.M.; Investigation, A.D.B., X.Du, X.Deng, E.U.G.-M., J.C., C.V.B., V.N., and J.B.M.; Resources, M.B., K.A.C., S.E.C., D.F., K.M.M., M.L.M., M.E.P., J.L.S., A.G.S., C.L.S., P.L.S., F.P.W., P.Y., J.B.M., K.A.L.K., A.P.B., M.E.H.N., M.E.S., and J.C.B.; Data Curation, W.G., A.D.B., X.Du, X.Deng, E.U.G.-M., J.C., C.V.B., and V.N.; Writing - Original Draft, W.G., A.D.B., X.Du, X.Deng, R.E.M., and J.C.B.; Writing - Review & Editing, all authors; Visualization, W.G., A.D.B., X.Du, X.Deng and J.C.; Supervision, A.P.B., H.J., M.E.H.N., M.E.S., R.E.M., and J.C.B.; Funding Acquisition, R.E.M. and J.C.B.

## Methods

### Sample preparation and DNA extraction

DNA extraction from FFPE specimens was conducted as previously detailed.^58,59^ A board-certified head and neck pathologist (J.B.M.) identified the tumor areas. Both tumor and adjacent normal tissue cores were collected, and genomic DNA was extracted using the Qiagen Allprep DNA/RNA FFPE kit (Qiagen, Hilden, Germany). The DNA was quantified with a Qubit, following the methodology previously described.^59^ Subsequently, the DNA samples were sent to the University of Michigan Advanced Genomics core for library preparation and sequencing.

### Targeted capture sequencing library preparation

A minimum of 100 ng of genomic DNA from both tumor and adjacent normal tissue isolations was utilized for library preparation using the Rubicon DNA Thruplex (Rubicon Genomics, Ann Arbor, MI, USA) following the manufacturer’s instructions. Targeted capture was subsequently performed with our Nextera custom capture library, which includes high-density probes covering the HPV16 genome and coding exons of 225 genes described previously.^60^ These genes represent 744 kb and were selected based on their alteration in over 2% of tumors from the initial HNSCC TCGA data release or due to mutations identified in our internal sequencing data. Overall, our custom capture library encompasses approximately 0.023% of the genome and 2.47% of the exome. Next generation sequencing was carried out on an Illumina HiSEQ4000 (Illumina, San Diego, CA, USA) using the paired-end 150 nucleotide reads.

### Reference genome construction

Custom hybrid reference genomes were generated for HPV integration and fusion analyses by concatenating the human reference genome with HPV reference sequences. For TCS and LR DNA sequencing analyses, the GRCh38 human reference genome from ENSEMBL was concatenated with HPV16 reference genome obtained from the PaVE database.^61^ For RNA-seq–based fusion detection, GRCh38 was concatenated with a transformed HPV16 genome (RNAref) to avoid disrupting HPV transcripts that span the arbitrary linear origin of the circular HPV genome.^62^ The RNAref is constructed by shifting the starting position of the circular genome to the PaVE HPV16 genome basepair 23.

### Targeted capture sequencing preprocessing and somatic variant analysis

Raw FASTQ files generated from the Illumina sequencing platform were initially assessed for quality using FastQC v0.11.5. Low-quality bases and adapter sequences were removed using Trim Galore v0.4.5. The preprocessed reads were then aligned to the human reference genome using BWA v0.7.15. Mapped reads were subsequently sorted, deduplicated, and indexed with PicardTools v2.4.1. Base quality score recalibration was performed using GATK v3.6, generating high-quality aligned reads for downstream variant analysis.

For each tumor–normal pair, pileup files were generated using SAMtools v1.9. Somatic variant calling was performed with VarScan v2.4.1 in somatic mode using default parameters. The resulting variants were annotated with ANNOVAR. Variants located in intronic or intergenic regions were excluded, and only variants supported by at least five alternate allele reads in the tumor sample were retained as candidate somatic variants.

### Targeted capture-based HPV integration detection

HPV–host integration events were initially identified in targeted capture sequencing samples using SearcHPV.^24^ HPV integration sites identified from targeted capture sequencing were used to guide downstream long-read analyses.

### Copy-number normalization in targeted capture sequencing

To normalize copy number estimates from targeted capture sequencing, we first evaluated genes with stable read depth across samples after normalization by total sequencing depth. Four genes, *E2F2, MAP2K4, CD52,* and *PPM1D*, showed low variance in normalized read depth across targeted capture sequencing samples. To correct for probe-related depth bias, *MAP2K4* was selected as the background reference gene for copy number normalization across all targeted capture sequencing samples. Following normalization, *E6* and *E2* read counts at probe sites were validated by ddPCR.

### Oxford nanopore library preparation and sequencing

Library preparation and Oxford Nanopore sequencing was carried out in the Advanced Genomics Core at the University of Michigan. For the initial quality control, samples were cleaned with Ampure XP beads as needed, to ensure they met the purity standards required by Oxford Nanopore Technologies (ONT).

Each sample was prepared separately for sequencing using the Ligation Sequencing Kit 9 (SQK-LSK109). Sequencing was performed on individual GridION R9.4.1 flow cells (FLO-MIN106) with 2 to 3 library loads for each sample. A nuclease wash was performed between each loading to recover the maximum number of available pores. Basecalling was conducted with Guppy v5.1.13 to convert the raw sequencing data into FASTQ reads.

### Long-read alignment and reconstruction of HPV integration events

Nanopore reads were aligned to the custom hybrid genome described in the Reference genome construction section, comprising GRCh38 from ENSEMBL and HPV16 sequence from the PaVE database. Alignment was performed using Minimap2.^63^ Chromosomes harboring HPV integration events were identified based on integrations detected from targeted capture sequencing.

Split reads containing both human- and HPV-aligned segments were selected as informative reads for reconstructing integration structures. For each informative read, all junctions were identified, including human–human, human–HPV, and HPV–HPV junctions. Each junction was represented as a node within a tree-based data structure.

In the initial filtering step, junction sites located within 50-bp of one another were collapsed into a single junction cluster, and the most frequently observed coordinate within each cluster was retained as the representative junction site. Tree structure was initiated using the most abundant junction across all informative reads as the root node. From this root, the structure was extended iteratively in both the 5′-to-3′ and 3′-to-5′ directions in two separate tree structures. After each round of extension, remaining informative reads were recursively processed, and their junctions were added to the tree until no informative reads remained.

The resulting two tree structures were traversed to enumerate all possible integration paths or assemblies, and read support was calculated for each junction. Because loops cannot be directly represented in a tree structure, redundant junctions observed within individual reads were recorded separately and used to resolve looped configurations. Read names were retained at each node to facilitate assignment of reads to distinct integration paths. SAMtools was used throughout the pipeline for BAM file processing.

### Classification of HPV integration types

HPV integration events were categorized according to structural complexity. Integrations located within 500-kb of one another were grouped into a single integration cluster based on previously described criteria.^8,15^ Samples without detectable HPV integrations were designated Type0 (e.g. episomal, non-ecDNA).

Type2 integrations were defined as events with a local copy number >1 at the host–HPV breakpoint, a maximum copy number >2 across the integration locus, and more than two HPV integrations within the same cluster. All remaining integration events were classified as Type1.

### Comparison of HPV integrations detected by nanopore and targeted capture sequencing

To avoid overcounting HPV integration events associated with complex rearrangements, we first grouped integrations occurring within 500-kb of one another as a single clustered integration event, based on criteria used in previous studies.^8,15^ Sequencing alignments around unique non-reference viral–host junctions can be imprecise, and many fusion or integration callers therefore apply small flanking windows, typically 10–50-bp, to merge nearby junction calls.^24,64,65^ Moreover, comparisons across different integration-detection methods have reported coordinate deviations exceeding 100-bp, with some studies using windows up to 200-bp to define shared HPV integration events.^8,17^ In this study, after grouping integrations into 500-kb clustered events, we considered integration junctions detected by long-read sequencing and targeted capture sequencing to represent the same junction if their closest coordinates were within 20-bp.

### RNA-sequencing library preparation and analysis

Sections of tissue with over 60% tumor content were identified from the corresponding H&E by pathologist Jonathon B. McHugh, M.D. These regions were collected for RNA isolation using the Qiagen AllPrep Kit, as described.^66^ RNA samples that met our predefined quality criteria, assessed by Qubit and Bioanalyzer analysis, were subsequently advanced for next-generation sequencing. Total RNA was then submitted to the University of Michigan Advanced Genomics Core for library preparation and Next Generation Sequencing was carried out on an Illumina NovaSeq using the paired-end 150 nucleotide reads.

Sequencing data were preprocessed and quality controlled using Cutadapt v2.3 to remove adapter sequences and low-quality bases. The quality of the trimmed reads was subsequently evaluated with FastQC v0.11.8. Potential contamination was assessed using FastQ Screen, which enabled screening against multiple reference datasets to detect contaminant sequences. Trimmed and quality-checked reads were then aligned to the GRCh38 human reference genome from ENSEMBL using STAR v2.7.8a. Gene-level expression was quantified with RSEM v1.3.3 by estimating read counts assigned to annotated genes. All alignment parameters followed ENCODE RNA-seq analysis standards.

### HPV fusion and integration detection

RNA-seq data were analyzed for HPV–host fusion transcripts using SurVirus. Reads were aligned to the transformed HPV16-containing hybrid reference genome RNAref as described in the **Reference genome construction** section.

### Comparison of HPV integrations called from targeted capture sequencing and RNA-seq

We set out to define common integration sites identified from both TCS and RNA-seq. We categorized HPV events into two scenarios for the purpose of comparing DNA and RNA host-viral breakpoints: genic and intergenic. First, we defined integrations that are within 5kb upstream or downstream to each known human gene as genic. For these genic integrations, we treat integrations that share the same gene range between TCS and RNA as co-localized common events. Second, we compare the intergenic integrations between TCS and RNA; if they are on the same chromosome and are within 200-bp of each other, we label them as co-localized between DNA and RNA (**Figure 3C**).

### RNA-seq expression level analysis

Gene expression matrices were generated from RNA sequencing data by STAR^67^ and RSEM^68^, resulting in three expression measures: raw gene count, FPKM, and CPM. Batch effects among samples were corrected using the ComBat function^69^ and the *removeBatchEffect* function within the Limma package.^70^

To evaluate the potential impact of HPV integration on gene expression, analyses were performed in two ways: (i) by stratifying samples into the three HPV integration types, and (ii) by comparing HPV fusion-positive versus fusion-negative samples. HPV fusion-positive samples were defined as samples with detectable HPV-host fusion transcripts in RNA-seq data, whereas fusion-negative samples had no detected HPV-host fusion transcripts. Within each category, we utilized the gene expression matrices to investigate molecular and immunological features across integration types. Specifically, immune cell abundance and cytokine signaling activity were estimated using xCell^71^ and CytoSig,^72^ respectively. HPV gene expression and HPVon scores were quantified and calculated by HPVon and HPVoff signature genes previously defined by Puram et al.^42^ We reported gene expression as mean-normalized log2 CPM, and filtered to exclude genes with a mean read count < 1. HPVon and HPVoff signature expression is calculated as the mean normalized log2CPM expression of all genes in each signature. We then generated a random control gene list. To maintain consistency with the original study and control for baseline expression differences,^42^ genes were sorted by expression, then binned into 30 bins, to ensure lowly and highly expressed genes were included in each group. A random bin was selected, then 100 genes were randomly selected from this bin to use as a random control gene list. HPVon signature enrichment is calculated as the difference between the mean HPVon signature and the random control signature. Host–host fusion events were detected in each sample with Arriba.^73^ Statistical significance was determined using the Kruskal–Wallis test with Dunn’s post hoc correction (p_val < 0.05).

Differential expression analysis between Type1 and Type2 samples was performed using Limma,^70^ with significant genes visualized selected at p_val < 0.05 and |log2FC| > 1. Gene Set Enrichment Analysis (GSEA)^40,41^ was then conducted to identify biological pathways enriched among the differentially expressed genes, employing the Hallmark and Gene Ontology Biological Process (GO BP) gene sets. Pathways were prioritized based on normalized enrichment score (NES), adjusted p-value, and false discovery rate (FDR). Sample 3266-CB-12 was identified as an outlier by unsupervised analysis of global gene-expression profiles and was therefore excluded from downstream differential expression analysis.

### Survival analysis

Disease-free survival was calculated using Kaplan-Meier Analysis. Clinical characteristics, HPV16 integration status, and pre-determined genomic and transcriptomic variables were tested in univariate analysis. Clinical variables included disease presentation, tumor stage/T stage, and other clinicopathologic factors available for analysis. HPV16 integration variables included integration type, defined as Type0/episomal versus Type1 or Type2/integrated, the presence of integration into a human gene, integration into a cancer-associated gene as defined by the COSMIC cancer gene list, the number of HPV16 integration events, and the number of host genes with HPV16 integration. Pre-determined genomic variables included host mutation status in cancer-associated pathways, with particular focus on epithelial differentiation pathway alterations. Additional pre-specified pathway-level variables included MAPK signaling, RAS signaling, ERBB signaling, downregulated UV-response genes, PI3K–AKT–mTOR signaling, WNT/β-catenin signaling, TNFα–NF-κB signaling, Hippo signaling, MYC target genes, non-canonical NF-κB signaling, and TP63γ target genes. Pre-determined transcriptomic variables included host-HPV fusion events, host-host fusion events, and HPV16 fusion-mediated disruption of the above pathways. Variables significant on univariate analysis were included in a multivariate cox regression analysis. Chi-square test was used to evaluate the association between clinical variables and integration status. A p-value of 0.05 was considered statistically significant. All statistical analysis was performed with SPSS (v29) and statistical graphs were created with Prism Graph Pad (v9).

## Supplemental figure legends

**Supplemental Figure 1:** Copy number analysis of HPV integration events in UM-SCC-47.

**A.** Targeted Capture Sequencing and published WGS data show variable HPV16 copy number (10–50 copies) at integration breakpoints, with a consistent overall copy number of 50. The grey line is the read depth of human reference, and the yellow line is the read depth of informative reads supporting HPV integration. Red arrows indicate breakpoints on the human genome and blue arrows represent breakpoints on the HPV16 genome.

For TCS data, alignment files generated by the SearcHPV pipeline were used to calculate coverage. Sample-level alignment statistics and read counts were obtained using samtools stats with a minimum mapping quality of 50. Per-base coverage was calculated using samtools depth with a minimum base quality of 50, producing the chromosome, genomic position, and raw read depth at each reported position. Background coverage was estimated from one housekeeping gene: MAP2K4 (chr17:12020877–12143828). Mean MAP2K4 coverage was calculated by dividing the sum of per-base read depths by the number of positions with reported depth. This mean coverage was used as the sample background coverage (bgCov).

HPV integration coordinates were obtained from the filtered SearcHPV output file, call_fusion_virus/filteredSelectedContig.pickle. Raw read depth was extracted for each genomic position within ±50 bp of every integration breakpoint. TCS depth was converted to normalized copy-number signal using normalized depth = 2 × raw depth/bgCov, where a value of 2 represents the sample background level. The final TCS output contained the chromosome, genomic position, and normalized depth for each integration-site window. For ONT long-read data, read depth at each position was used directly as the copy-number estimate.

**B. PCR validation of complex integration structures.** Host-virus breakpoints were validated in cell line UD-SCC-2 by designing primers on either side of the predicted breakpoints. Left - predicted structures using our assembler. Colored arrows represent primers; black bars represent the predicted amplicon lengths for the primer pairs. Right - gel electrophoresis of PCR products, with primer pairs labeled above. Each predicted amplicon was visualized at the expected length. Lane K-M is included with no visible product to show the reverse orientation of the X-chromosome segment in this region (in contrast with the product generated by primer pair K-L). The assembler allowed for the prediction of long, complex structures with multiple host-virus breakpoints assembled in the correct orientations. Note: Please refer to Figure 1 for legend of the symbols.

**Supplemental Figure 2:** HPV integration in UM-SCC-104 corroborated by TCS data and Nanopore assembly from wtbdg2.

**Supplemental Figure 3:** HPV integration structures in additional HPV+ HNSCC samples. Summary of HPV integration events and structural variations across 13 additional HPV+ HNSCC cell lines and frozen tumor tissue samples, resolved using the novel approach described.

Note: Please refer to Figure 1 for legend of the symbols.

**Supplemental Figure 4:** Phased long read from 10X Linked-read sequencing data for a Type2 event in *FXON2* of PDX-294R.

**Supplemental Figure 5: Five-year disease-free survival by HPV genotype.** Kaplan–Meier analysis demonstrating no significant difference in median five-year disease-free survival between patients with HPV16 mono-infection and those infected with HPV18 or HPV33 (74.9% vs. 55.6%, respectively; p_val > 0.5).

**Supplemental Figure 6: A:** Five-year disease-free survival in Type1 and Type2 tumors by number of host genes with integration.

**B:** Bar plot showing the total number of host genes with integration events (blue bars, left axis) and the number of genes from the COSMIC Cancer Gene Census Tier 1 or 2 list (orange bars, right axis) identified in each tumor sample.

**Supplemental Figure 7:** Pathway enrichment analysis of HPV integration–associated genes identified across all tumors, including both Type1 and Type2 integration events.

**Supplemental Figure 8: Landscape of somatic alterations to human genes in HPV16+ OPSCC tumors from the overall UMICH cohort as summarized by maftools.** Targeted capture sequencing was completed on 225 human genes from each tumor and **A.** Variant classification, **B.** Variant type, **C.** SNV classification, **D.** Variants per sample, **E.** Variant classification summary, and **F.** the top ten mutated human genes are shown.

**Supplemental Figure 9: Lollipop plots showing distribution of somatic alterations in frequently mutated human genes in the overall HPV16+ OPSCC cohort.** Type and location of somatic alterations are illustrated along the protein sequence for each gene. Highly recurrent hotspot mutations, those identified in 4 or more tumors in the overall cohort are highlighted. All plots were generated using maftools.

**Supplemental Figure 10: Comparative pathway mutation landscape in episomal vs. integrated tumors.** Pathway-level comparison of mutation frequencies between episomal (n = 28) and integrated (n = 155) tumors using Fisher’s exact test. P-values were adjusted for multiple comparisons using the Benjamini–Hochberg FDR method. Only TP53/MDM2 mutations remained significant after correction (p_val < 0.01).

**Supplemental Figure 11: Kaplan-Meier survival analysis of 183 patients assessing the impact of clinicopathologic variables, integration status, and genomic events on overall survival.**

**A.** Five-year disease-free survival (DFS) for total cohort by primary or recurrent/metastatic tumor status. **B.** For the primary tumors, five-year DFS is seen stratified by T-stage, **C.** Integration type (Type0, 1, or 2), **D**. total number of host genes with integration, **E.** binary integration type (no integration or integration), **F.** epithelial differentiation mutation status. **G.** Multivariate analysis with cox regression analysis demonstrates hazard ratios, 95% confidence intervals, and p values for T stage, epithelial differentiation status, and integration type.

**Supplemental Figure 12:** Kaplan–Meier analysis of DFS stratified by the total number of HPV integration events in HPV-integrated tumors.

**Supplemental Figure 13: A.** Host-host fusion status survival analysis.

**B.** Overlap between tumors harboring HPV–host fusions and tumors harboring host–host fusions. Bars indicate the number of tumors with both fusion types, with HPV–host fusions only, or with host–host fusions only. Fisher’s exact test was used to evaluate the association.

**Supplemental Figure 14:** Gene fusion structures for MAPK9–DNMT1, MAPK9– RASGEF1C, NBEAL1–ERBB4, ALDH1L1–ERBB4, MET–KAT2B, C3orf52–TIGIT, and FGFR3–TACC3 identified across seven RNA-seq samples: 3266-19, 3266-200, 3266-290, 3266-291, 3266-315, 3266-34, and 3266-49 using STAR chimeric alignment and Arriba. Individual fusion events were detected in subsets of the samples analyzed. Plots show exon structures, breakpoint positions, read support, and predicted retained protein domains.

**Supplemental Figure 15:** Pathway enrichment analysis of fusion-associated genes revealed significant enrichment in ERBB4-related signaling, non-canonical NF-κB signaling, and IL6– JAK–STAT3 signaling pathways.

**Supplemental Figure 16:** Molecular and immune features across HPV integration types in HPV16+ OPSCC tumors.

**Supplemental Figure 17: Relationships between HPV fusion status and transcriptional signatures.**

**A.** HPVon Score and HPV oncogene expression comparison between fusion-positive and fusion-negative samples.

**B.** Correlation matrices for HPV gene expression with cytokine and immune cell abundance.

**C**. Heatmap of normalized gene expression matrix clustering between fusion-positive and fusion-negative samples after batch correction. Genes with log2FC > |1| are shown in the clustered heatmap, highlighting distinct expression profiles.

**D.** Gene Set Enrichment Analysis (GSEA) of fusion-positive and fusion-negative samples, showing enriched biological pathways associated with differential gene expression between fusion-positive and fusion-negative.

**Supplemental Figure 18: A-D.** Immune cell and cytokines score by HPV integration types and HPV-Host fusion status.

**Supplemental Figure 19: A-B.** Immune and cytokines correlation matrix with HPV integration enrichment pathways.

## Supplemental Table legends

**Table S1:** TCS, RNA-seq and Long read Nanopore sequencing of 17 cell lines and tumor samples.

**Table S2:** Clinical, genomic, and HPV integration characteristics of 197 OPSCCs (183 HPV16+ OPSCC cases in sheet2).

**Table S3:** Association between HPV16 integration–associated pathway disruption and survival outcomes.

**Table S4:** Clinicopathologic and genomic variables associated with integration status.

**Table S5:** Clinical, HPV gene expression, gene fusion events, tumor microenvironment features of 102 HPV+ OPSCC samples (96/102 (94%) were HPV16+).

**Table S6:** 76 HPV16+ oropharyngeal squamous cell carcinoma (OPSCC) samples with available TCS data in S5 for HPV-host fusion analysis.

**Table S7:** Distance of host gene from integration breakpoint and impact on gene expression.

**Table S8**: Significant gene lists selected at p_val < 0.05 and |log2FC| > 1 between Type1 and Type2 for Figure 4D.

