## Supplemental Figure 1 for "Spatiotemporal Heterogeneity of HPV Integration Architectures Drive Oncogenic Features of Oropharyngeal Cancer"

### Slide 1
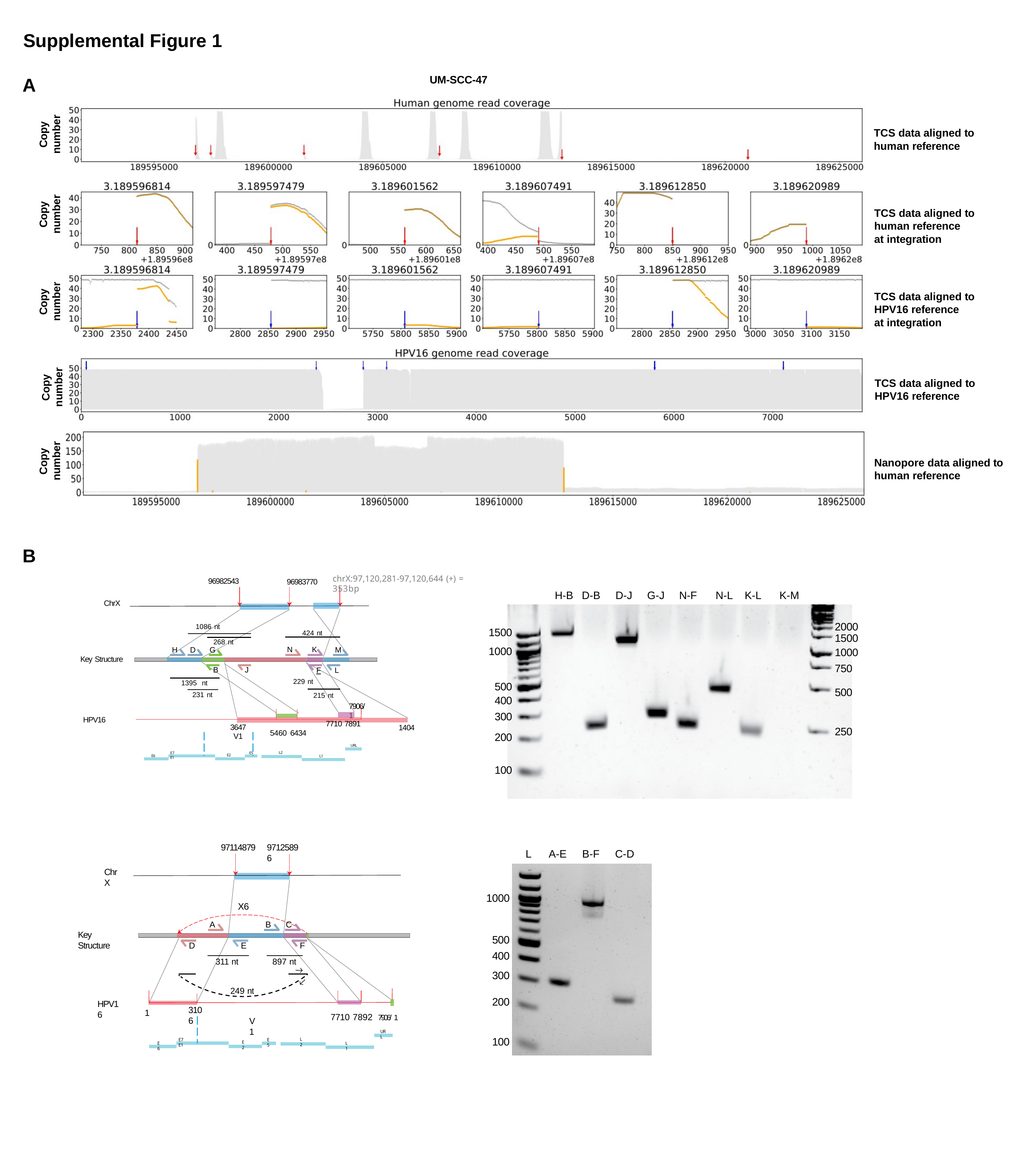

Supplemental Figure 1
UM-SCC-47
A
Copy number
TCS data aligned to human reference
Copy number
TCS data aligned to human reference
at integration
Copy number
TCS data aligned to HPV16 reference
at integration
Copy number
TCS data aligned to HPV16 reference
Copy number
Nanopore data aligned to human reference
B
chrX:97,120,281-97,120,644 (+) = 353bp
96982543
96983770
| | H-B | D-B | D-J | G-J | N-F | N-L | K-L | K-M |
| --- | --- | --- | --- | --- | --- | --- | --- | --- |
| 1500 | | | | | | | | |
| 1000 | | | | | | | | |
| 500 | | | | | | | | |
| 400 | | | | | | | | |
| 300 | | | | | | | | |
| 200 | | | | | | | | |
| 100 | | | | | | | | |
ChrX
2000
1500
1000
750
1086 nt
424 nt
N	K
	F	L
229 nt
268 nt
H	D	G
M
Key Structure
B
J
1395 nt
500
231 nt
215 nt
7906/ 1
HPV16
7710 7891
3647
V1
1404
250
5460 6434
URL
L2
E7 E1
E5
E2
E6
L1
97114879
97125896
| | L | A-E | B-F | C-D |
| --- | --- | --- | --- | --- |
| 1000 | | | | |
| 500 | | | | |
| 400 | | | | |
| 300 | | | | |
| 200 | | | | |
| 100 | | | | |
ChrX
X6
A
B	C
Key Structure
D
E
F
311 nt
897 nt
249 nt
HPV16
3106
1
7710 7892 ^7906/^ ^1^
V1
URL
L2
E7 E1
E5
E2
E6
L1
