## Supplemental Figure 2 for "Spatiotemporal Heterogeneity of HPV Integration Architectures Drive Oncogenic Features of Oropharyngeal Cancer"

### Slide 1
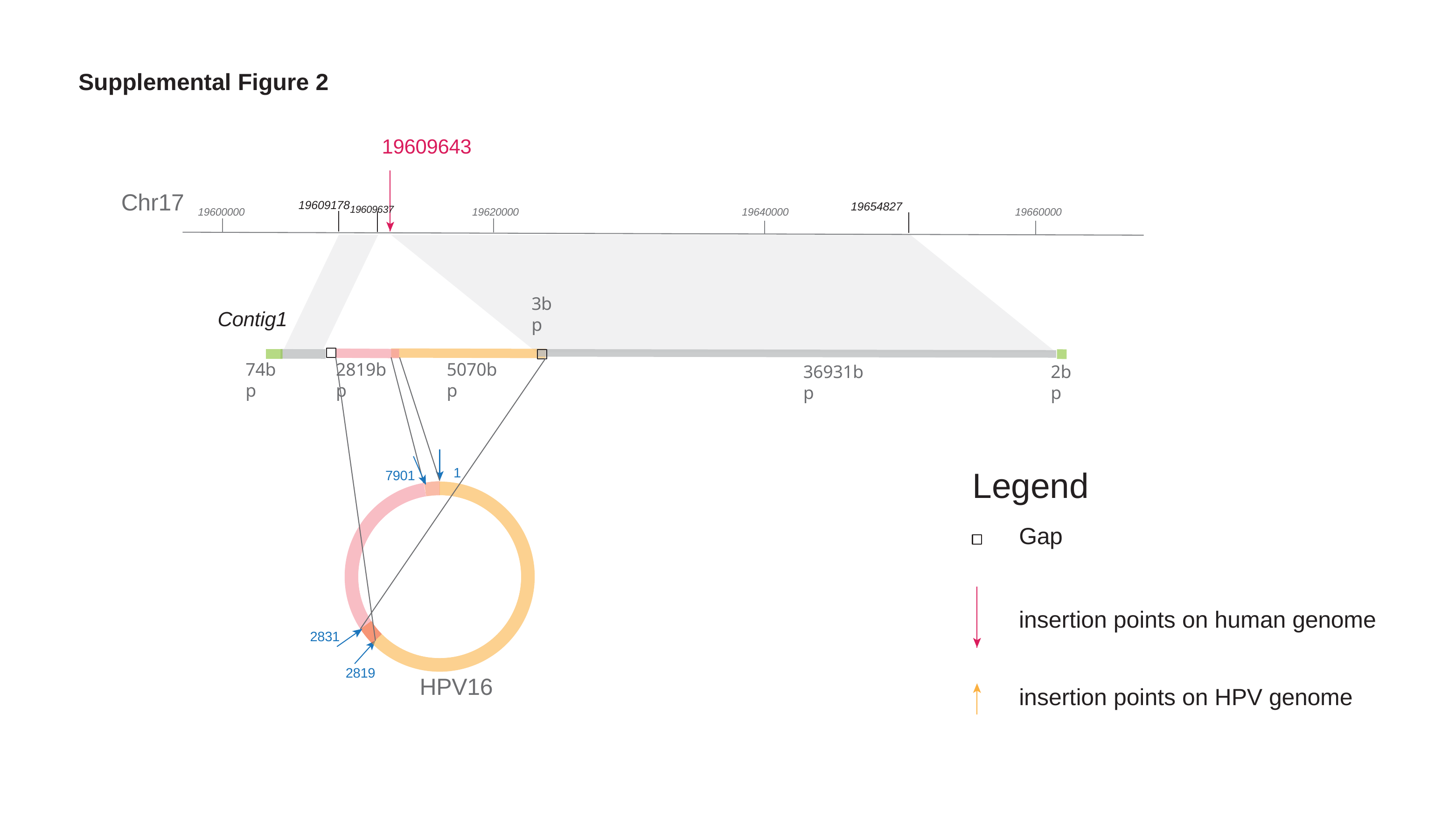

#### Supplemental Figure 2
19609643
Chr17
19609178_19609637_
19654827
19600000
19620000
19640000
19660000
3bp
Contig1
74bp
2819bp
5070bp
2bp
36931bp
Legend
Gap
1
7901
insertion points on human genome
2831
2819
HPV16
insertion points on HPV genome
