## Supplemental Figure 3 for "Spatiotemporal Heterogeneity of HPV Integration Architectures Drive Oncogenic Features of Oropharyngeal Cancer"

### Slide 1
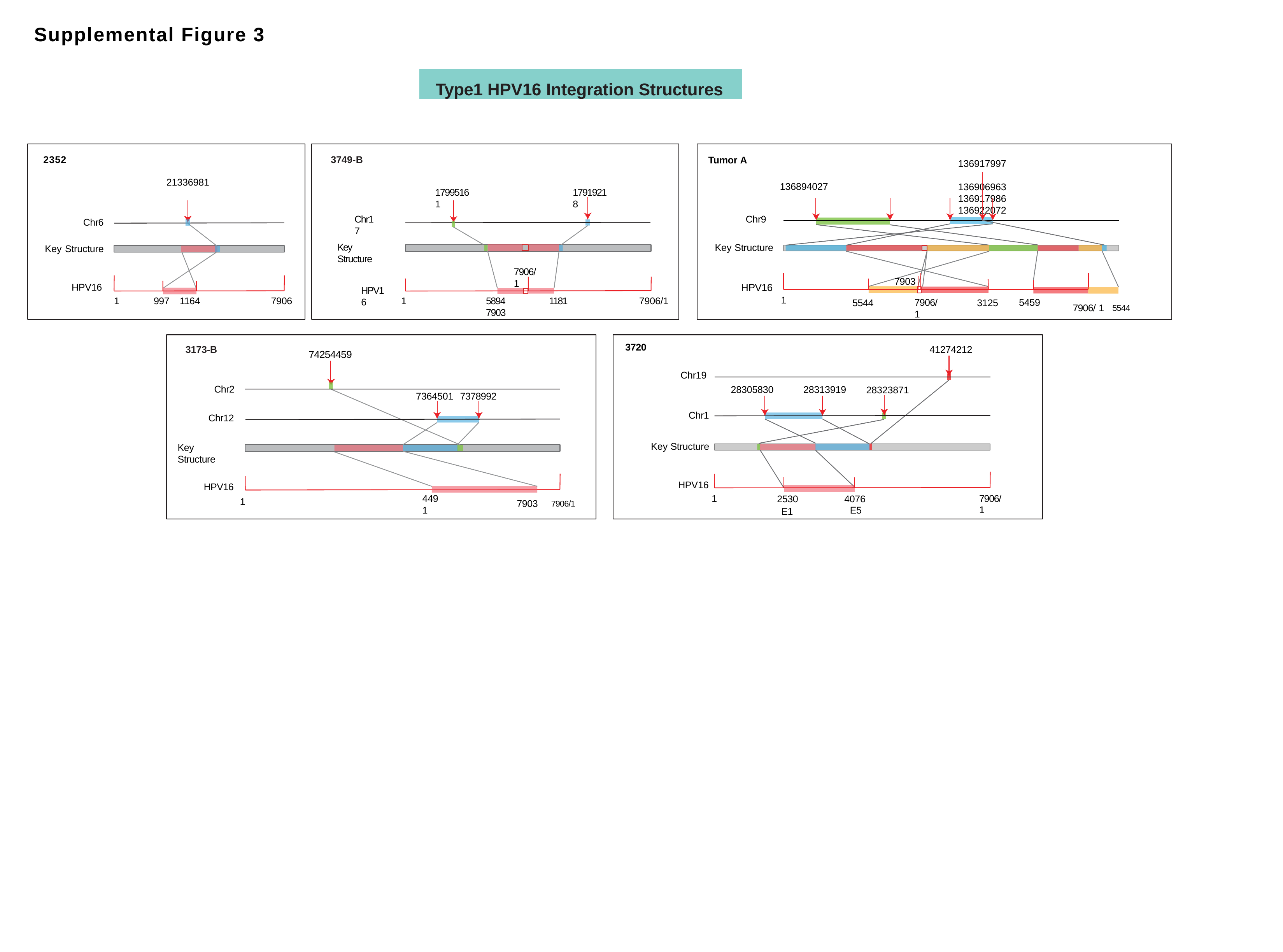

#### Supplemental Figure 3
Type1 HPV16 Integration Structures
2352
3749-B
Tumor A
136917997
136906963	136917986	136922072
21336981
136894027
17995161
17919218
Chr9
Chr17
Chr6
Key Structure
Key Structure
Key Structure
7906/1
7903
HPV16
HPV16
HPV16
1
7906/1
1181
1
997	1164
5894 7903
1
7906
7906/ 1
5459
3125
5544
7906/ 1	^5544^
3720
3173-B
41274212
74254459
Chr19
Chr2
28305830
28313919
28323871
7364501	7378992
Chr1
Chr12
Key Structure
Key Structure
HPV16
HPV16
2530
E1
7906/ 1
1
4491
7903	^7906/1^
4076
E5
1

### Slide 2
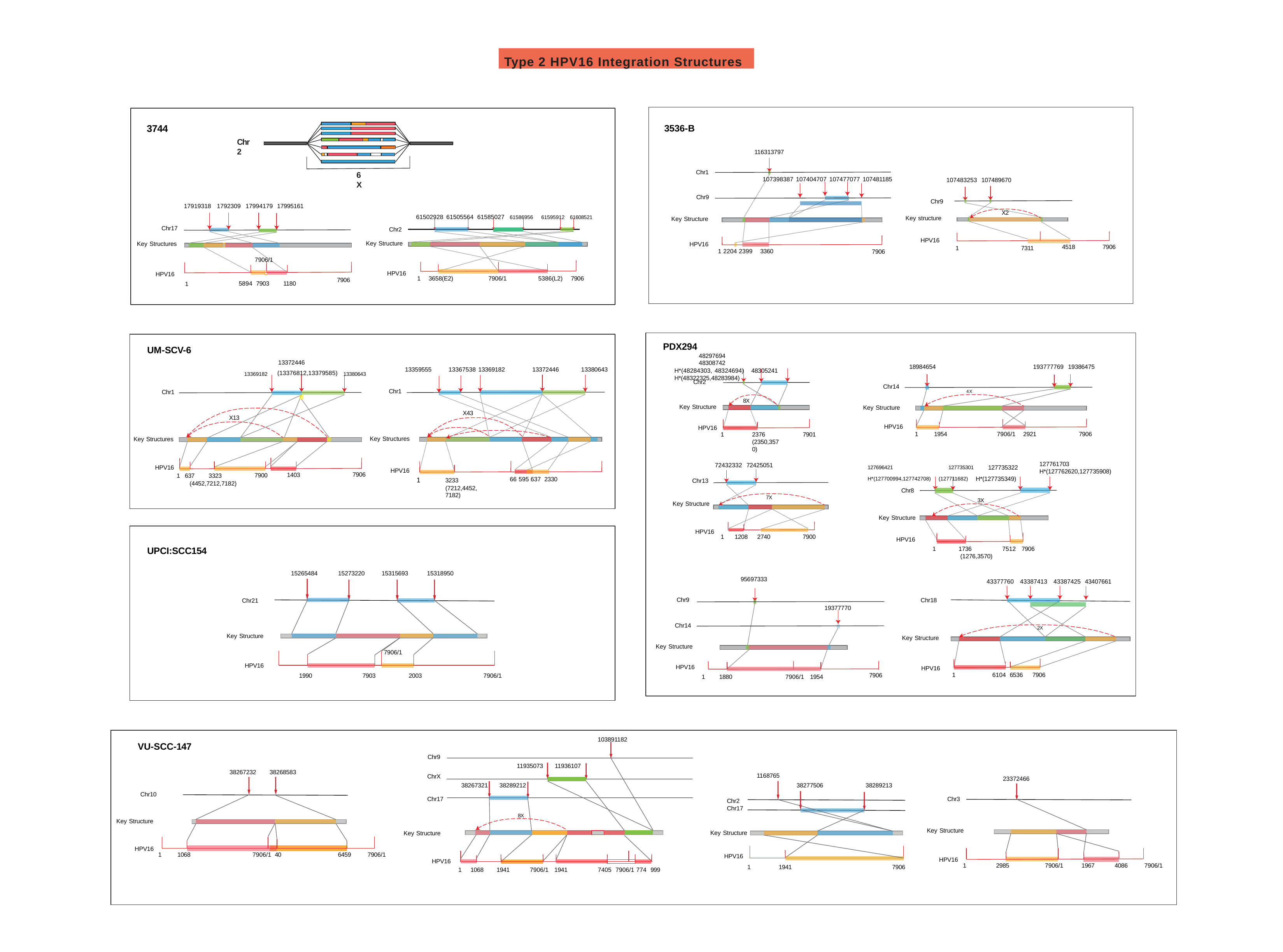

Type 2 HPV16 Integration Structures
3536-B
3744
Chr2
116313797
Chr1
6X
107398387 107404707 107477077 107481185
107483253 107489670
Chr9
Chr9
17919318 1792309 17994179 17995161
X2
61502928 61505564 61585027 ^61586956^ ^61595912^ ^61608521^
Key structure
Key Structure
Chr17
Chr2
HPV16
Key Structure
Key Structures
HPV16
7906
4518
1
7311
1 2204 2399 3360
7906
7906/1
HPV16
HPV16
1 3658(E2)
7906/1
5386(L2) 7906
_1_	5894 7903	1180
7906
PDX294
48297694	48308742
H*(48284303, 48324694) 48305241	H*(48322325,48283984)
UM-SCV-6
13372446
_13369182_ (13376812,13379585) _13380643_
18984654
193777769 19386475
13359555	13367538 13369182
13372446
13380643
Chr2
Chr14
Chr1
Chr1
4X
8X
Key Structure
Key Structure
X43
X13
HPV16
HPV16
1	1954
7906/1 2921
7906
1
2376
(2350,3570)
7901
Key Structures
Key Structures
127761703
H*(127762620,127735908)
^127696421^	^127735301^	127735322
^H*(127700994,127742708)^ ^(127711682)^ H*(127735349)
72432332 72425051
HPV16
HPV16
7906
1403
7900
1 637	3323
66 595 637 2330
1
3233
(7212,4452,7182)
Chr13
(4452,7212,7182)
Chr8
7X
3X
Key Structure
Key Structure
HPV16
1	1208	2740
7900
HPV16
UPCI:SCC154
1
1736	7512 7906
(1276,3570)
15265484
15273220	15315693	15318950
95697333
43377760 43387413 43387425 43407661
Chr9
Chr18
Chr21
19377770
Chr14
2X
Key Structure
Key Structure
Key Structure
HPV16
7906/1
HPV16
HPV16
1
6104 6536	7906
7906
7903
7906/1
2003
1990
1	1880	7906/1 1954
103891182
VU-SCC-147
Chr9
11935073	11936107
38267232	38268583
1168765
ChrX
23372466
38267321	38289212	38277506	38289213
Chr10
Chr17
Chr3
Chr2 Chr17
8X
Key Structure
Key Structure
Key Structure
Key Structure
HPV16
1	1068
6459	7906/1
7906/1 40
HPV16
HPV16
HPV16
1
2985
7906/1
1967
4086	7906/1
1
1941
7906
1	1068	1941
7906/1 1941
7405 7906/1 774 999
