## Supplementary figures and images for "Spatiotemporal Heterogeneity of HPV Integration Architectures Drive Oncogenic Features of Oropharyngeal Cancer"

### Supplemental Figure 4

#### Slide 1
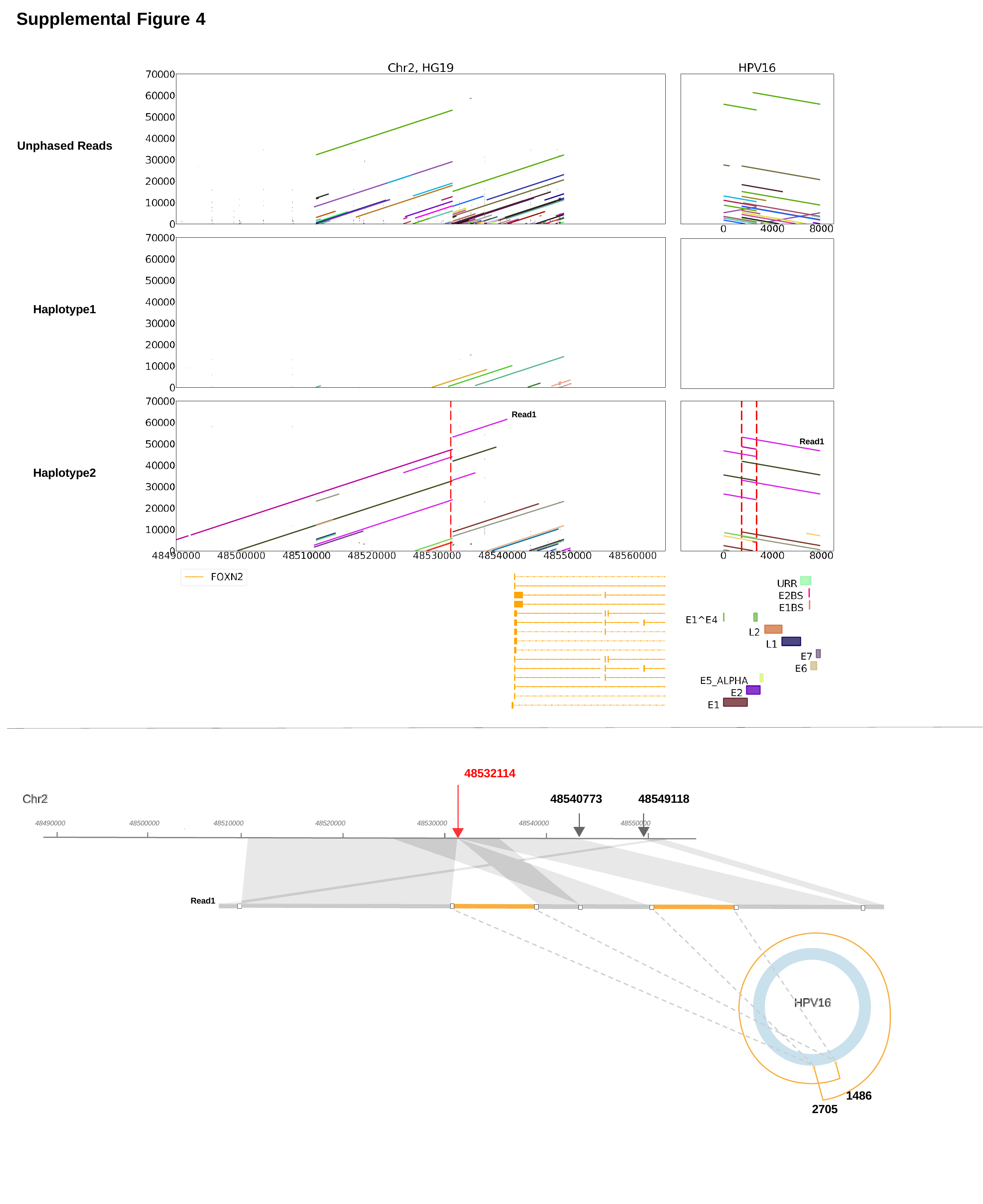

Supplemental Figure 4
Unphased Reads
Haplotype1
Read1
Read1
Haplotype2
48532114
Chr2
48540773
48549118
48530000
48490000
48500000
48510000
48520000
48540000
48550000
Chr2
Read1
HPV16
1486
2705

### Supplemental Figure 5

#### Slide 1
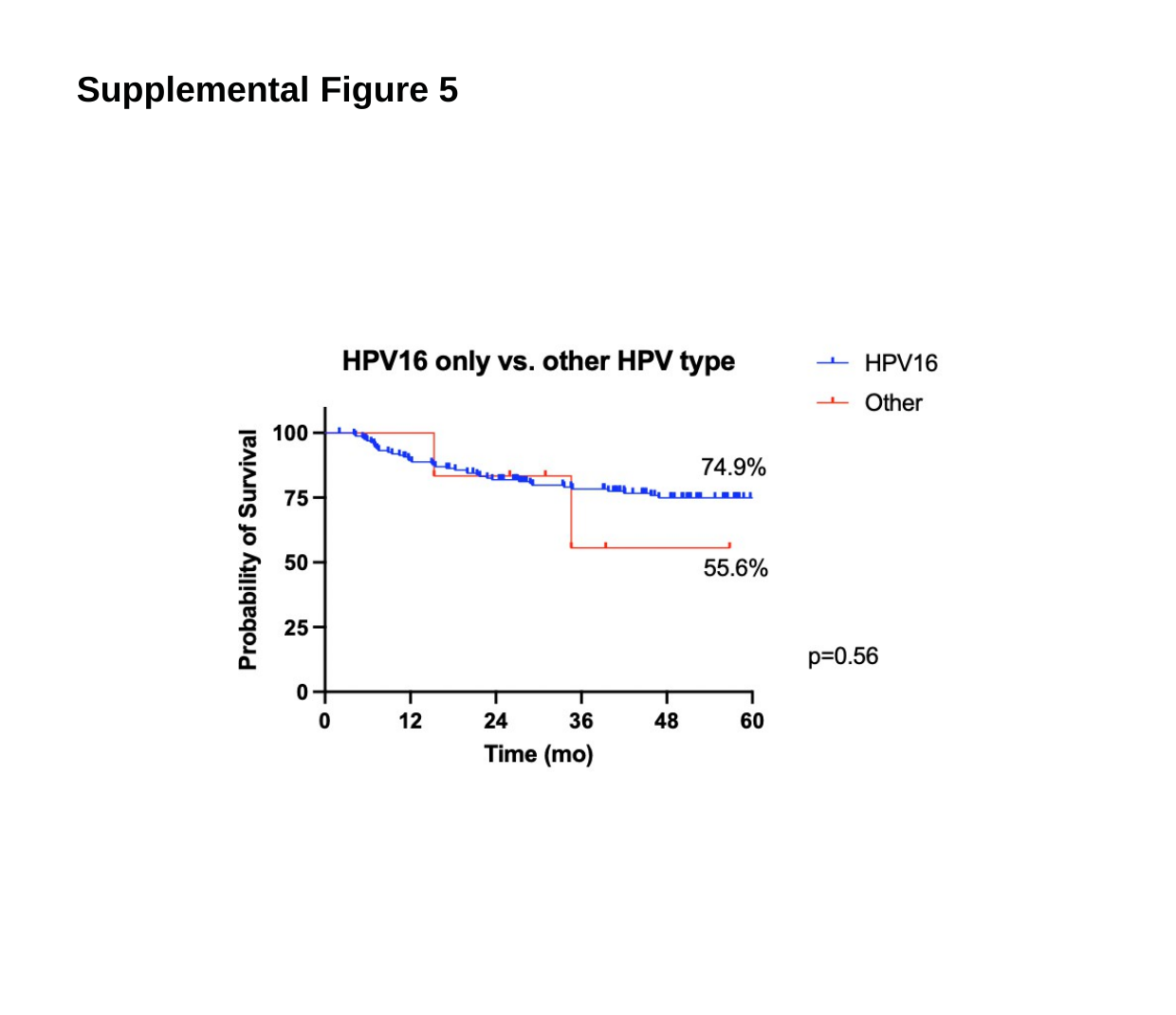

Supplemental Figure 5

### Supplemental Figure 6

#### Slide 1
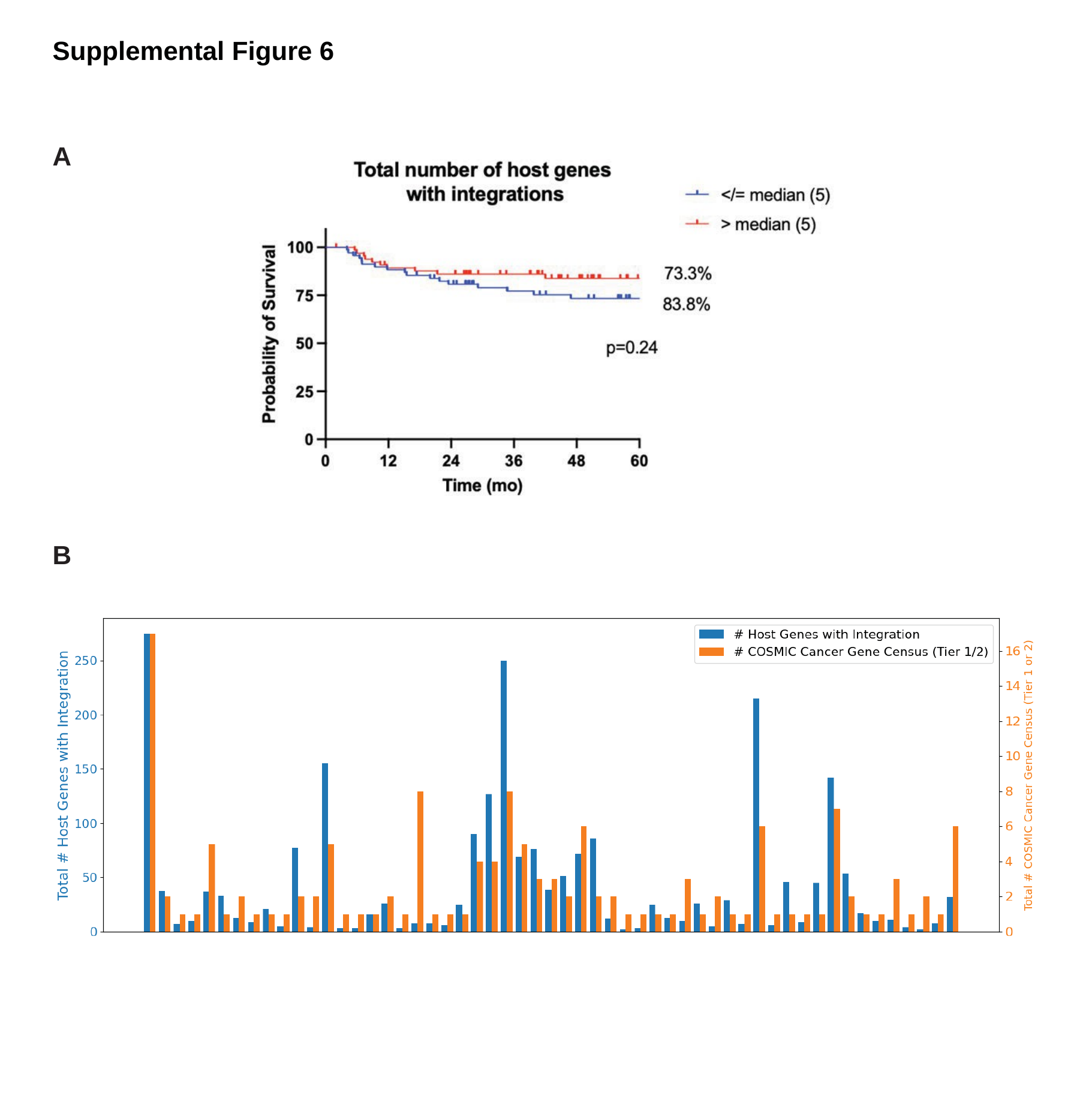

Supplemental Figure 6
A
B

### Supplemental Figure 9

#### Slide 1
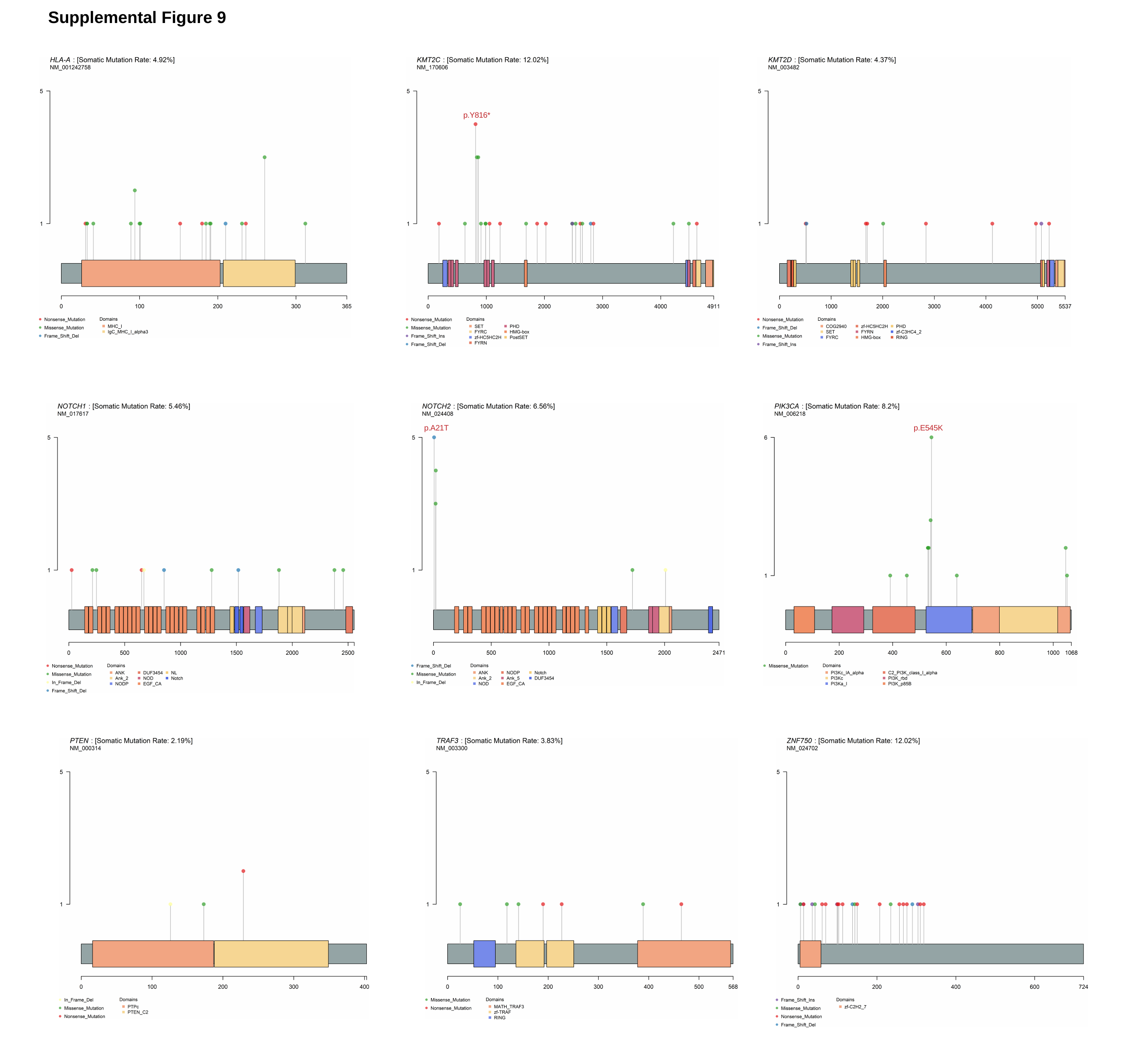

Supplemental Figure 9
p.Y816*
p.A21T
p.E545K

### Supplemental Figure 10

#### Slide 1
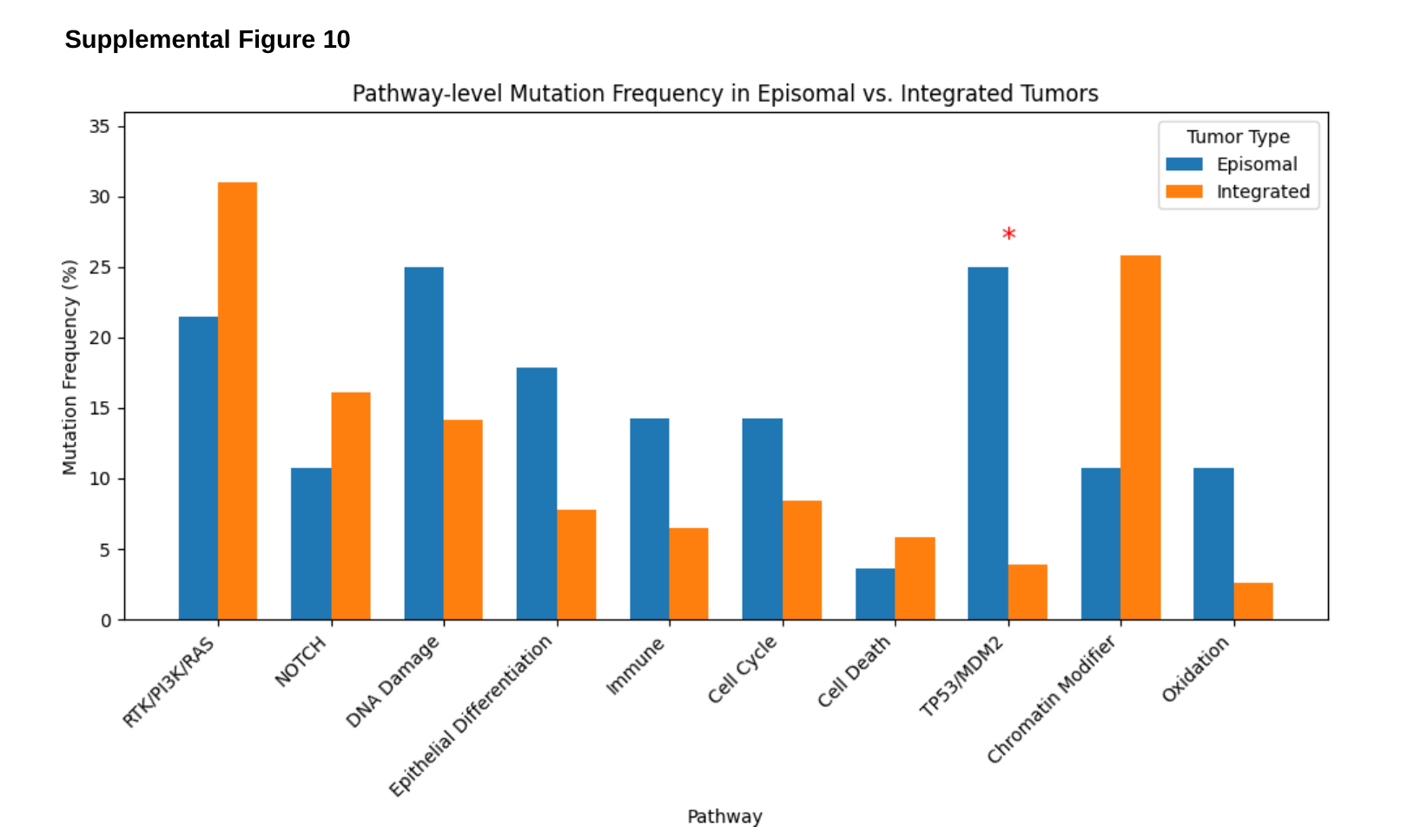

Supplemental Figure 10

### Supplemental Figure 12

#### Slide 1
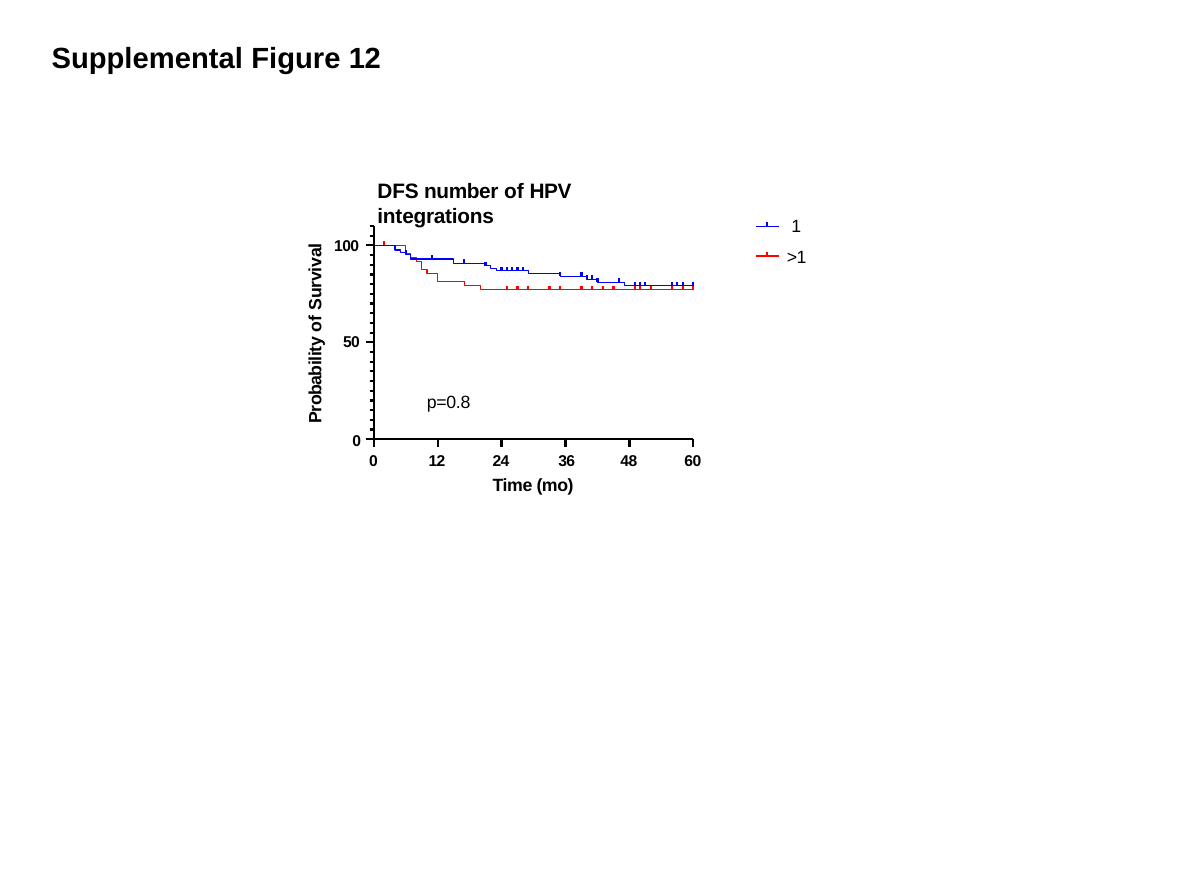

Supplemental Figure 12
DFS number of HPV integrations
1
>1
100
Probability of Survival
50
p=0.8
0
24	36
Time (mo)
0
12
48
60

### Supplemental Figure 18

#### Slide 1
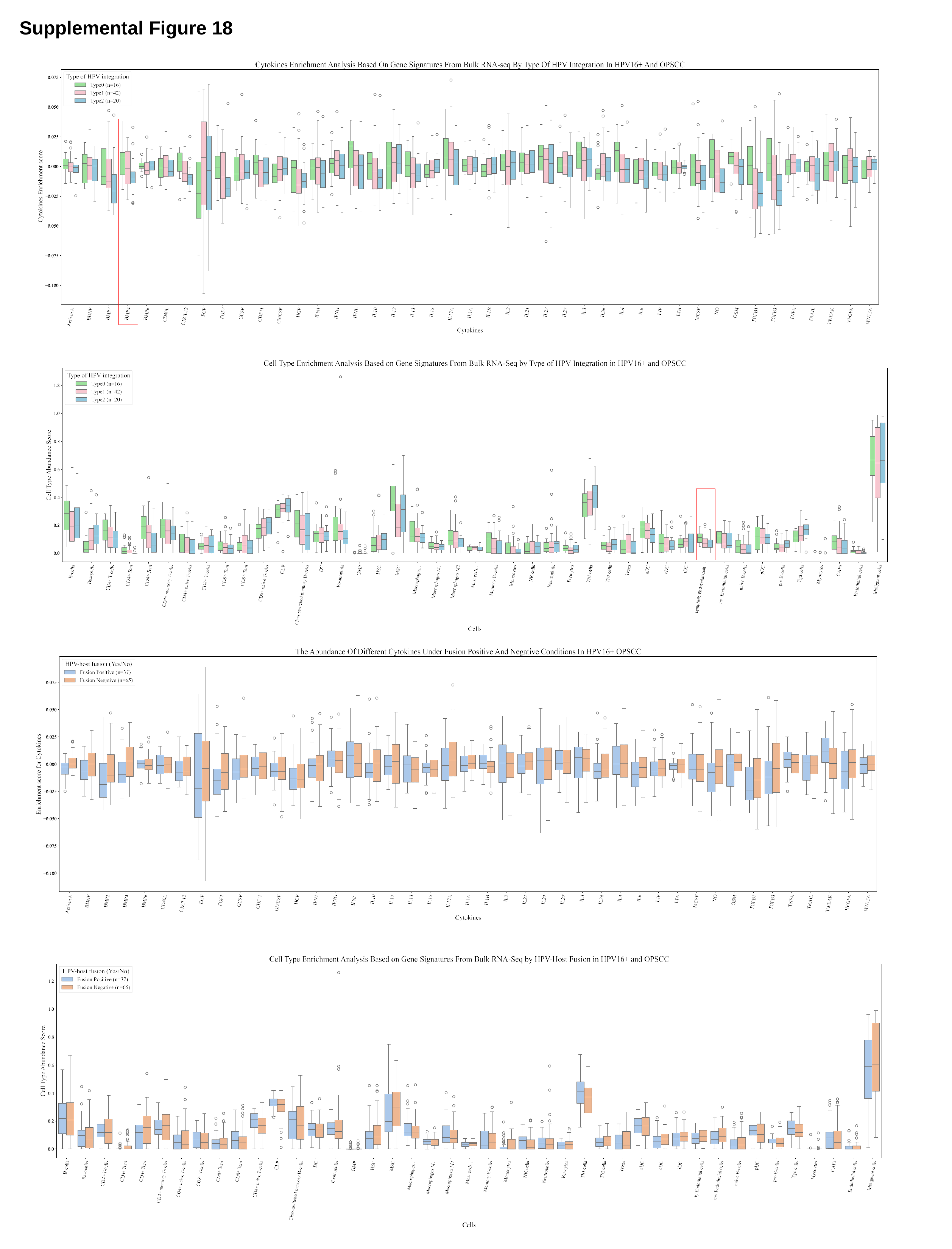

Supplemental Figure 18
Lymphatic Endothelial Cells

### Supplemental Figure 19

#### Slide 1
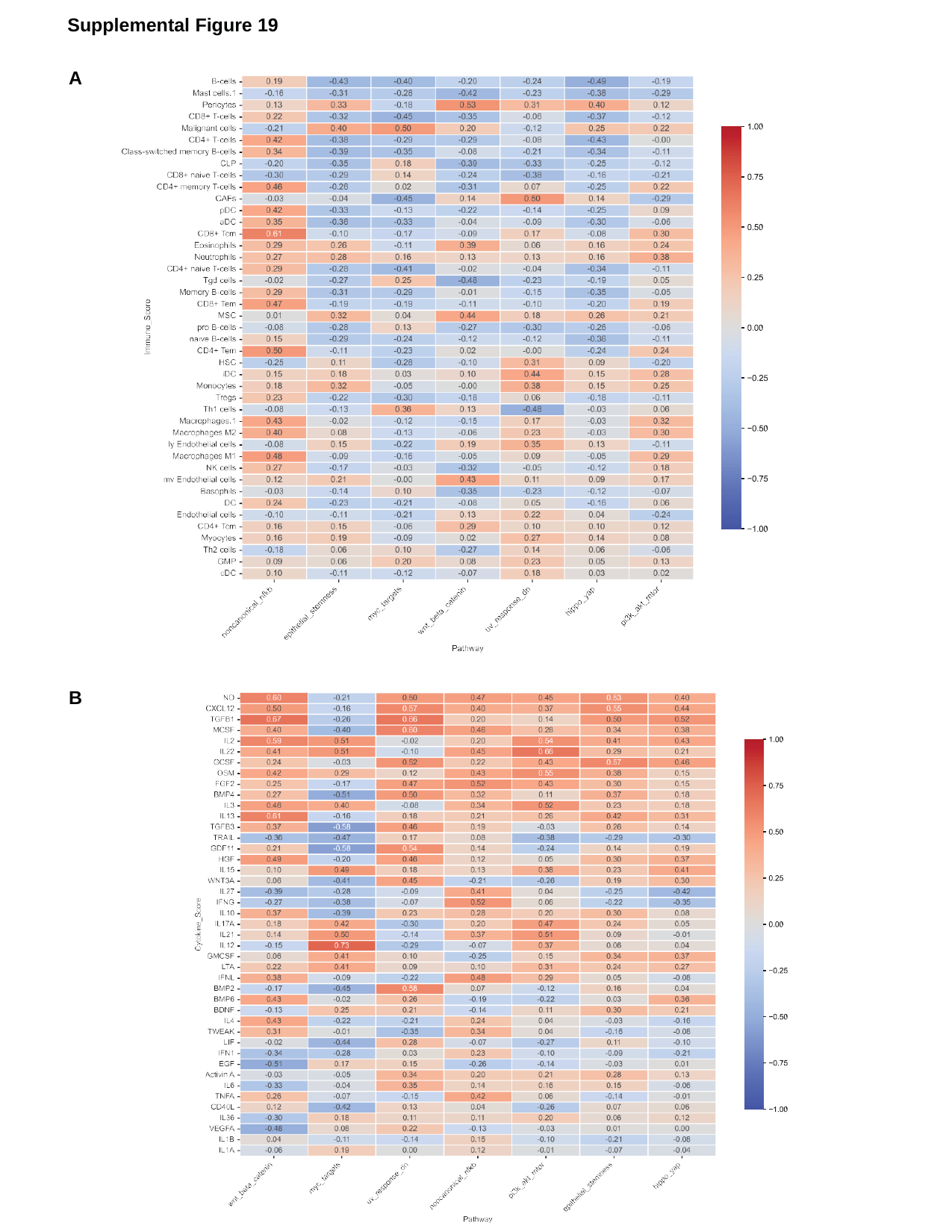

Supplemental Figure 19
A
B
