## Supplemental Figure 7 for "Spatiotemporal Heterogeneity of HPV Integration Architectures Drive Oncogenic Features of Oropharyngeal Cancer"

### Slide 1
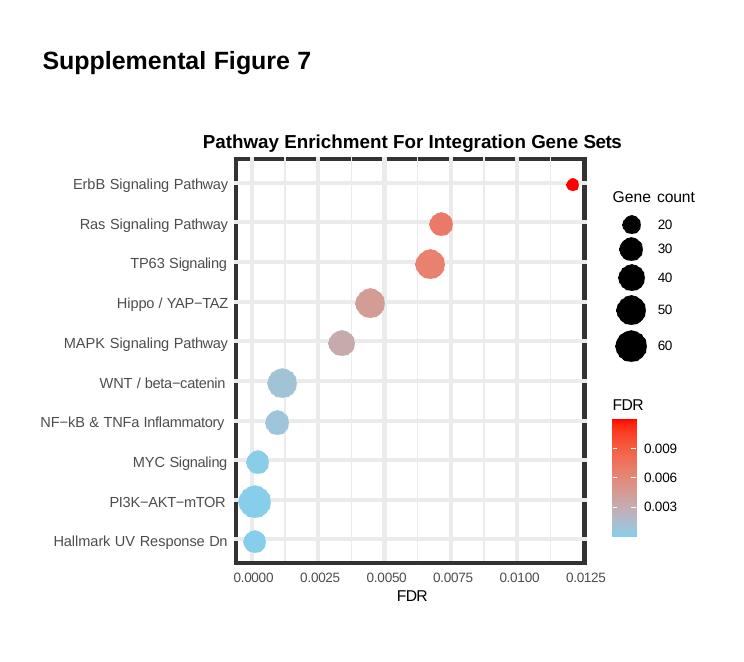

#### Supplemental Figure 7
Pathway Enrichment For Integration Gene Sets
ErbB Signaling Pathway
Gene count
20
30
Ras Signaling Pathway
TP63 Signaling
40
Hippo / YAP−TAZ
50
MAPK Signaling Pathway
60
WNT / beta−catenin
FDR
NF−kB & TNFa Inflammatory
0.009
MYC Signaling
0.006
PI3K−AKT−mTOR
0.003
Hallmark UV Response Dn
0.0000
0.0025
0.0050
FDR
0.0075
0.0100
0.0125
