## Supplemental Figure 8 for "Spatiotemporal Heterogeneity of HPV Integration Architectures Drive Oncogenic Features of Oropharyngeal Cancer"

### Slide 1
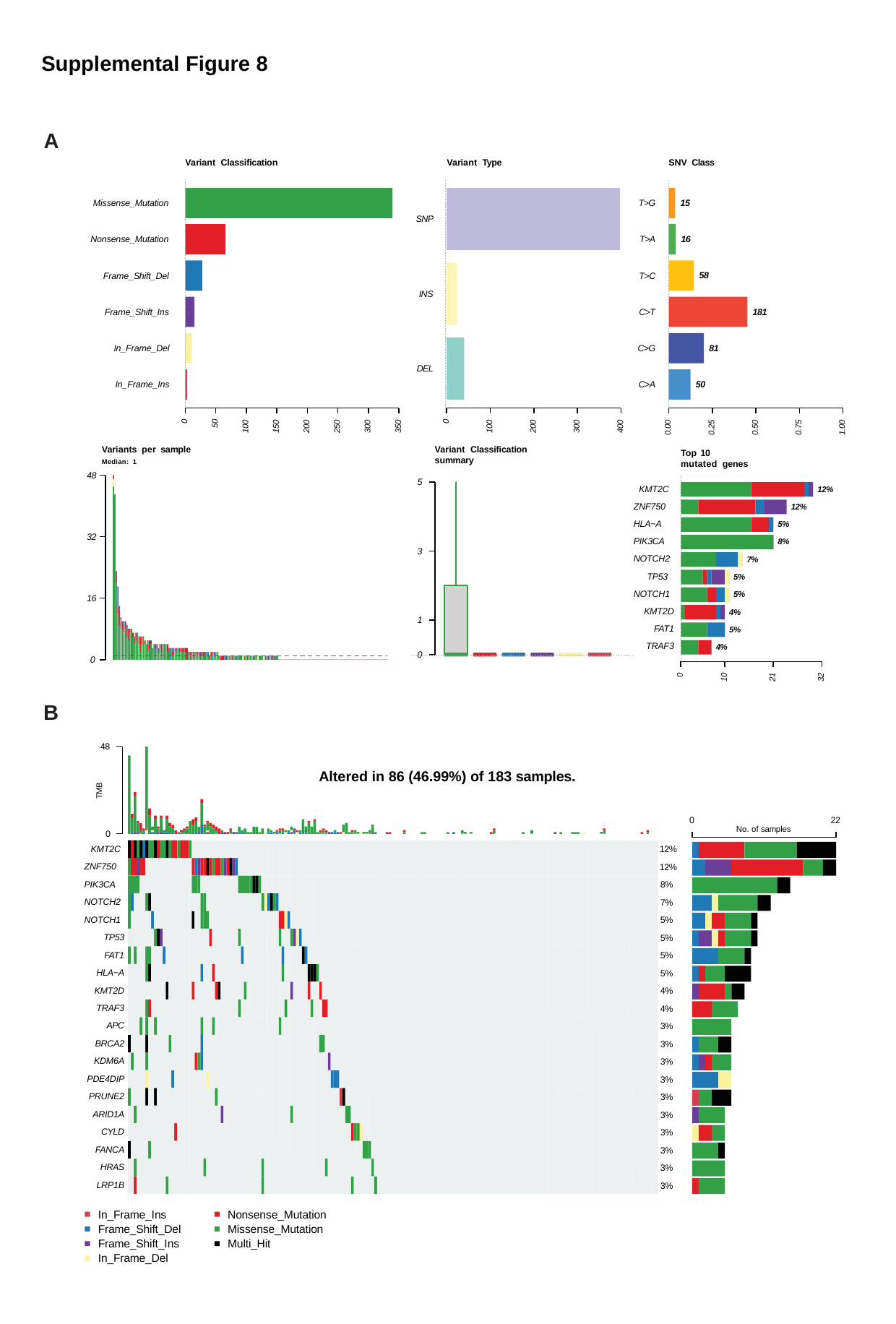

Supplemental Figure 8
A
Variant Classification
Variant Type
SNV Class
15
Missense_Mutation
T>G
SNP
16
Nonsense_Mutation
T>A
58
Frame_Shift_Del
T>C
INS
181
Frame_Shift_Ins
C>T
81
In_Frame_Del
C>G
DEL
50
In_Frame_Ins
C>A
100
200
300
400
0
0
50
100
150
200
250
300
350
0.00
0.25
0.50
0.75
1.00
Variants per sample
Median: 1
Variant Classification summary
Top 10 mutated genes
48
5
KMT2C ZNF750 HLA−A PIK3CA NOTCH2
TP53 NOTCH1
KMT2D FAT1 TRAF3
12%
12%
5%
8%
32
3
7%
5%
5%
16
4%
5%
1
4%
0
0
0
10
21
32
B
48
Altered in 86 (46.99%) of 183 samples.
TMB
0
22
No. of samples
0
KMT2C ZNF750 PIK3CA NOTCH2 NOTCH1
TP53 FAT1 HLA−A KMT2D TRAF3 APC BRCA2 KDM6A PDE4DIP PRUNE2 ARID1A CYLD FANCA HRAS LRP1B
12%
12%
8%
7%
5%
5%
5%
5%
4%
4%
3%
3%
3%
3%
3%
3%
3%
3%
3%
3%
In_Frame_Ins Frame_Shift_Del Frame_Shift_Ins In_Frame_Del
Nonsense_Mutation Missense_Mutation Multi_Hit
