## Supplemental Figure 11 for "Spatiotemporal Heterogeneity of HPV Integration Architectures Drive Oncogenic Features of Oropharyngeal Cancer"

### Slide 1
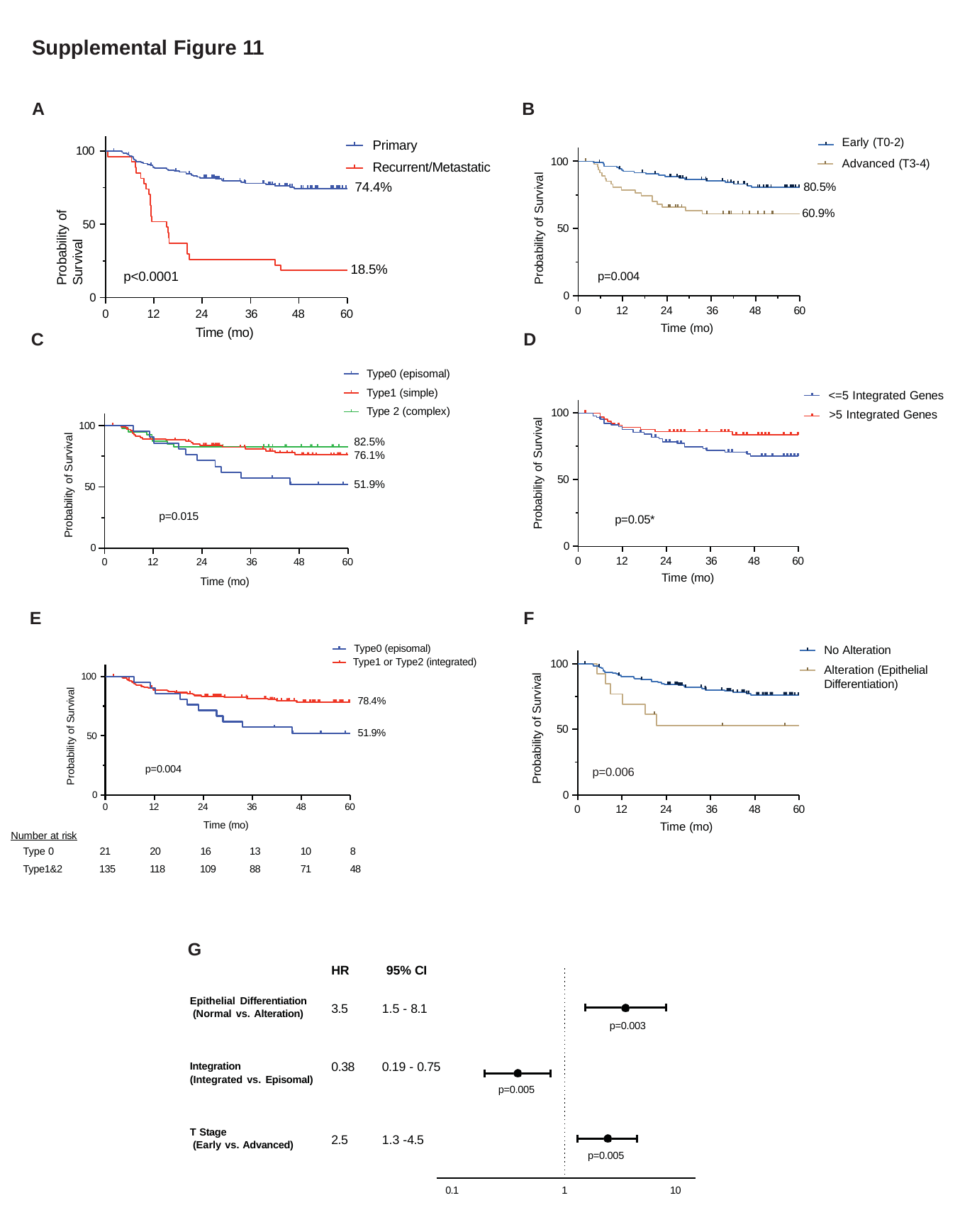

Supplemental Figure 11
A
B
Early (T0-2) Advanced (T3-4)
Primary Recurrent/Metastatic
74.4%
100
100
Probability of Survival
Probability of Survival
80.5%
60.9%
50
50
18.5%
p<0.0001
p=0.004
0
0
24	36
Time (mo)
24	36
Time (mo)
0
12
48
60
0
12
48
60
D
C
Type0 (episomal) Type1 (simple) Type 2 (complex)
<=5 Integrated Genes
>5 Integrated Genes
100
Probability of Survival
100
Probability of Survival
82.5%
76.1%
50
51.9%
50
p=0.015
p=0.05*
0
0
24	36
Time (mo)
24	36
Time (mo)
0
12
48
60
0
12
48
60
F
E
No Alteration
Alteration (Epithelial Differentiation)
Type0 (episomal)
Type1 or Type2 (integrated)
100
100
Probability of Survival
Probability of Survival
78.4%
50
51.9%
50
p=0.004
p=0.006
0
0
24	36
Time (mo)
0
12
24
36
48
60
0
12
48
60
Time (mo)
Number at risk
| Type 0 | 21 | 20 | 16 | 13 | 10 | 8 |
| --- | --- | --- | --- | --- | --- | --- |
| Type1&2 | 135 | 118 | 109 | 88 | 71 | 48 |
G
HR
95% CI
Epithelial Differentiation (Normal vs. Alteration)
3.5
1.5 - 8.1
p=0.003
0.38
0.19 - 0.75
Integration
(Integrated vs. Episomal)
p=0.005
T Stage
(Early vs. Advanced)
2.5
1.3 -4.5
p=0.005
0.1
1
10
