## Supplemental Figure 13 for "Spatiotemporal Heterogeneity of HPV Integration Architectures Drive Oncogenic Features of Oropharyngeal Cancer"

### Slide 1
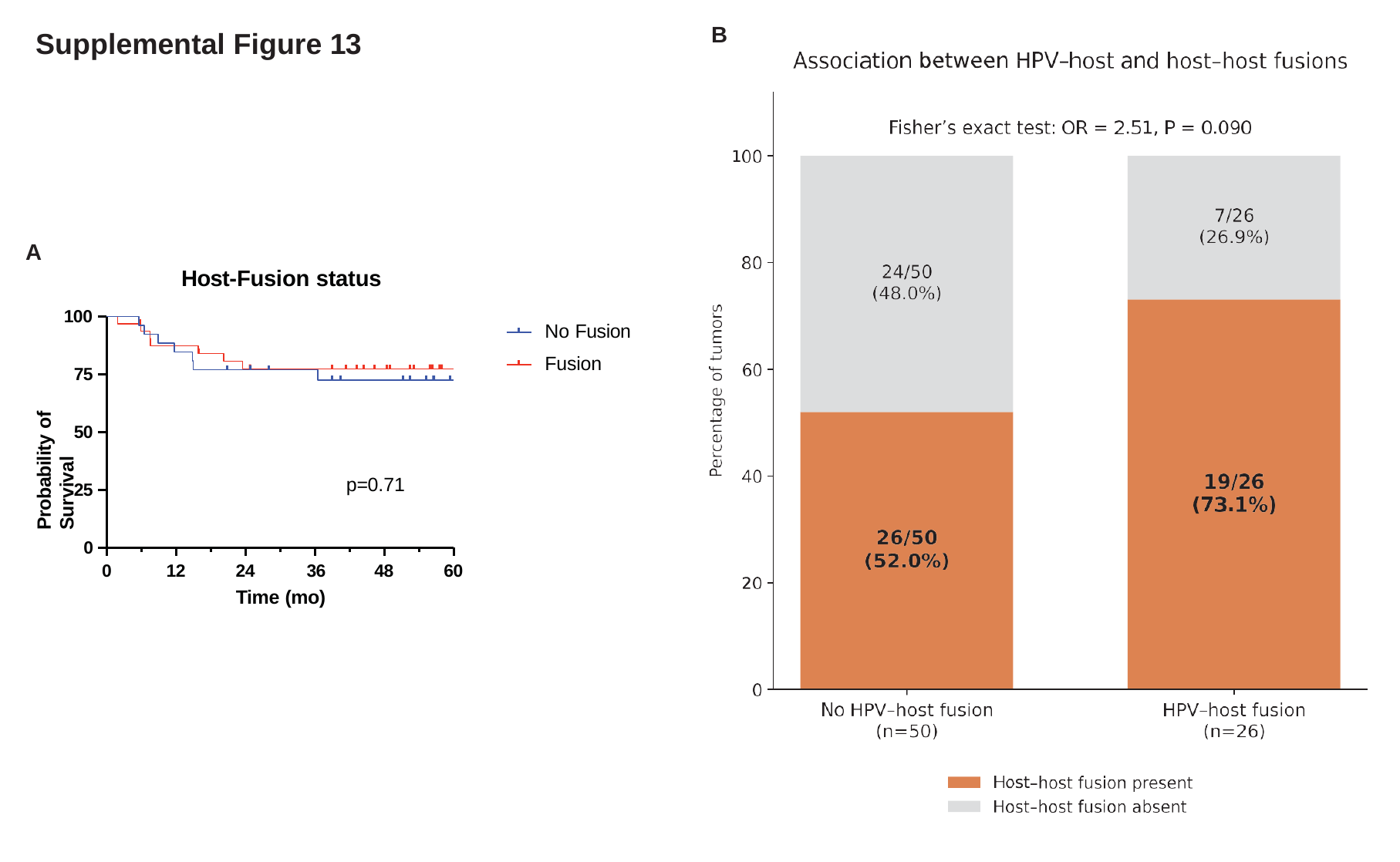

B
#### Supplemental Figure 13
A
Host-Fusion status
100
No Fusion Fusion
Probability of Survival
75
50
p=0.71
25
0
24	36
Time (mo)
0
12
48
60
