## Supplemental Figure 14 for "Spatiotemporal Heterogeneity of HPV Integration Architectures Drive Oncogenic Features of Oropharyngeal Cancer"

### Slide 1
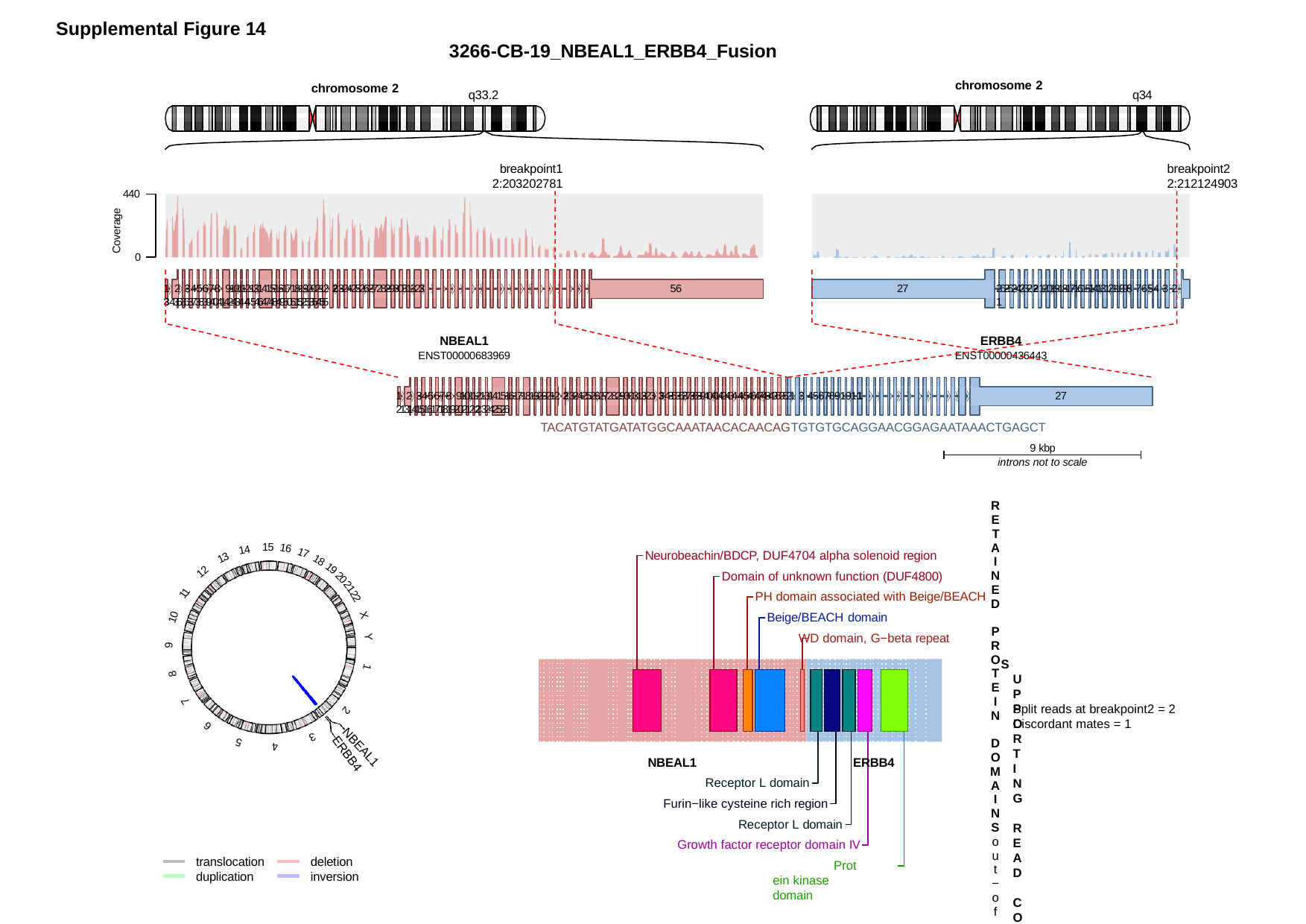

Supplemental Figure 14
3266-CB-19_NBEAL1_ERBB4_Fusion
chromosome 2
chromosome 2
q33.2
q34
breakpoint1 2:203202781
breakpoint2 2:212124903
440
Coverage
0
56
1 2 345678 9101121314151617181920212 232425262728293031323 3435363738394041424344546474849505152535455
27
2625242322120191817161514131211098 7654 3 2 1
NBEAL1
ENST00000683969
ERBB4
ENST00000436443
1 2 345678 9101121314151617181920212 232425262728293031323 343536373839404142434454647484950521 3 4567891011 2131415161718192021223242526
27
TACATGTATGATATGGCAAATAACACAACAGTGTGTGCAGGAACGGAGAATAAACTGAGCT
9 kbp
introns not to scale
RETAINED PROTEIN DOMAINS
out−of−frame fusion
15
16
14
Neurobeachin/BDCP, DUF4704 alpha solenoid region
17
13
18
Domain of unknown function (DUF4800)
PH domain associated with Beige/BEACH Beige/BEACH domain
WD domain, G−beta repeat
SUPPORTING READ COUNT
Split reads at breakpoint1 = 7
19
12
20
21
11
22
10
X
Y
9
1
8
7
Split reads at breakpoint2 = 2 Discordant mates = 1
2
6
3
5
4
NBEAL1
ERBB4
NBEAL1
ERBB4
Receptor L domain
Furin−like cysteine rich region
Receptor L domain Growth factor receptor domain IV
Protein kinase domain
translocation duplication
deletion inversion

### Slide 2
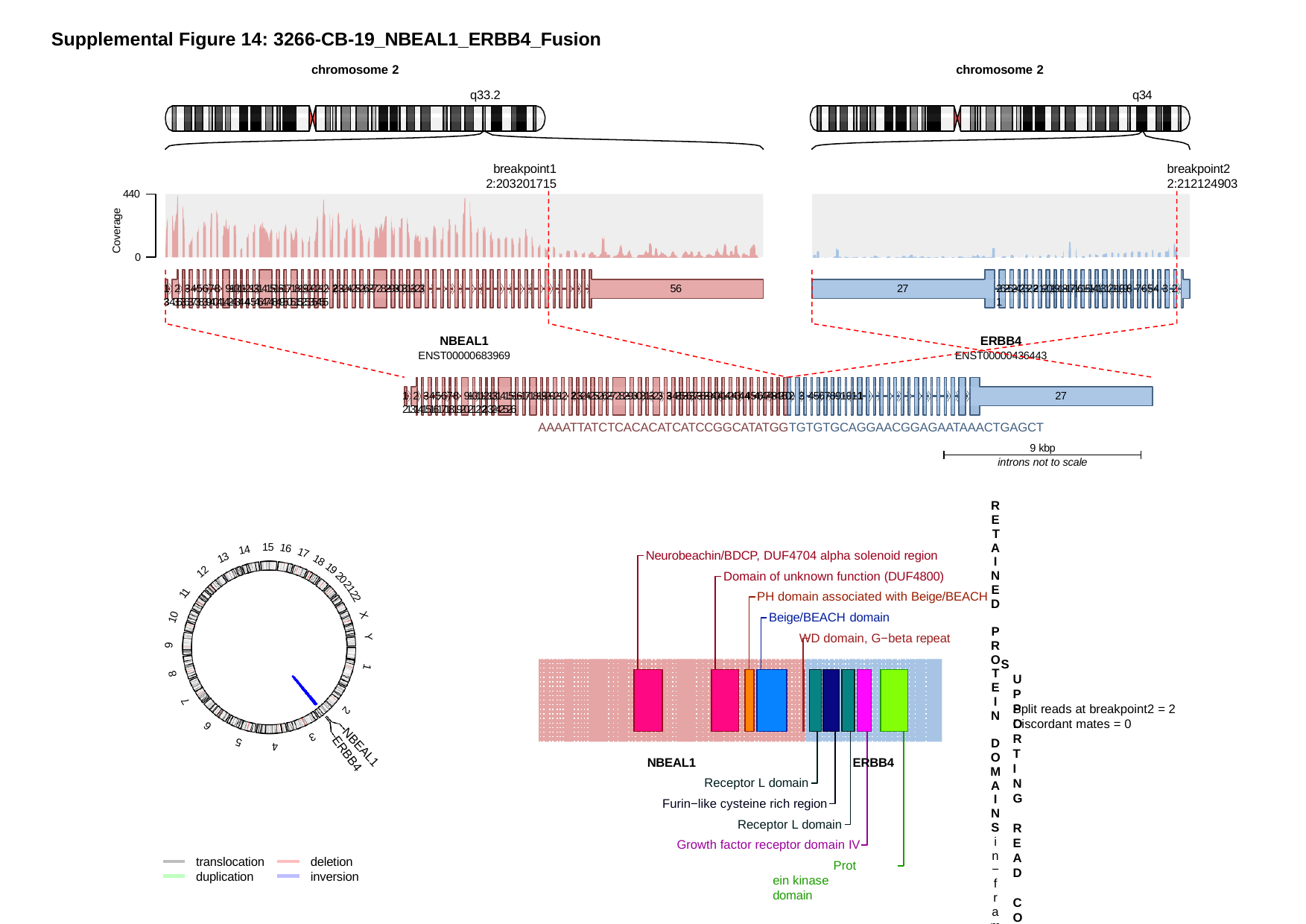

Supplemental Figure 14: 3266-CB-19_NBEAL1_ERBB4_Fusion
chromosome 2
q33.2
chromosome 2
q34
breakpoint1 2:203201715
breakpoint2 2:212124903
440
Coverage
0
56
1 2 345678 9101121314151617181920212 232425262728293031323 3435363738394041424344546474849505152535455
27
2625242322120191817161514131211098 7654 3 2 1
NBEAL1
ENST00000683969
ERBB4
ENST00000436443
1 2 345678 9101121314151617181920212 232425262728293031323 3435363738394041424344546474849502 3 4567891011 2131415161718192021223242526
27
AAAATTATCTCACACATCATCCGGCATATGGTGTGTGCAGGAACGGAGAATAAACTGAGCT
9 kbp
introns not to scale
RETAINED PROTEIN DOMAINS
in−frame fusion
15
16
14
Neurobeachin/BDCP, DUF4704 alpha solenoid region
17
13
18
Domain of unknown function (DUF4800)
PH domain associated with Beige/BEACH Beige/BEACH domain
WD domain, G−beta repeat
SUPPORTING READ COUNT
Split reads at breakpoint1 = 0
19
12
20
21
11
22
10
X
Y
9
1
8
7
Split reads at breakpoint2 = 2 Discordant mates = 0
2
6
3
5
4
NBEAL1
ERBB4
NBEAL1
ERBB4
Receptor L domain
Furin−like cysteine rich region
Receptor L domain Growth factor receptor domain IV
Protein kinase domain
translocation duplication
deletion inversion

### Slide 3
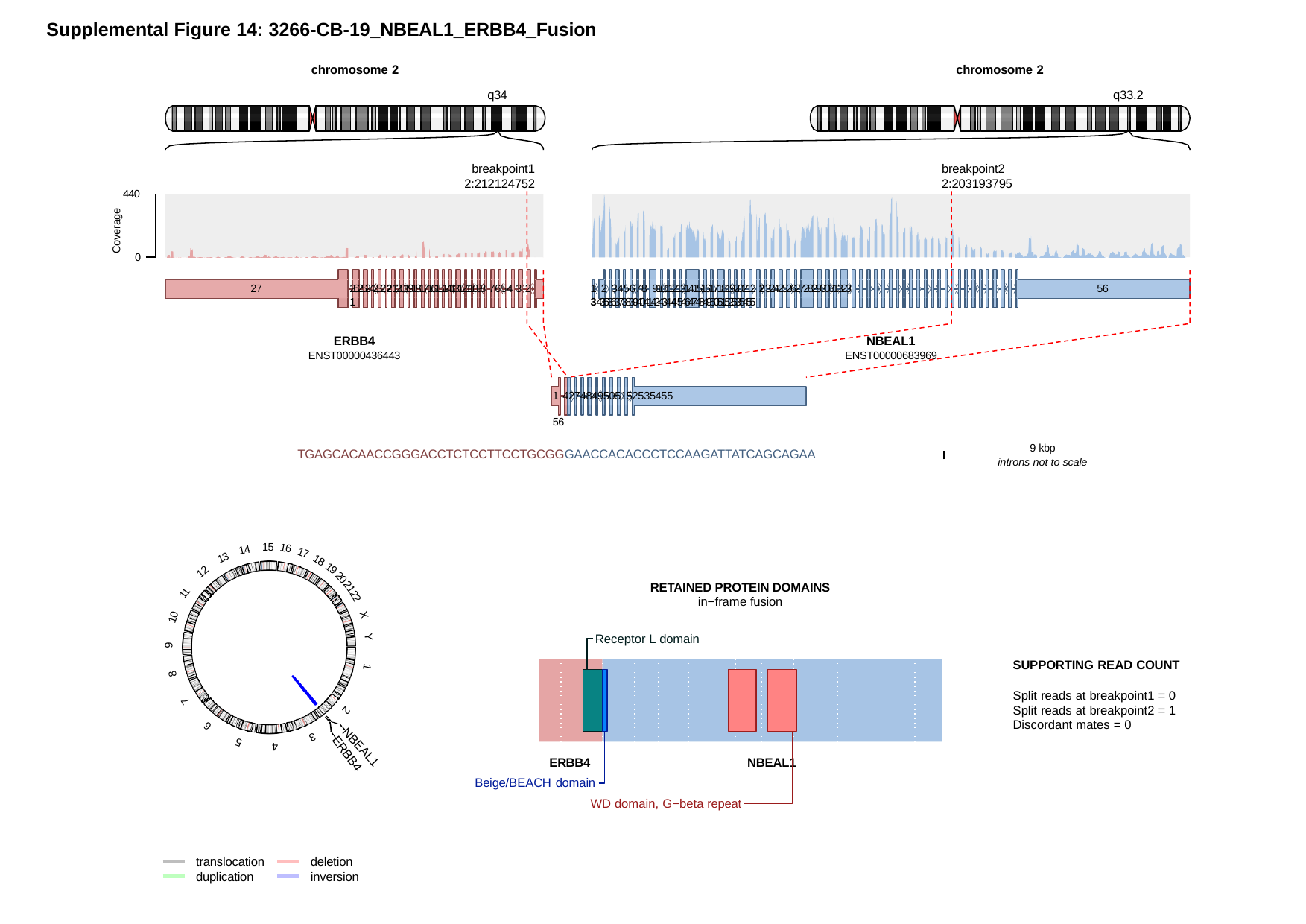

Supplemental Figure 14: 3266-CB-19_NBEAL1_ERBB4_Fusion
chromosome 2
chromosome 2
q34
q33.2
breakpoint1 2:212124752
breakpoint2 2:203193795
440
Coverage
0
56
27
2625242322120191817161514131211098 7654 3 2 1
1 2 345678 9101121314151617181920212 232425262728293031323 3435363738394041424344546474849505152535455
ERBB4
ENST00000436443
NBEAL1
ENST00000683969
1 4274849505152535455	56
TGAGCACAACCGGGACCTCTCCTTCCTGCGGGAACCACACCCTCCAAGATTATCAGCAGAA
9 kbp
introns not to scale
15
16
14
17
13
18
19
12
20
RETAINED PROTEIN DOMAINS
in−frame fusion
Receptor L domain
21
11
22
10
X
Y
9
SUPPORTING READ COUNT
Split reads at breakpoint1 = 0 Split reads at breakpoint2 = 1 Discordant mates = 0
1
8
7
2
6
3
5
4
NBEAL1
ERBB4
ERBB4
NBEAL1
Beige/BEACH domain
WD domain, G−beta repeat
translocation duplication
deletion inversion

### Slide 4
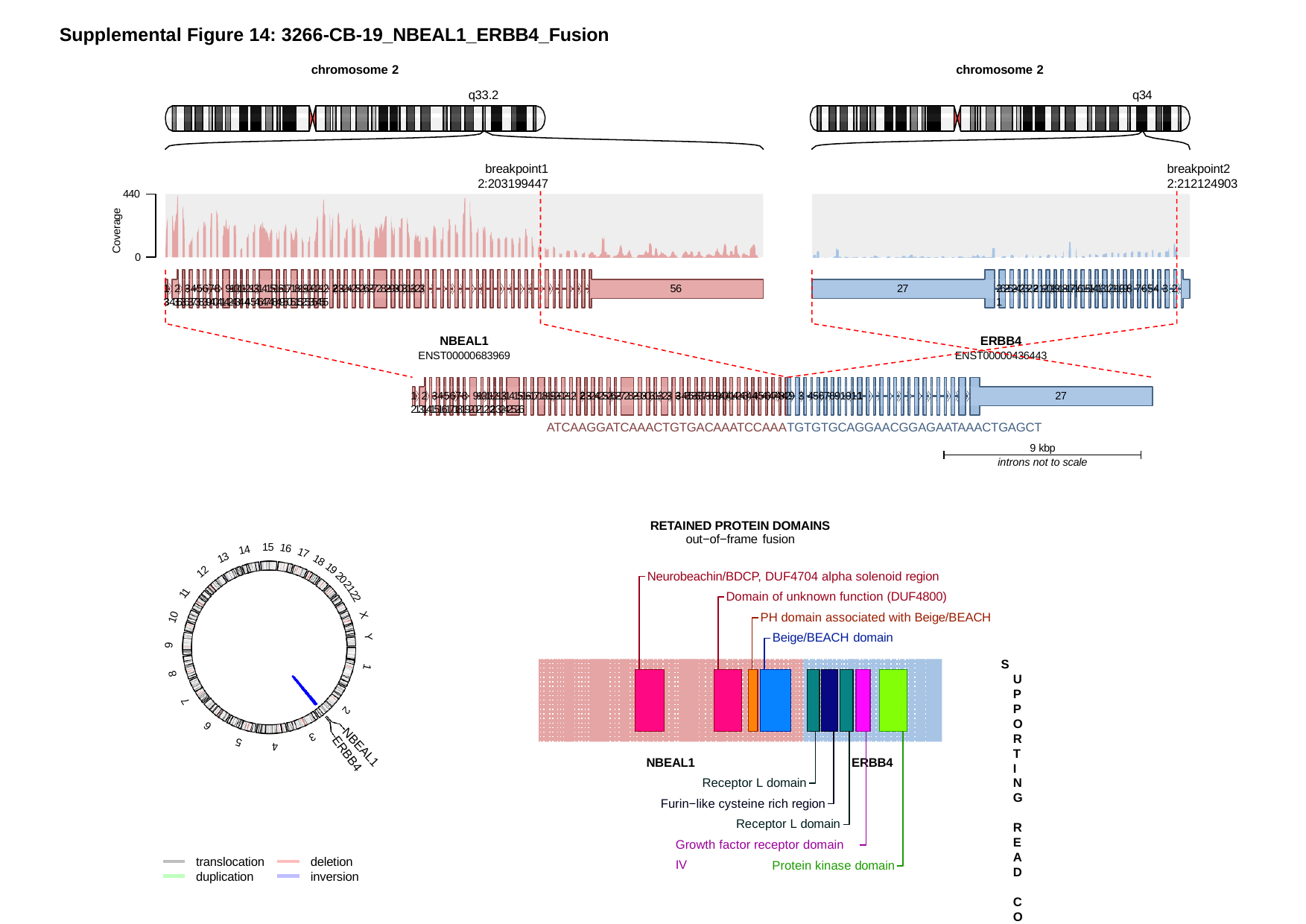

Supplemental Figure 14: 3266-CB-19_NBEAL1_ERBB4_Fusion
chromosome 2
chromosome 2
q33.2
q34
breakpoint1 2:203199447
breakpoint2 2:212124903
440
Coverage
0
56
1 2 345678 9101121314151617181920212 232425262728293031323 3435363738394041424344546474849505152535455
27
2625242322120191817161514131211098 7654 3 2 1
NBEAL1
ENST00000683969
ERBB4
ENST00000436443
1 2 345678 9101121314151617181920212 232425262728293031323 34353637383940414243445464748429 3 4567891011 2131415161718192021223242526
27
ATCAAGGATCAAACTGTGACAAATCCAAATGTGTGCAGGAACGGAGAATAAACTGAGCT
9 kbp
introns not to scale
RETAINED PROTEIN DOMAINS
out−of−frame fusion
15
16
14
17
13
18
Neurobeachin/BDCP, DUF4704 alpha solenoid region Domain of unknown function (DUF4800)
PH domain associated with Beige/BEACH Beige/BEACH domain
SUPPORTING READ COUNT
Split reads at breakpoint1 = 0 Split reads at breakpoint2 = 1 Discordant mates = 0
19
12
20
21
11
22
10
X
Y
9
1
8
7
2
6
3
5
4
NBEAL1
ERBB4
NBEAL1
ERBB4
Receptor L domain Furin−like cysteine rich region
Receptor L domain Growth factor receptor domain IV
translocation duplication
deletion inversion
Protein kinase domain

### Slide 5
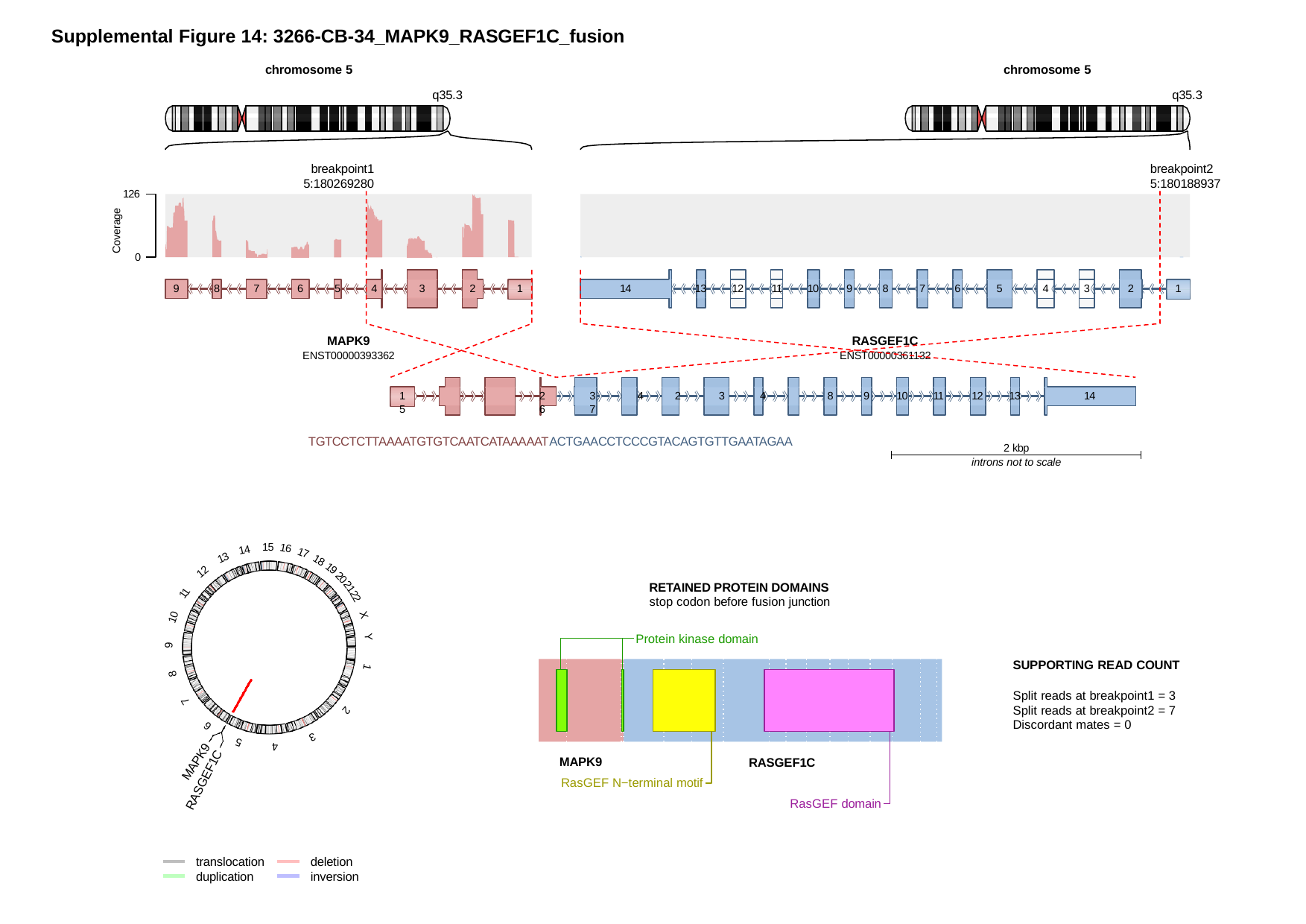

Supplemental Figure 14: 3266-CB-34_MAPK9_RASGEF1C_fusion
chromosome 5
q35.3
chromosome 5
q35.3
breakpoint1 5:180269280
breakpoint2 5:180188937
126
Coverage
0
12
11
4
3
5
5
9
8
6
3
13
10
9
8
6
7
7
4
2
1
14
2
1
MAPK9
ENST00000393362
RASGEF1C
ENST00000361132
8
9
10
13
1	2	3	4	2	3	4	5	6	7
TGTCCTCTTAAAATGTGTCAATCATAAAAATACTGAACCTCCCGTACAGTGTTGAATAGAA
11
12
14
2 kbp
introns not to scale
15
16
14
17
13
18
19
12
20
RETAINED PROTEIN DOMAINS
stop codon before fusion junction
Protein kinase domain
21
11
22
10
X
Y
9
SUPPORTING READ COUNT
Split reads at breakpoint1 = 3 Split reads at breakpoint2 = 7 Discordant mates = 0
1
8
7
2
6
3
5
4
MAPK9
RASGEF1C
MAPK9
RASGEF1C
RasGEF N−terminal motif
RasGEF domain
translocation duplication
deletion inversion

### Slide 6
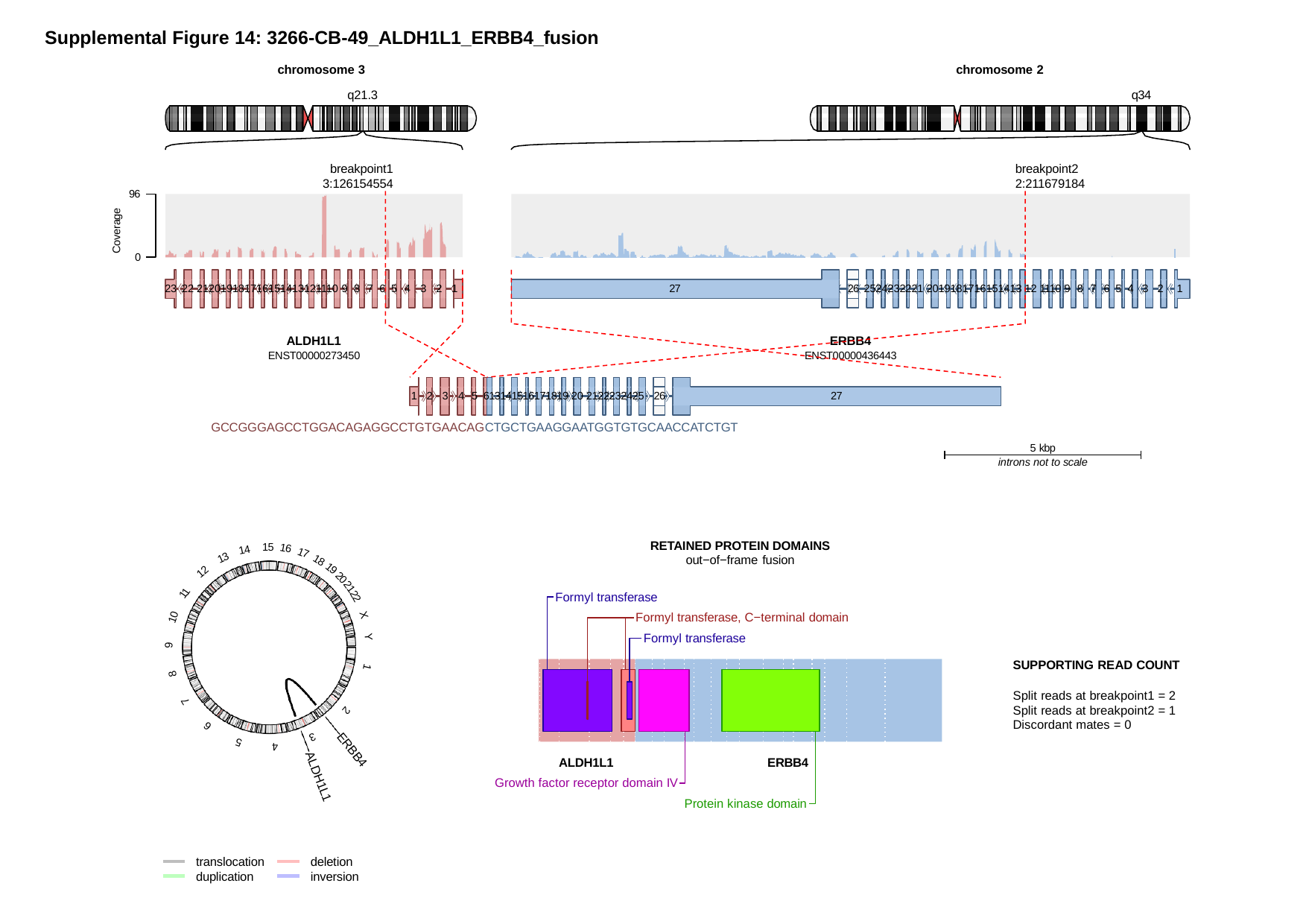

Supplemental Figure 14: 3266-CB-49_ALDH1L1_ERBB4_fusion
chromosome 3
q21.3
chromosome 2
q34
breakpoint1 3:126154554
breakpoint2 2:211679184
96
Coverage
0
26
23 22 212019181716151413121110 9 8 7 6 5 4 3 2 1
27
2524232221 2019181716151413 12 1110 9 8 7 6 5 4 3 2 1
ALDH1L1
ENST00000273450
ERBB4
ENST00000436443
26
1 2 3 4 5 613141516171819 20 2122232425
27
GCCGGGAGCCTGGACAGAGGCCTGTGAACAGCTGCTGAAGGAATGGTGTGCAACCATCTGT
5 kbp
introns not to scale
RETAINED PROTEIN DOMAINS
out−of−frame fusion
15
16
14
17
13
18
19
12
20
Formyl transferase
Formyl transferase, C−terminal domain Formyl transferase
21
11
22
10
X
Y
9
SUPPORTING READ COUNT
Split reads at breakpoint1 = 2 Split reads at breakpoint2 = 1 Discordant mates = 0
1
8
7
2
6
3
5
4
ERBB4
ALDH1L1
ERBB4
Growth factor receptor domain IV
Protein kinase domain
ALDH1L1
translocation duplication
deletion inversion

### Slide 7
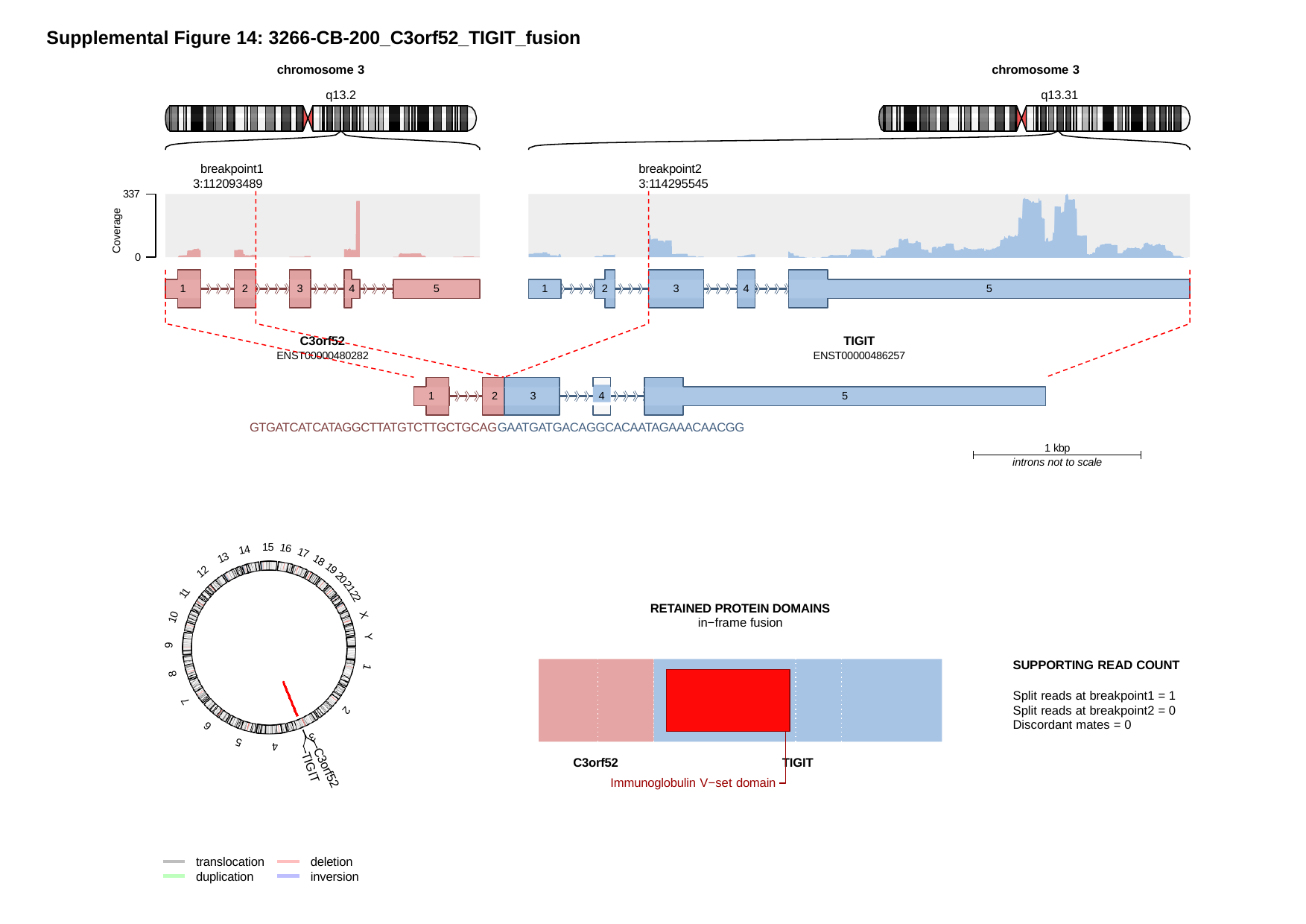

Supplemental Figure 14: 3266-CB-200_C3orf52_TIGIT_fusion
chromosome 3
q13.2
chromosome 3
q13.31
breakpoint1 3:112093489
breakpoint2 3:114295545
337
Coverage
0
5
5
3
3
1
2
4
1
2
4
C3orf52
ENST00000480282
TIGIT
ENST00000486257
4
5
1	2	3
GTGATCATCATAGGCTTATGTCTTGCTGCAGGAATGATGACAGGCACAATAGAAACAACGG
1 kbp
introns not to scale
15
16
14
17
13
18
19
12
20
21
11
22
RETAINED PROTEIN DOMAINS
in−frame fusion
10
X
Y
9
SUPPORTING READ COUNT
Split reads at breakpoint1 = 1 Split reads at breakpoint2 = 0 Discordant mates = 0
1
8
7
2
6
3
5
4
C3orf52
TIGIT
TIGIT
C3orf52
Immunoglobulin V−set domain
translocation duplication
deletion inversion

### Slide 8
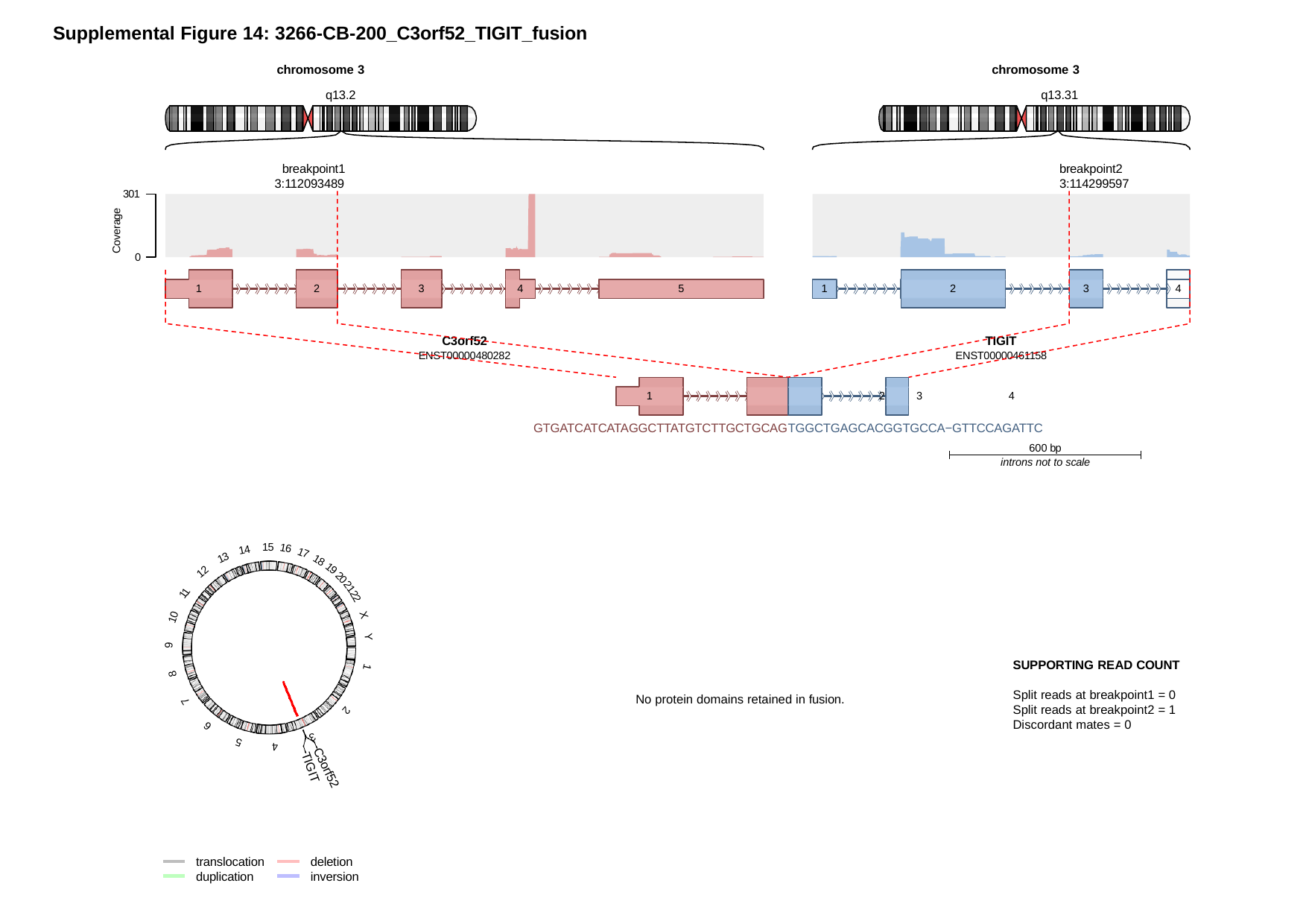

Supplemental Figure 14: 3266-CB-200_C3orf52_TIGIT_fusion
chromosome 3
q13.2
chromosome 3
q13.31
breakpoint1 3:112093489
breakpoint2 3:114299597
301
Coverage
0
4
5
3
3
1
2
4
1
2
C3orf52
ENST00000480282
TIGIT
ENST00000461158
1	2	3	4
GTGATCATCATAGGCTTATGTCTTGCTGCAGTGGCTGAGCACGGTGCCA−GTTCCAGATTC
600 bp
introns not to scale
15
16
14
17
13
18
19
12
20
21
11
22
10
X
Y
9
SUPPORTING READ COUNT
1
8
Split reads at breakpoint1 = 0 Split reads at breakpoint2 = 1 Discordant mates = 0
No protein domains retained in fusion.
7
2
6
3
5
4
TIGIT
C3orf52
translocation duplication
deletion inversion

### Slide 9
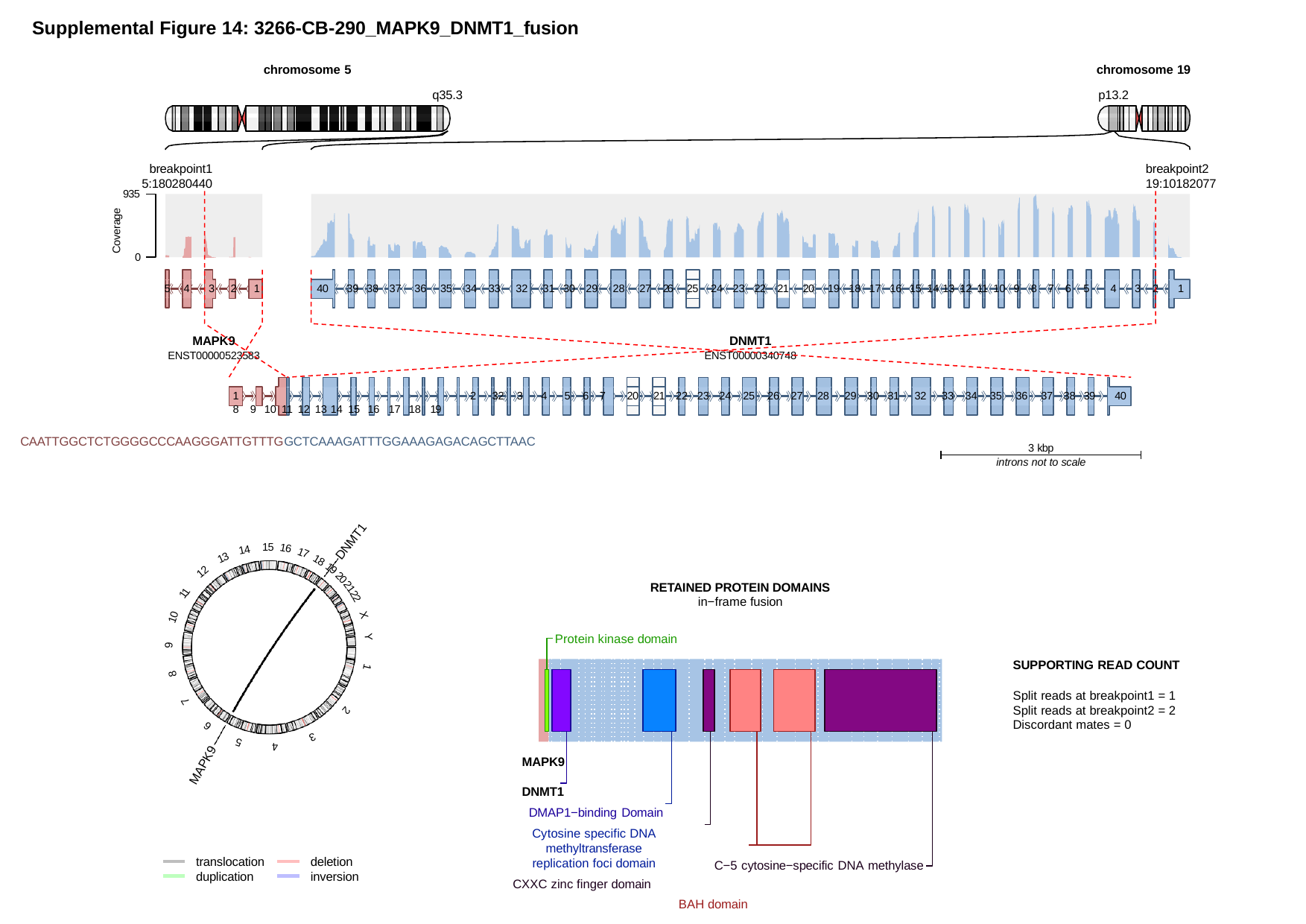

Supplemental Figure 14: 3266-CB-290_MAPK9_DNMT1_fusion
chromosome 5
chromosome 19
p13.2
q35.3
breakpoint1 5:180280440
breakpoint2 19:10182077
935
Coverage
0
25
21
20
5 4	3	2	1
40	39 38 37 36 35 34 33 32 31 30 29 28 27 26
24 23 22
19 18 17 16 15 14 13 12 11 10 9 8 7 6 5	4	3 2	1
MAPK9
ENST00000523583
DNMT1
ENST00000340748
1	2	32 3	4	5 6 7 8 9 10 11 12 13 14 15 16 17 18 19
CAATTGGCTCTGGGGCCCAAGGGATTGTTTGGCTCAAAGATTTGGAAAGAGACAGCTTAAC
22 23 24 25 26 27 28 29 30 31 32 33 34 35 36 37 38 39	40
20
21
3 kbp
introns not to scale
DNMT1
15
16
14
17
13
18
19
12
20
RETAINED PROTEIN DOMAINS
in−frame fusion
Protein kinase domain
21
11
22
10
X
Y
9
SUPPORTING READ COUNT
Split reads at breakpoint1 = 1 Split reads at breakpoint2 = 2 Discordant mates = 0
1
8
7
2
6
3
5
4
MAPK9	DNMT1
DMAP1−binding Domain
Cytosine specific DNA methyltransferase replication foci domain
CXXC zinc finger domain
BAH domain
MAPK9
translocation duplication
deletion inversion
C−5 cytosine−specific DNA methylase

### Slide 10
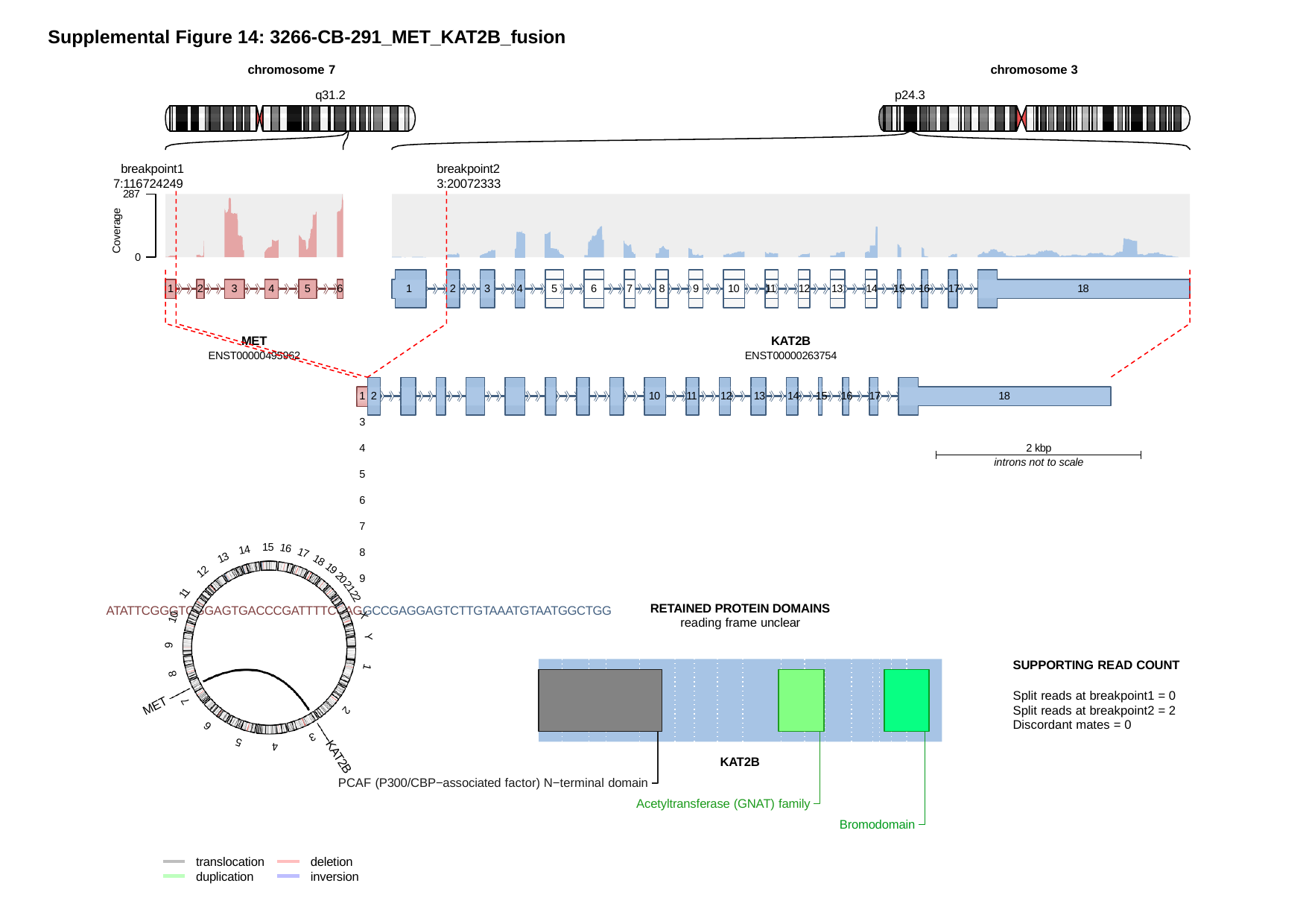

Supplemental Figure 14: 3266-CB-291_MET_KAT2B_fusion
chromosome 7
q31.2
chromosome 3
p24.3
breakpoint1
breakpoint2 3:20072333
7:116724249
287
Coverage
0
5
6
7
8
9
10
11
12
13
14
5
3
6
3
18
1
2
4
1
2
4
15 16	17
MET
ENST00000495962
KAT2B
ENST00000263754
10
18
1 2	3	4	5	6	7	8	9
ATATTCGGGTGGGAGTGACCCGATTTTCCAGGCCGAGGAGTCTTGTAAATGTAATGGCTGG
11	12	13	14	15 16	17
2 kbp
introns not to scale
15
16
14
17
13
18
19
12
20
21
11
22
RETAINED PROTEIN DOMAINS
reading frame unclear
10
X
Y
9
SUPPORTING READ COUNT
Split reads at breakpoint1 = 0 Split reads at breakpoint2 = 2 Discordant mates = 0
1
8
7
MET
2
6
3
5
4
KAT2B
PCAF (P300/CBP−associated factor) N−terminal domain
Acetyltransferase (GNAT) family
Bromodomain
KAT2B
translocation duplication
deletion inversion
