## Supplemental Figure 15 for "Spatiotemporal Heterogeneity of HPV Integration Architectures Drive Oncogenic Features of Oropharyngeal Cancer"

### Slide 1
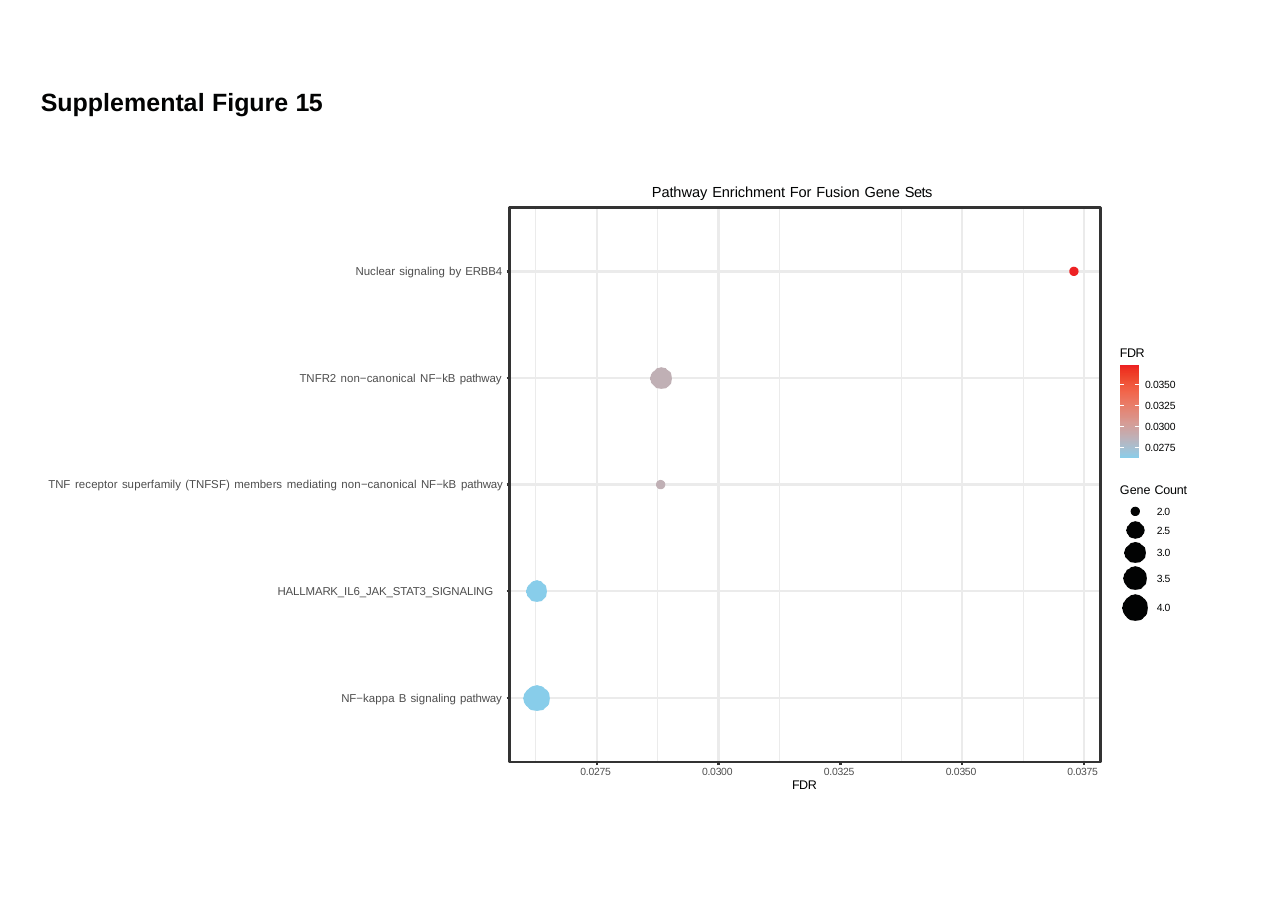

#### Supplemental Figure 15
Pathway Enrichment For Fusion Gene Sets
Nuclear signaling by ERBB4
FDR
TNFR2 non−canonical NF−kB pathway
0.0350
0.0325
0.0300
0.0275
TNF receptor superfamily (TNFSF) members mediating non−canonical NF−kB pathway
Gene Count
2.0
2.5
3.0
3.5
HALLMARK_IL6_JAK_STAT3_SIGNALING
4.0
NF−kappa B signaling pathway
0.0275
0.0300
0.0325
0.0350
0.0375
FDR
