## Supplemental Figure 16 for "Spatiotemporal Heterogeneity of HPV Integration Architectures Drive Oncogenic Features of Oropharyngeal Cancer"

### Slide 1
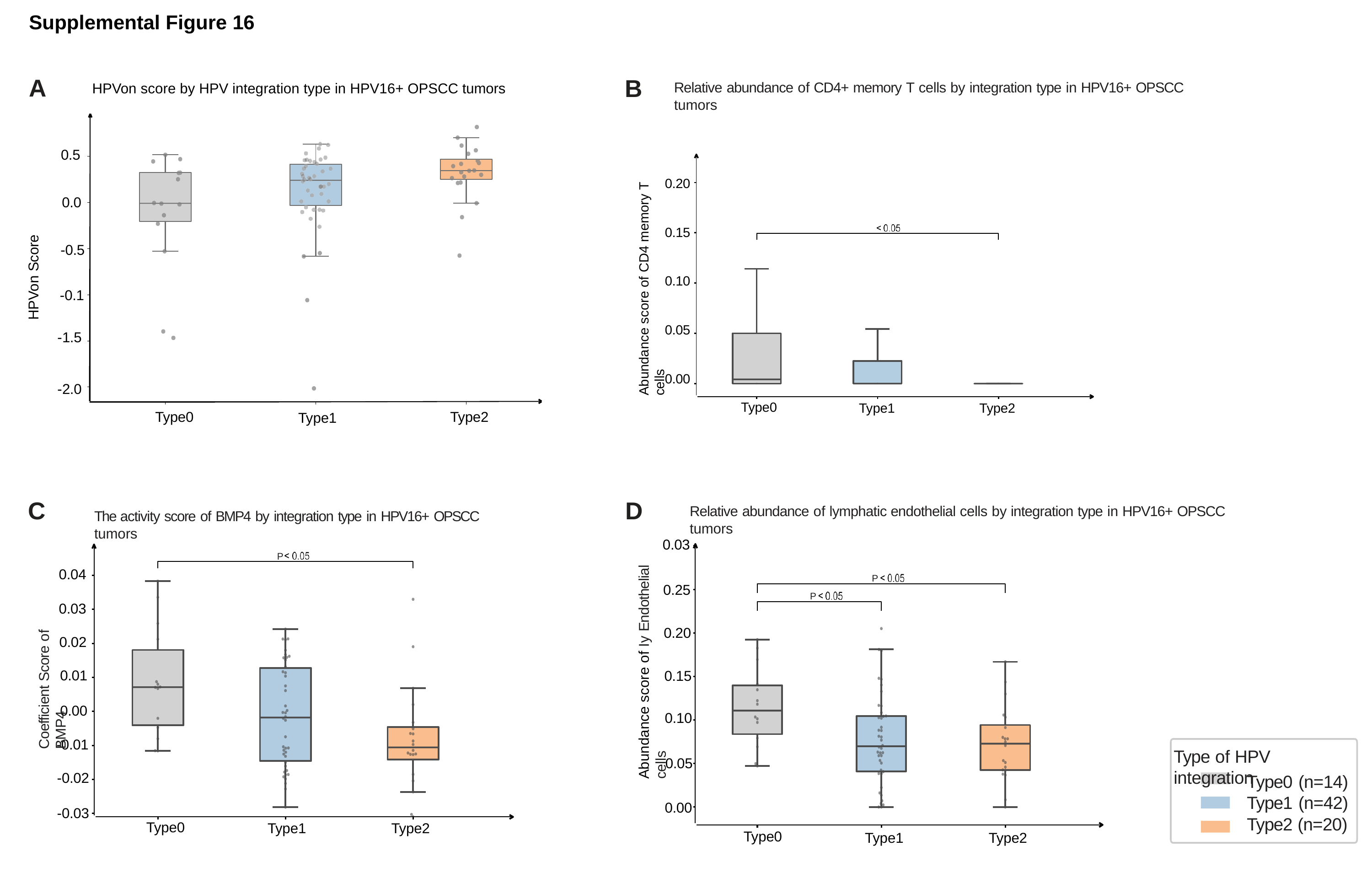

Supplemental Figure 16
A
B
Relative abundance of CD4+ memory T cells by integration type in HPV16+ OPSCC tumors
HPVon score by HPV integration type in HPV16+ OPSCC tumors
0.5
Abundance score of CD4 memory T cells
0.20
0.0
P
0.15
HPVon Score
-0.5
0.10
-0.1
0.05
-1.5
0.00
-2.0
Type0
Type1
Type2
Type0
Type2
Type1
C
D
Relative abundance of lymphatic endothelial cells by integration type in HPV16+ OPSCC tumors
The activity score of BMP4 by integration type in HPV16+ OPSCC tumors
Abundance score of Iy Endothelial cells
0.03
P
0.04
P
0.25
Coefficient Score of BMP4
P
0.03
0.20
0.02
0.01
0.15
0.00
0.10
-0.01
Type of HPV integration
0.05
-0.02
Type0 (n=14) Type1 (n=42) Type2 (n=20)
0.00
-0.03
Type0
Type1
Type2
Type0
Type1
Type2
