## Supplemental Figure 17 for "Spatiotemporal Heterogeneity of HPV Integration Architectures Drive Oncogenic Features of Oropharyngeal Cancer"

### Slide 1
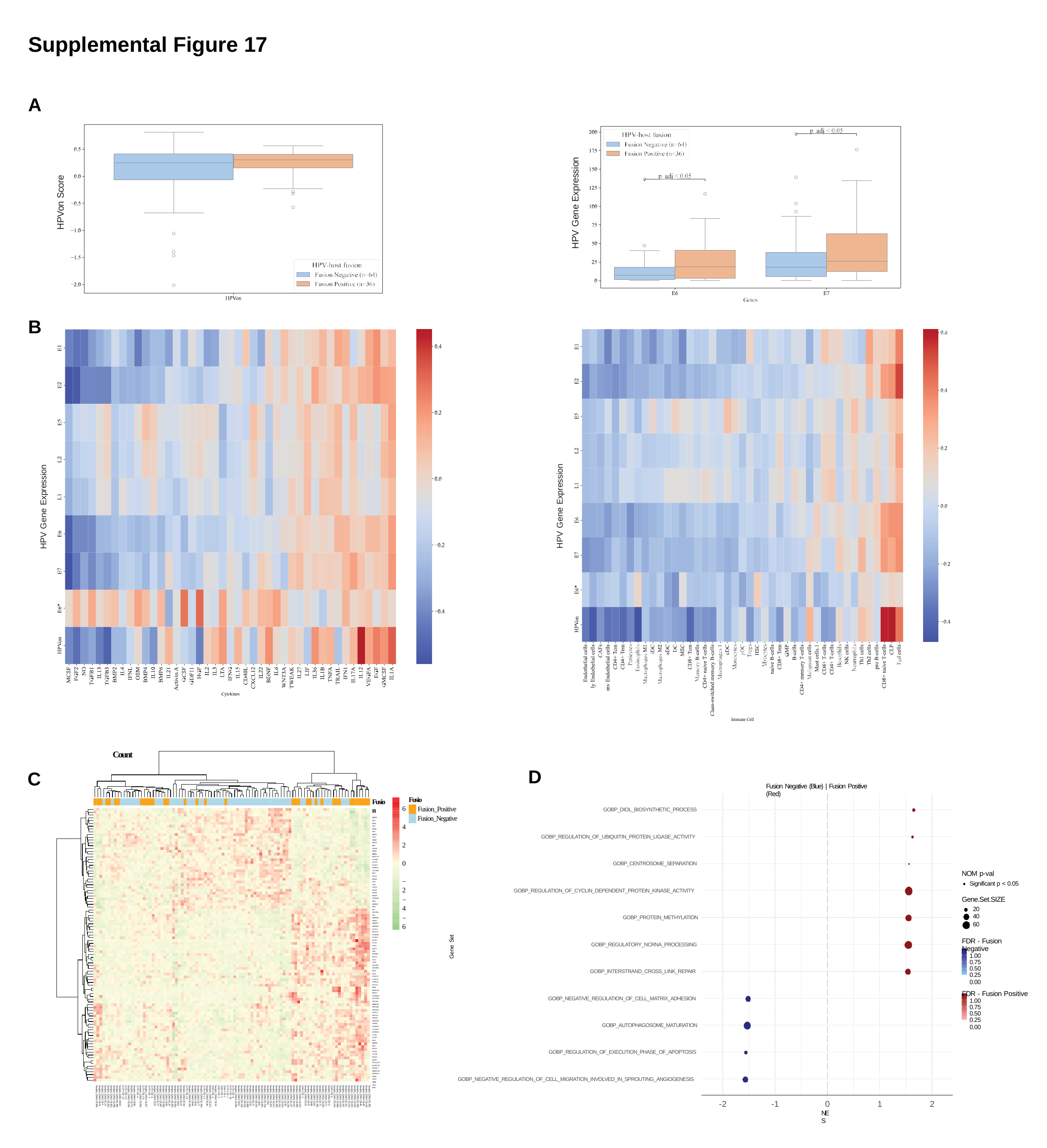

Supplemental Figure 17
A
HPV Gene Expression
HPVon Score
B
HPV Gene Expression
HPV Gene Expression
Significant Genes in Fusion Positive vs Fusion Negative with Normalized Count
D
C
Fusion Negative (Blue) | Fusion Positive (Red)
Fusion
Fusion
KRT17P3 PLS3 RND3 SAT1 PTPRH SLPI GPRC5A CD55 TIPARP DIO3OS MN1 CCDC80 DPYSL3 RPS6KA2
IGHV5−10−1 LONRF2 SCN3B KCND3 HSPB1P1 TAX1BP3 BNC1
MAOA PLSCR4 CHL1 VAV3 CHST9 FOXP2 COL21A1 SMG1P5
AC022149.1 NEFL KBTBD12 P3H2
DST TBC1D8B WLS AP001029.2 SPIRE1 AP001057.1 AP001056.2 ICOSLG MEGF10 AC110760.2 GVQW3 SFXN2 SYCP3 YPEL1 KRT32
SIM1 ZBTB8B KRT36 OCA2 GAD1 AC013562.1 AC022893.2 MYB
DLX6 GPR137C CLDN16 TCERG1L ELOVL7 XKR9 DEPDC1B MEIOC AC024610.2 AC123769.1 TSPEAR TSPEAR−AS2 DPY19L2P1 LINC02613 GLB1L2 SAPCD2 NEURL1 TMEM201 WDR62 AC091057.1 TCAM1P AL590226.2 CCNI2 DGAT2
FZD7 LDLRAD3 ATF5 KLRG2 KCNJ6
C2orf91 FHAD1 SPESP1 FP565260.3 KLHDC7B DNAH17 DNAH17−AS1 AP000525.1 TRIB3
GSTO2 GATD3A CBX2 ESRG
Fusion_Positive Fusion_Negative
6
GOBP_DIOL_BIOSYNTHETIC_PROCESS
4
GOBP_REGULATION_OF_UBIQUITIN_PROTEIN_LIGASE_ACTIVITY
2
0
GOBP_CENTROSOME_SEPARATION
NOM p-val
Significant p < 0.05
−2
GOBP_REGULATION_OF_CYCLIN_DEPENDENT_PROTEIN_KINASE_ACTIVITY
Gene.Set.SIZE
20
40
60
−4
−6
GOBP_PROTEIN_METHYLATION
Gene Set
FDR - Fusion Negative
1.00
0.75
0.50
0.25
0.00
FDR - Fusion Positive
1.00
0.75
0.50
0.25
0.00
GOBP_REGULATORY_NCRNA_PROCESSING
GOBP_INTERSTRAND_CROSS_LINK_REPAIR
GOBP_NEGATIVE_REGULATION_OF_CELL_MATRIX_ADHESION
GOBP_AUTOPHAGOSOME_MATURATION
GOBP_REGULATION_OF_EXECUTION_PHASE_OF_APOPTOSIS
GOBP_NEGATIVE_REGULATION_OF_CELL_MIGRATION_INVOLVED_IN_SPROUTING_ANGIOGENESIS
Sample_3266.CB.385 Sample_3266.CB.146 Sample_3266.CB.49 Sample_3266.CB.46 Sample_3266.CB.268 Sample_3266.CB.183 Sample_3266.CB.182 Sample_3266.CB.314 Sample_3266.CB.311 Sample_3266.CB.297 Sample_3266.CB.103 Sample_3266.CB.386 Sample_3266.CB.316 5343−CB−6
Sample_3266.CB.51
Sample_3266.CB.72 Sample_3266.CB.259 Sample_3266.CB.75 Sample_3266.CB.392 Sample_3266.CB.93 Sample_3266.CB.89 Sample_3266.CB.47 Sample_3266.CB.65 5343−CB−5
Sample_3266.CB.102
Sample_3266.CB.309 Sample_3266.CB.107 Sample_3266.CB.237 Sample_3266.CB.236 Sample_3266.CB.39 Sample_3266.CB.318 Sample_3266.CB.292 Sample_3266.CB.290 Sample_3266.CB.313 Sample_3266.CB.291 Sample_3266.CB.6 Sample_3266.CB.104 Sample_3266.CB.285 Sample_3266.CB.61 Sample_3266.CB.319 Sample_3266.CB.27 Sample_3266.CB.50 Sample_3266.CB.388 Sample_3266.CB.109 Sample_3266.CB.108 Sample_3266.CB.63 Sample_3266.CB.296 5170−CB−8
5170−CB−20
5170−CB−1
5343−CB−1
5343−CB−3
5343−CB−2
Sample_3266.CB.34 5343−CB−4
Sample_3266.CB.315
Sample_3266.CB.86 5170−CB−12
Sample_3266.CB.394
Sample_3266.CB.77 Sample_3266.CB.95 Sample_3266.CB.60 Sample_3266.CB.391 5170−CB−10
Sample_3266.CB.94
Sample_3266.CB.87 Sample_3266.CB.62 Sample_3266.CB.283 Sample_3266.CB.395 Sample_3266.CB.20 Sample_3266.CB.98 Sample_3266.CB.390 Sample_3266.CB.387 Sample_3266.CB.57 Sample_3266.CB.55 5170−CB−6
5170−CB−3
Sample_3266.CB.267 5170−CB−16
Sample_3266.CB.206
5170−CB−4
Sample_3266.CB.96 Sample_3266.CB.272 Sample_3266.CB.106 5170−CB−14
5170−CB−22
Sample_3266.CB.64 Sample_3266.CB.396 Sample_3266.CB.298 Sample_3266.CB.303 Sample_3266.CB.302 Sample_3266.CB.21 Sample_3266.CB.4 Sample_3266.CB.19 Sample_3266.CB.300
-2
-1
0
NES
1
2
